# A neural circuit architecture for rapid behavioral flexibility in goal-directed navigation

**DOI:** 10.1101/2021.08.18.456004

**Authors:** Chuntao Dan, Brad K. Hulse, Ramya Kappagantula, Vivek Jayaraman, Ann M. Hermundstad

**Affiliations:** Janelia Research Campus, Howard Hughes Medical Institute, Ashburn, VA, USA

## Abstract

Anchoring goals to spatial representations enables flexible navigation in both animals and artificial agents. However, using this strategy can be challenging in novel environments, when both spatial and goal representations must be acquired quickly and simultaneously. Here, we propose a framework for how *Drosophila* use their internal representation of head direction to build a goal heading representation upon selective thermal reinforcement. We show that flies in a well-established operant visual learning paradigm use stochastically generated fixations and directed saccades to express heading preferences, and that compass neurons, which represent flies’ head direction, are required to modify these preferences based on reinforcement. We describe how flies’ ability to quickly map their surroundings and adapt their behavior to the rules of their environment may rest on a behavioral policy whose parameters are flexible but whose form and dependence on head direction and goal representations are genetically encoded in the modular structure of their circuits. Using a symmetric visual setting, which predictably alters the dynamics of the head direction system, enabled us to describe how interactions between the evolving representations of head direction and goal impact behavior. We show how a policy tethered to these two internal representations can facilitate rapid learning of new goal headings, drive more exploitative behavior about stronger goal headings, and ensure that separate learning processes involved in mapping the environment and forming goals within that environment remain consistent with one another. Many of the mechanisms we outline may be broadly relevant for rapidly adaptive behavior driven by internal representations.

## INTRODUCTION

Behavior often depends on the transformation of sensory information into motor commands based on an animal’s internal needs. Some direct responses to sensory stimuli do not require a brain [1] or even neurons [2], but neural networks enable animals to more precisely direct their actions, and to adapt their responses to sensory stimuli based on context, internal state, and experience [3–5]. However, sensory cues are not always reliable or even available, and many animals have evolved the ability to behave still more flexibly by generating and using internal representations of their relationship to their surroundings [6, 7]. These internal representations —for example, those carried by head direction (HD), grid, and place cells [8, 9]— are often tethered to sensory cues, but they allow animals to achieve behavioral goals without directly depending on those cues. Thus, goal-oriented behavior is often conceptualized as operating in two phases: first, a latent learning phase in which an animal builds internal representations of its spatial relationship to the environment, and second, a phase in which the animal uses good or bad experiences to learn representations of goals anchored to these spatial representations, and then modifies its behavior appropriately [10]. Many studies of learned behavior and its neural correlates, particularly those involving mammals, focus on the latter part of the second phase, using trained animals that have already learned the basic structure of tasks and environments; in doing so, they study task performance more than task acquisition (but see, for example, [11–15]). By contrast, in many natural settings, animals must develop spatial and goal representations simultaneously. Thus, any complexities in one evolving representation—for example, those induced in spatial representations by repetitive visual patterns in the surroundings [16–19]—impacts the other representations that are built upon it. Further, animals must use these still-evolving spatial and goal representations to select appropriate actions (or, in the parlance of reinforcement learning (RL) [20], to guide their behavioral policy), which, in turn, shapes how these representations develop over time. Finally, behavioral goals can themselves change over time based on environmental conditions and internal state, requiring animals to balance exploitation of a good situation with explorations away from it.

In this study, we delve into the dynamic process by which two learning systems, one unsupervised and the other reinforced, interact to enable flexible behavior (Fig 1a). Specifically, we study how internal representations of head direction (HD) and goals develop alongside one another—the former through unsupervised learning and the latter through reinforcement—and guide behavior in the context of rapid, visually-guided operant learning in the fly, *Drosophila melanogaster* [21]. We explore how this process might be implemented by identified circuits within an insect brain region called the central complex (CX) [22–29] (Fig 1b). We show that silencing so-called EPG or compass neurons (Fig 1b) that maintain flies’ HD representation [16] significantly impairs performance in a variant of a well-established operant learning paradigm for tethered flies [30]. The paradigm requires flies to modify their actions in response to heat punishment associated with one of two repeating visual patterns arranged symmetrically around the fly (Fig 1c, upper right panel), a simplification of the types of symmetries observed in natural scenes [31]. We used two-photon calcium imaging of compass neurons in tethered flying flies to demonstrate that this visual symmetry alters the dynamics of the fly’s HD representation in predictable ways, which allowed us to explore how these dynamics impact behavior. To characterize these behavioral changes, we quantitatively analyzed patterns of actions taken by individual tethered flies, used this analysis to isolate a set of control parameters that capture observed variability in the action selection process, and determined how these control parameters should optimally evolve over time and at different headings in order to support rapid learning. We show that flies’ behavior is consistent with an HD-dependent behavioral policy whose *form* is hardwired but whose *parameters* can be modified over time via a flexible goal heading. The resulting behavior depends only on the relative difference between the fly’s current and goal headings, and it reflects the symmetries of the visual scene through the underlying dynamics of the HD representation to which the policy is tethered. Importantly, rather than needing to learn the entire space of appropriate actions in a new setting, the hardwired form of the policy ensures that actions remain efficiently structured relative to the goal heading, even as that goal heading is changing over time. This, in turn, likely enables rapid learning and immediate adjustments to new goal headings. We use these results, together with existing physiological, behavioral, and anatomical observations [16, 29, 31–37], to construct a model for how CX circuits downstream of compass neurons might use the HD representation to both learn a goal heading in the fly’s visual environment and select actions that are driven by the fly’s current heading relative to its goal (note that we use the terms heading and HD interchangeably in our head-fixed setting). Finally, we used the model, together with the predictable changes in HD dynamics induced by the symmetry of the visual setting, to understand the interplay between generating HD and goal representations and selecting actions. We then suggest how this interplay shapes individual variability in learning and performance. Our observations provide a window into how rapid behavioral flexibility can be enabled by evolutionarily-conserved circuit motifs that provide strong inductive biases [38] for guiding behavior. These hardwired circuit motifs enable a modular implementation of an adaptive policy tethered to internal representations of an animal’s current and goal headings, and they underscore the importance of the reliability of and interaction between those representations in shaping the animal’s behavior.

**Figure 1:**
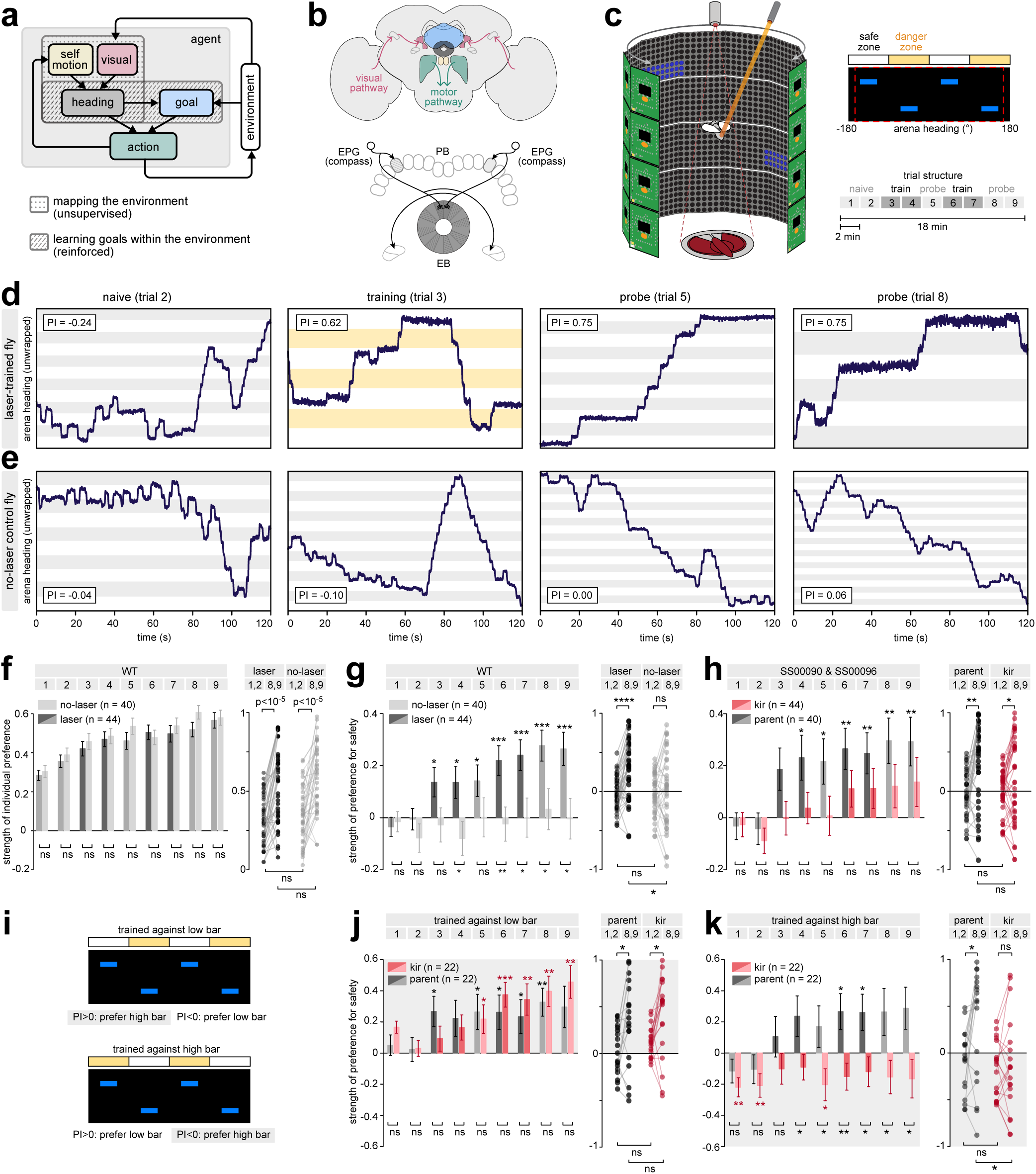
Flies need their neural compass to operantly learn to avoid heat punishment in a 1D virtual visual environment. **a)** This study explores how two distinct learning systems interact and influence one another to guide behavior. An unsupervised learning system combines self-motion and visual information to generate an internal representation of heading tethered to the environment. Downstream, a reinforced learning system uses this internal representation of heading to build and update an internal representation of a goal heading, which, in turn, modulates action selection based on heading. **b)** Upper: High-level schematic of the fly central complex (CX), highlighting visual pathways (pink) to the ellipsoid body (EB, gray) and motor pathways leaving the CX. Also shown are the protocerebral bridge (PB, yellow), fan-shaped body (FB, blue), and lateral accessory lobe (LAL, green). Lower: Schematic illustrating simplified representation of EPG (compass) neurons that maintain an internal representation of the fly’s head direction. Shown are two compass neurons, each innervating a single ‘wedge’ of the EB and a single ‘glomerulus’ of the protocerebral bridge (PB). The population of compass neurons tiles the entire EB and PB. **c)** Left: Schematic of flight simulator and LED arena used for behavioral experiments (see Methods). Upper right: The arena is divided into four quadrants; one set of opposing quadrants contains horizontal bars at a high elevation, and the other set of quadrants contains horizontal bars at a low elevation. During training trials, one set of quadrants (‘danger zone’) is paired with an aversive heat punishment (orange bars), while the other set of quadrants (‘safe zone’) remain unpunished (white bars). The red dashed box indicates the span of the visual arena (see Methods). Lower right: training protocol. During training trials, laser punishment is delivered whenever the fly’s arena heading falls within the danger zone. During naive and probe trials, no punishment is delivered. Each trial lasts 2 min. **d)** Example heading trajectories and performance indices (PI scores) from a single fly that underwent laser training. Trajectories were unwrapped to reveal the overall structure in the behavior (see Methods). Gray and yellow bars indicate arena headings within the danger zone that are punished (yellow bars; training trial), or that will be/have been punished (gray bars; naive/probe trials). Flies exhibit periods of straight flight in which they maintain a constant arena heading (fixations), punctuated by abrupt turns that lead to changes in arena heading (saccades). **e)** Same as **(d)**, for a no-laser control fly that did not undergo laser training. **f)** Average strength of individual preferences, without regard to safe and danger zones, for one genotype of flies (‘WT’), compared between groups that did (‘laser’) versus did not (‘no-laser’) undergo laser training. Behavioral preferences were measured by computing PI scores centered on flies’ trial-specific preferences (Methods); see SI Fig S2a for PI scores aligned to the final preference in trial 9. Left: average strength of individual preferences across trials. Error bars: mean +/- standard error. Significance between groups: two-sided Wilcoxon rank sum test against the null hypothesis that the strengths of individual preferences come from distributions with different medians. Right: changes in strength of individual preferences between early and final trials, shown for individual flies. Significance within groups: paired, two-sided Wilcoxon signed rank test against the null hypothesis that the difference in strengths of individual preferences between naive and probe trials has a median of 0. Significance between groups: two-sided Wilcoxon rank sum test against the null hypothesis that the strengths of individual preferences come from distributions with different medians. **g)** Learning performance for WT flies that did or did not undergo laser training (‘laser’ and ‘no-laser’, respectively). Left: average PI scores across trials. Error bars: mean +/- standard error. Right: changes in PI scores before and after laser training, shown for individual flies. Numbers in colored boxes at top indicate trials used to compute PI scores. Significance: same as in **(f)**. **h)** Same as **(g)**, but for two different genotypyes of flies (‘SS00090’ and ‘SS00096’) that underwent laser training. One group (‘Kir’) had Kir expressed in EPGs; a second group of genetically matched flies went through the same laser training but did not express Kir in EPGs (‘parent’). SI Fig S2b-f shows PI scores split out by genotype. See Methods for additional details regarding these double-blind experiments. **i)** Schematic illustrating training conditions in which the low versus high bar was punished (upper versus lower panels, respectively). **j-k)** Same as **(h)**, but split out into groups for which the low bar **(j)** versus the high bar **(k)** was punished. In both panels, red bars fall within the grey regions that indicate a high-bar preference; in contrast, gray bars are positive in both panels, indicating a preference for safety. \**p ≤* 0.05; \*\**p ≤* 0.01; \*\*\**p ≤* 0.001; \*\*\*\**p ≤* 0.0001.

## RESULTS

### Tethered flying flies change their visually-guided behavior after thermal conditioning

To study how flies adapt their behavior to changes in their surroundings, we modified a well-established visual learning paradigm [30]. In our modified paradigm, tethered flying flies in an LED arena were given closed-loop control of their angular orientation relative to a visual scene by locking angular rotations of visual patterns on the arena to differences in flies’ left and right wingbeat amplitude, a proxy for their intended yaw movements (Fig 1c, left panel) [39, 40] (see Methods). We used a periodic visual scene consisting of four quadrants of horizontal bars (Fig 1c, upper right panel). In two opposing quadrants, bars were positioned at a low elevation; in the other two quadrants, bars were positioned at a high elevation. We assessed flies’ naive preferences for different quadrants of the visual scene during a pair of 2-min-long ‘naive trials’ (Fig 1c, bottom right panel). During subsequent ‘training trials’, two symmetric quadrants (the ‘danger zone’) of the visual scene were paired with an aversive heat punishment delivered via an infrared laser to the abdomen of the fly; the remaining two quadrants (the ‘safe zone’) were left unpunished. In ‘probe trials’ with no heat punishment, we assessed whether flies formed lasting associations between different quadrants of the arena and the aversive heat.

Prior to training, flies explored different parts of the visual scene; the left columns of Fig 1d,e, show sample trajectories for two naive flies. This changed during training trials with laser punishment, when flies typically avoided spending time in the danger zones (Fig 1d, middle column). In contrast, ‘no-laser’ control flies that did not receive punishment continued to explore different parts of the visual scene (Fig 1e, middle column). This trend continued in the probe trials, when laser-trained flies continued to avoid the danger zones even after the punishment was removed (Fig 1d, right two columns), in contrast to control flies (Fig 1e, right two columns). Similar observations have been made in previous tethered fly visual learning studies in torque-meter-based flight arena setups [21, 30, 41] (see Methods for differences between these experiments and the setup we use here).

Previous studies have quantified fly behavior in this task using a performance index, or ‘PI score’ [42, 43], that measures the relative fraction of time spent in safe versus dangerous quadrants. Larger PI scores indicate a stronger preference for safety. Before we performed this analysis, we focused on a clear trend that we noticed across flies. When we ignored safe and danger zones and merely evaluated how strongly flies expressed their individual heading preferences in each trial, we found that both laser-trained and no-laser control flies expressed a preferred arena heading even in naive trials, and showed a strengthening of their preferred arena heading across trials (Fig 1f; SI Fig S1). This strengthening may be the result of flies improving the precision and control of their saccades and fixations in this new visual setting; in free flight, visual textures are known to influence the structure of flies’ behavior [44, 45]. This general feature of flies’ behavior in this paradigm is relevant to interpreting any analyses of PI scores. Notably though, while both laser-trained and no-laser control flies increased the strength of their preferred arena headings across trials, the laser-trained flies did so while also shifting their preferred arena headings significantly closer to the safe zone (Fig 1g, left column, dark bars), consistent with sample trajectories (Fig 1c,d) and with past results [32, 42, 43]. No-laser control flies did not significantly change their preference for either quadrant (Fig 1g, left column, light bars). These preferences were also reflected at the level of individual flies, with laser-trained flies showing significant shifts in their preferences (Fig 1g, right column, black points), and no-laser control flies showing no consistent shifts in preference (Fig 1g, right column, gray points).

Past studies have shown that perturbing various CX neuron types significantly impacts the ability of flies to perform this operant learning behavior [32]. Furthermore, compass neurons [33, 35, 46] and their inputs from the anterior visual pathway [47] are required for flies to display and maintain individualized heading preferences relative to a single visual landmark; this ‘menotaxis’ behavior is thought to aid dispersal and long-range navigation [48–53]. Additionally, inputs to the compass neurons in the EB have been linked to flies’ ability to remember specific orientations relative to a disappearing visual landmark [54, 55]. We therefore sought to test whether flies’ ability to flexibly modify their heading preferences in our more complex visual setting also depended on the compass neurons.

To explore whether compass neurons are required for operant visual learning behavior, we silenced compass neuron activity by selectively expressing the inwardly rectifying potassium channel, Kir2.1, in these neurons using two different split-GAL4 lines (Fig 1h-k; SI Fig S2-S3). These flies and flies from their parental control groups did not fly as well as wild type flies (data not shown), but displayed normal heading preferences in the single stripe environments that we exposed all flies to before beginning the main experiment (SI Fig S4; see Methods). In the operant learning task, flies from the control groups avoided the danger zone during training and showed good learning performance in the probe trials (Fig 1h, left panel, gray bars). In contrast, flies with silenced compass neurons showed consistently low PI scores (Fig 1h, left panel, red bars). However, these average trends were not entirely reflected in the behavioral preferences of individual flies; both parental control flies and compass-neuron-silenced flies showed significant shifts in their PI scores after training (Fig 1h, right panel).

Why might flies display consistent shifts in preference even when their compass neurons are silenced? Previous results from both walking and flying flies have shown that silencing compass neurons or visual inputs to the compass neurons exposes hardwired phototactic behaviors [33, 35, 46, 47]. These behaviors are likely controlled by more direct visuomotor pathways that are sensitive to the specific shapes and positions of visual stimuli [56]. In particular, flying flies are known to fly directly towards a sun-like stimulus when their compass neurons are silenced [33]. Given that flies increase their individual preferences across trials (Fig 1f), we asked if the high bar’s resemblance to this sun-like stimulus and a general increase in this innate preference across trials might account for our unexpected results. An innate preference for high bars would manifest in positive PI scores when trained against low bars (Fig 1i, top panel), and negative PI scores when trained against high bars (Fig 1i, bottom panel). This is indeed what we observed when we silenced the compass neurons: flies exhibited a consistent preference for the high bar, regardless of which pattern they were trained against (Fig 1j-k). In contrast, parental control flies showed consistent preferences for safety (Fig 1j-k). Thus, when compass neurons are silenced, it is likely that flies’ behavior in this paradigm is driven by visuomotor pathways that do not involve the CX. Taken together, these experiments suggest that wild type flies quickly learn to avoid dangerous parts of the visual scene in this paradigm, and that an intact HD representation is required for normal visual learning in this operant paradigm.

### Visual symmetries trigger jumps in flies’ internal HD representation

Having established that flies’ adaptive behavior in this paradigm depends on an intact HD representation, we next sought to understand the co-evolution of this representation and flies’ behavior. In what follows, we break this down into two separate learning processes, and then study their interactions: we first study changes in the HD representation as it tethers to a visual scene with symmetries, and model these through the lens of unsupervised learning (Fig 2); we then study changes in behavior in response to thermal conditioning (Fig 3), and model these through the lens of reinforcement learning (Fig 4); finally, we study how these two learning systems interact with one another to shape individual variability in performance (Figs 5-6).

**Figure 2:**
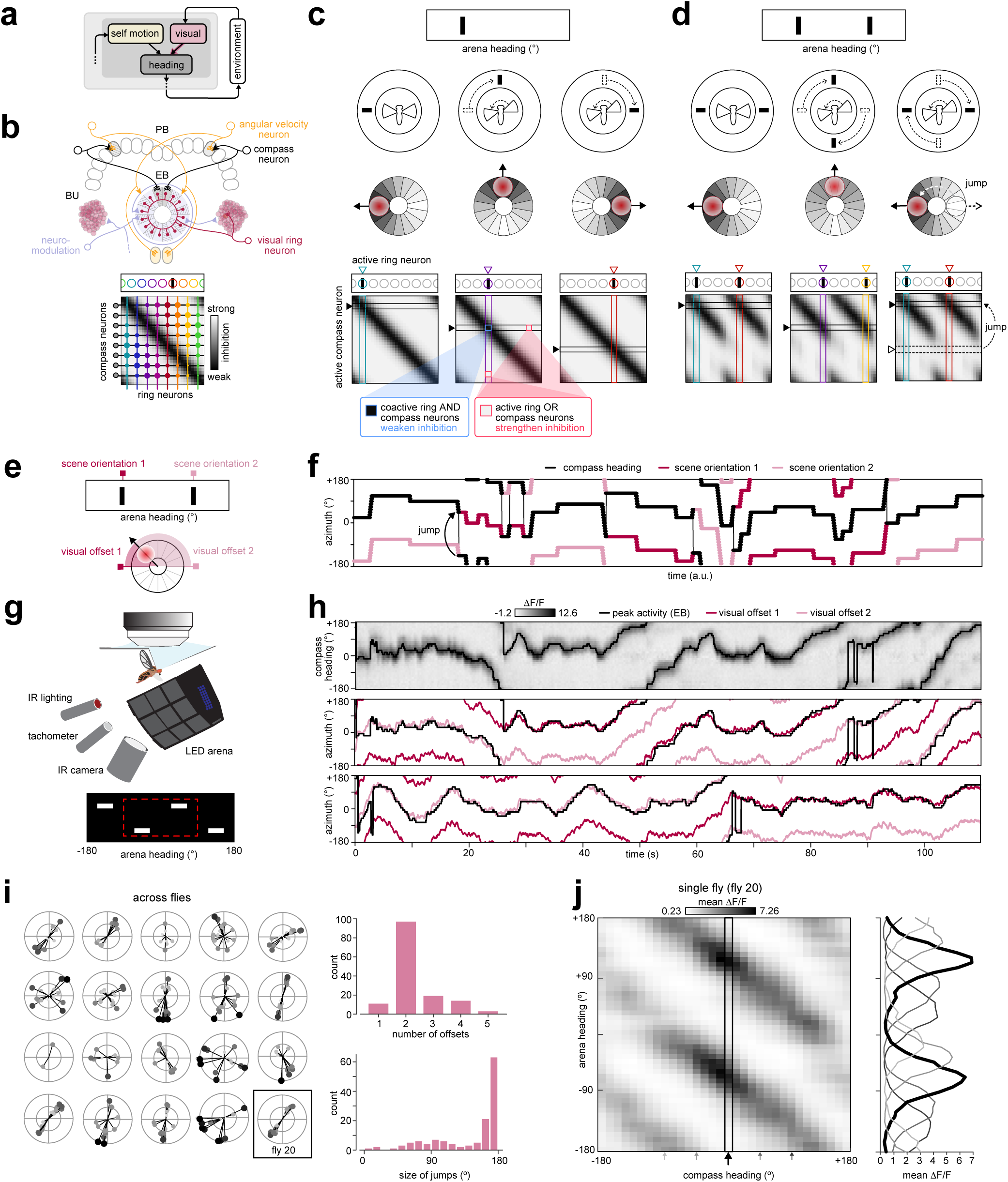
Mapping of a symmetric visual scene onto an internal heading representation triggers predictable jumps in compass neuron dynamics. **a)** Schematic of the unsupervised learning process that tethers the compass heading to the visual environment. **b)** Upper: visual ring neurons (red lines) make all-to-all inhibitory synapses (filled red circles in EB) onto compass neurons (black) that are idealized as only receiving input in the EB and sending outputs to the PB. Neuromodulatory neurons whose activity carries the fly’s motor state—in our model, the fly’s angular velocity [70]—project throughout the EB and BU. See *SI: Linking the Conceptual Model to Known Anatomy* for more details. Lower: Heatmap illustrates the strength of total ring neuron inhibition onto compass neurons; synaptic strength of individual ring neuron connections onto different compass neurons is shown by the radius of filled colored circles (different colors correspond to different ring neurons whose RFs are schematized above). Inhibitory Hebbian-like plasticity weakens synapses based on co-activity between ring and compass neurons during saccades [31, 70]. **c)** In an asymmetric scene with a single visual pattern (upper row), plasticity stabilizes a self-consistent mapping between ring and compass neurons (lower row, indicated by the diagonal structure in the weight matrix). This mapping ensures that both visual scene shifts and saccade-driven angular velocity inputs consistently and accurately update the heading representation (middle row shows bump in EB). Ring neurons that are activated by a particular view of the visual scene (colored circles that overlap with black bar in visual scene, and colored outline of columns in weight matrices) weakly inhibit compass neurons with the appropriate heading tuning (black outline of rows in weight matrices; note the dark color, and thus weak inhibition, from the active ring neuron onto the active compass neuron) and more strongly inhibit compass neurons with different heading tuning. When the fly saccades (different columns), this mapping ensures that the compass neuron activity bump remains tethered to movements of the scene. **d)** In a symmetric scene with two repeating visual patterns separated by 180° (upper row), plasticity stabilizes a two-fold symmetric mapping (lower row, denoted by repeating light and dark diagonal bands in the weight matrix) in which two arena headings opposite one another map onto the same internal compass heading. This mapping results in some compass neurons (black outline of row in weight matrix) being weakly inhibited (dark color in mapping) and others strongly inhibited (light color in mapping) by multiple ring neurons with 180°-opposite spatial receptive fields (colored outlines of multiple columns in weight matrices). Left column: A snapshot when the bump occupies an EB location in which the active compass neurons are weakly inhibited by the active ring neurons (see position of bump in EB schematic in middle row). Middle column: When the fly saccades, a different set of compass and ring neurons are activated (compare highlighted columns in first and second columns; note the shift in position of the visual stimuli). Right column: A further saccade results in the compass bump moving into an EB location in which the active compass neurons receive strong inhibitory synaptic input from the active ring neurons (black dashed outline of row in weight matrix, and black dashed outline of bump in EB). This would likely cause the bump to jump to a different location in the EB, where these same active ring neurons only weakly inhibit compass neurons (solid black outline of row in weight matrix, and red bump in EB). In a two-fold symmetric scene, this corresponds to a jump of 180°. **e)** We tracked the orientation of the compass heading relative to different reference orientations of the visual scene (pink square markers). For a scene with two repeating visual patterns, there are two ‘visual offsets’ (measured as the circular distance between the peak compass neuron activity and an arbitrary reference orientation of the visual scene) that correspond to visually-identical views of the scene. **f)** As a model fly navigates with respect to the scene shown in **(d)**, the compass heading (black trace) tracks one view of the scene (pink trace) for some time, and then jumps to the symmetric view of the scene (red trace). **g)** Schematic of two-photon calcium imaging setup for tethered flies flying within an LED arena. **h)** First row: Compass neuron calcium activity (heatmap) and location of peak activity (black line) during closed-loop tethered flight with a symmetric visual scene. Second and third rows: the peak activity (black) jumps between two different offsets (pink and maroon) that correspond to symmetric views of the scene. Shown for trials 1-2. **i)** Left: Polar plots showing offsets for all trials for each fly. Top right: Number of unique offsets, aggregated across flies and trials. Bottom right: Angular size of bump jumps, aggregated across flies and trials. A majority of jumps match the 180°-symmetry of the visual scene; see *SI: Linking the Conceptual Model to Known Anatomy* for discussion of the smaller peak at 90°. **j)** Main panel: Average Δ*F/F* of different wedges in the EB (compass heading) as a function of arena heading, averaged across all 9 trials for the fly shown in **(h)**. Right panel: Tuning curve for different EB wedges (curves corresponds to the wedges denoted by arrows in the main panel).

**Figure 3:**
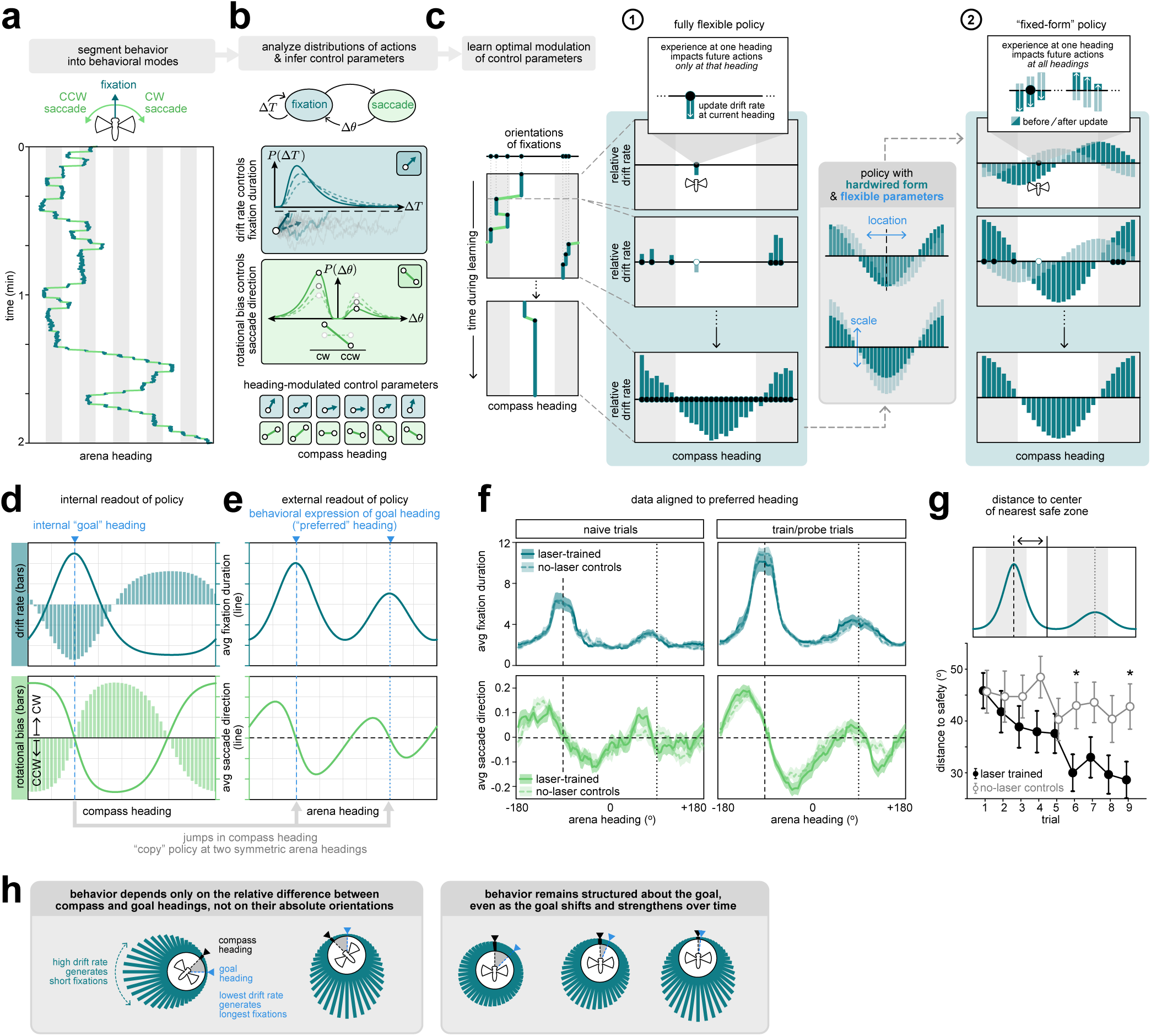
An inferred behavioral policy captures conditioned and unconditioned behavior. **a)** Left: same behavioral trace as shown in the left panel of Fig 1d, segmented into fixations (dark green) and saccades (light green). **b)** A behavioral policy generates alternating modes of fixations and saccades in model flies. Each mode is approximated by sampling a fixed angular velocity that is maintained over a duration of time. The distribution of fixation durations, *P* (Δ*T*), is well fit by a drift-diffusion process with an adaptive drift rate (dark green box); the distribution of turn sizes, *P* (Δ*θ*), is well fit by a lognormal distribution with an adaptive rotational bias (light green box; see Methods for more details). We hypothesize that the drift rate and rotational bias are flexibly modified as a function of the fly’s internal compass heading in order to control the average duration of fixations and the average direction of saccades. **c)** We schematize two different types of behavioral policies that could be used to modify the behavior of an agent (that is, a model fly) over time based on reinforcement. **(1)** With a fully flexible policy, the agent could store flexible associations between individual headings and the control parameters that govern behavior (illustrated here for the fixational drift rate). The agent would begin without any heading-dependent biases in its behavior. Experiences at one heading would then be used to alter the control parameters at that heading alone, and thus the agent must iteratively sample all headings in order to learn the correct settings for the drift rate. With sufficient training time, the agent would learn to modulate the drift rate in order to generate low drift (long fixations) in the safe zone, and high drift (short fixations) in the danger zone. See *RL Framework: Fully Flexible Policy* for details regarding this policy class. **(2)** The agent could instead use a behavioral policy whose form is fixed and resembles the long-time-limit outcome of scenario (1). Experiences at one heading would then be used to flexibly modify the location and scale of this policy, and thus an association made at one heading would impact future actions taken at all other headings. This significantly speeds up the learning process. See *RL Framework: Fixed-Form Policy* for details regarding this policy class. **d)** We train a reinforcement learning agent using a flexible policy (similar to that schematized in panel **(c-1)**) to optimize performance on this task by maintaining a single goal heading (see Methods). After training (dark curves), the agent learns to increase the average duration of fixations around the goal heading, and to bias saccades toward the goal heading. We hypothesize that flies use a policy of this fixed form that can be flexibly shifted and scaled based on experience (as schematized in scenario (2)). **e)** When the drift rate and turn bias are coupled to an unstable compass heading, the resulting behavioral readout is bimodal. This bimodality arises because the compass heading can jump between orientations in the EB that correspond to symmetric views of the visual scene, and thus leads to a ‘copying-over’ of the behavioral policy at symmetric arena headings. Dashed and dotted lines indicate the preferred and anti-preferred heading, respectively, separated by 180°. **f)** Average duration of fixations (upper panel) and direction of saccades (lower panels) generated by laser-trained (*n* = 44) and no-laser control flies (*n* = 40), measured before training (naive trials; left column) and after training (probe trials; right column). All data is aligned to the preferred arena headings of individual flies (vertical dashed line). Dark lines and shaded regions mark the mean +/- s.e.m. **g)** The preferred arena headings of laser-trained flies shift away from danger and toward safety (lower). See S6 for the duration of fixations and direction of saccades aligned to the safe/danger zones. Significance: two-sided Wilcoxon rank sum test (\**p ≤* 0.05) against the null hypothesis that quantities measured for laser-trained and no-laser control flies come from continuous distributions with the same medians. **h)** Minimal algorithmic requirements to tether a fixed-form behavioral policy to a flexible goal heading: the behavioral output must depend only on the difference between compass and goal headings (left box), and must preserve its form as the goal heading is shifted and scaled (right box).

**Figure 4:**
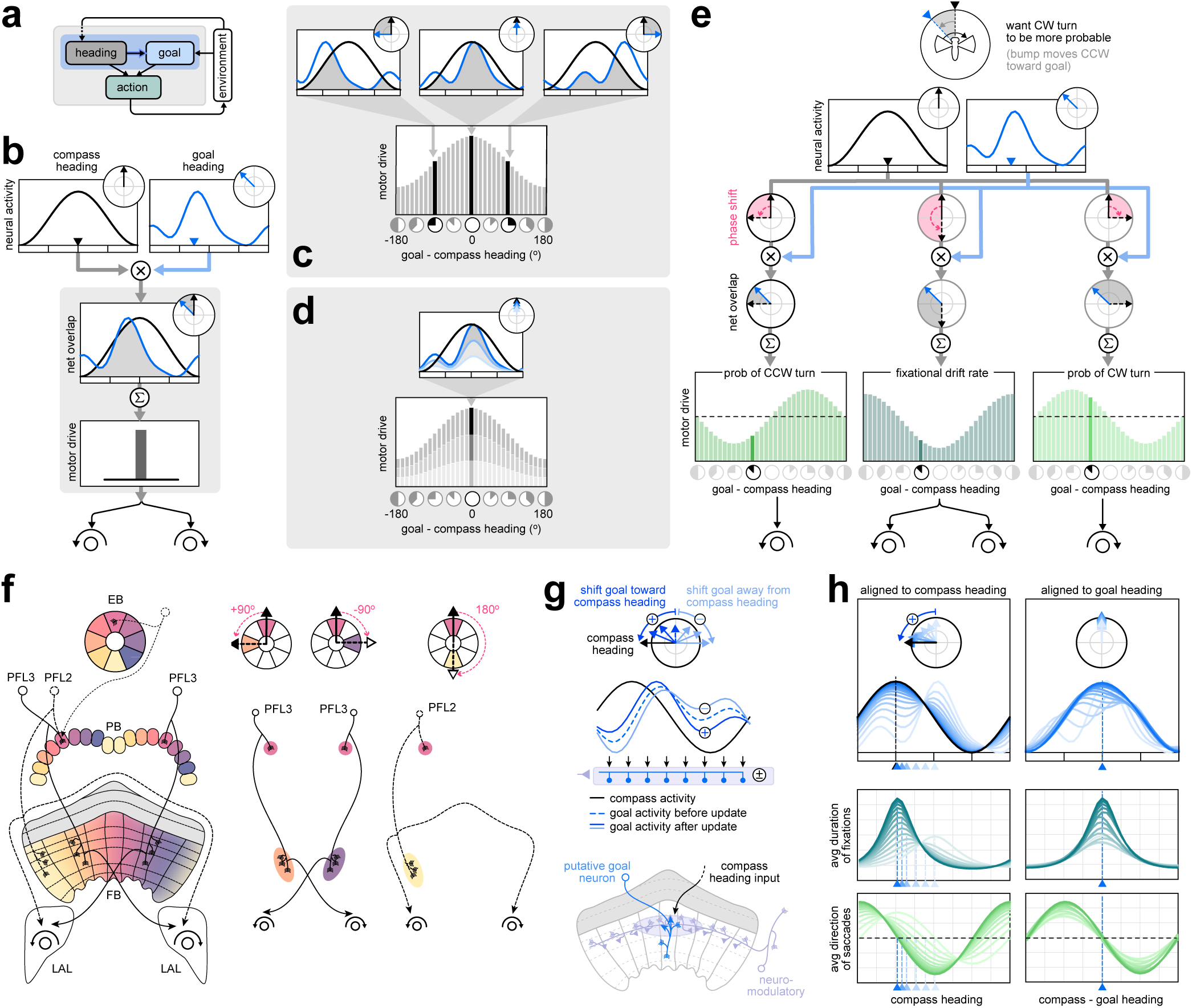
Hypothesized neural implementation of a fixed-form behavioral policy tethered to a flexible goal heading. **a)** Schematic of the reinforcement learning process that tethers action (via a fixed-form policy) to the difference between two internal representations—a compass heading and a flexible goal heading. **b)** To construct the fixed-form policy, we multiply a sinusoidal compass activity profile (black curve) and a goal activity profile of arbitrary shape (blue curve) to generate a motor drive (black bar) that controls a set of rotational controllers. The multiplicative operation is similar to computing the overlap between the two activity profiles (grey shaded region). Phasor representations above each activity profile illustrate the angular orientation, or phase, of the compass and goal headings, computed as the circular mean of each activity profile. **c)** For a given goal profile (blue curve), different orientations relative to a fixed compass heading (black curve) will generate different motor drives (grey filled bars in histogram). Sweeping across all orientations of the goal profile reveals a structured, sinusoidal motor drive that peaks when the compass heading is aligned with the goal heading and decreases as a function of the phase difference between the compass and goal heading, regardless of their absolute orientations. **d)** Weakening the goal profile reduces the motor drive across all compass headings. **e)** Shifting the compass heading by phase shifts of +90°, +180°, and *−*90° results in motor drives whose peak is shifted relative to the goal by the same amount. If these are used to drive downstream rotational controllers, we recover the overall architecture and fixed form of the policy schematized in Fig 3d,h. **f)** Left: The anatomical projection patterns of PFL ‘action neurons’ may implement the phase shifts and premotor projections schematized in **(c)**. Colors representing phases were propagated from EB wedges to PB glomeruli based on compass neuron projection patterns (see Fig 1b). They were then projected to columns of another CX region, the fan-shaped body (FB), based on the morphology of ‘zero-phase-shift’ PB-FB columnar neurons that project from PB glomeruli with approximately the same phase to an overlapping columnar region in the FB [29, 60, 80]. Note that phases shown here are a simplification of true phases; see [29] for details of phase assignments. Center: two populations of PFL3 neurons implement *−*90° and +90° phase shifts and project unilaterally to one of two premotor regions (the lateral accessory lobes, LALs) that drive either CW or CCW saccades. These phase shifts can be seen by noting how a neuron from one PFL3 population projects from the pink glomerulus in the PB to the orange columns in the FB, and then projects unilaterally to the LAL region that controls CW saccades. A neuron from the other PFL3 population projects from the pink glomerulus in the PB to the purple columns in the FB, and then unilaterally to the LAL region that controls CCW saccades. All left and right PFL3 neurons display phase shift motifs similar to the two sample neurons shown here. Together, the populations tile the FB and cover all angles. Right: The PFL2 neuron population implements a 180° phase shift between the PB and FB, and drives fixations by projecting bilaterally to both LALs. The phase shift can be seen by noting how one PFL2 neuron projects from the pink glomerulus in the PB to yellow columns in the FB. All PFL2 neurons display similar phase shift motifs and together tile the FB, covering all angles. **g)** We hypothesize that neuromodulatory neurons modify the goal heading based on the fly’s current compass heading. Positive reinforcement shifts the goal heading toward the compass heading, while negative reinforcement shifts the goal heading away from the compass heading. See S7 and *SI: Linking the Conceptual Model to Known Anatomy* for details about putative neurons that store and update the goal heading. **h)** Illustration of the evolution of the goal profile (blue curves, top row) when the model fly is at a fixed compass heading (black arrows/curves, first/second rows) and experiencing positive reinforcement. Over time (left column), the goal profile is strengthened at the current compass heading and weakened away from the current compass heading, leading to a shift in the goal heading. The behavioral readout (middle/lower rows) shifts with the goal heading. If we align different temporal snapshots to the goal heading (right column), we see that the behavioral readout is always aligned to the goal heading, even as the goal heading is shifting over time. As the goal heading becomes stronger, with a larger amplitude and more sinusoidal shape (darker blue curves), the behavioral readout also strengthens (darker green curves).

**Figure 5:**
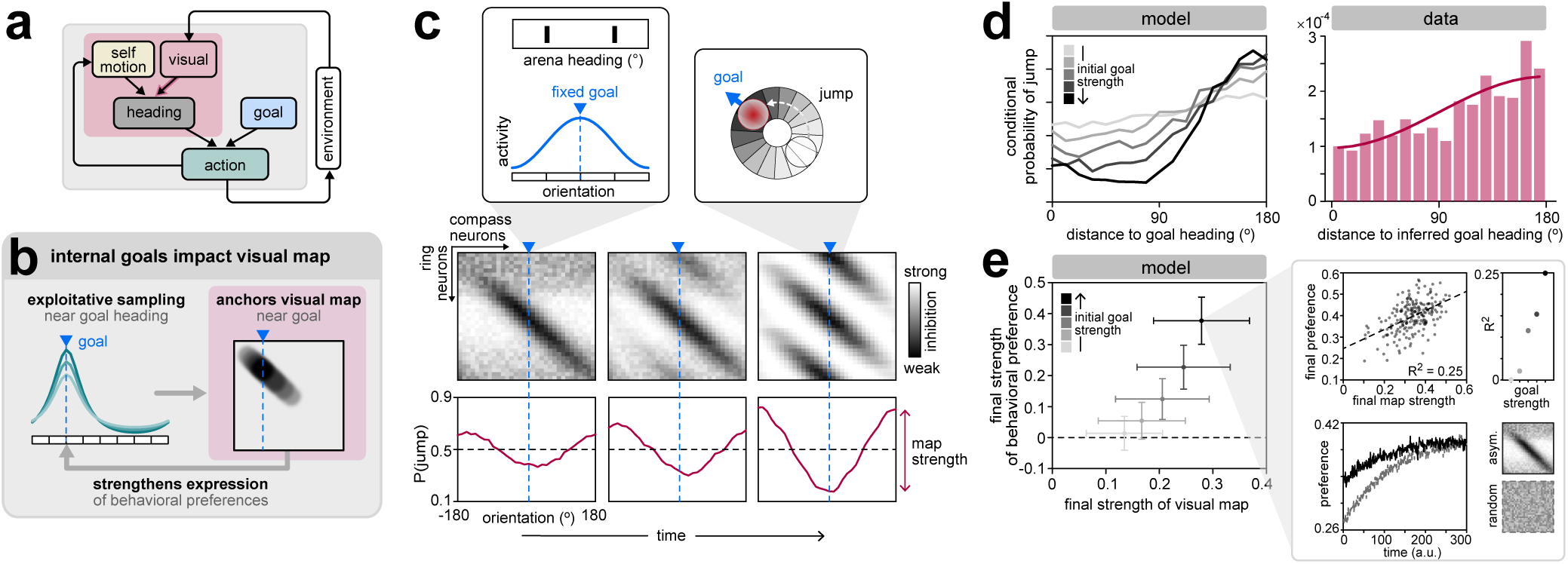
The location and strength of the internal goal shapes the evolution of the visual map. **a)** Schematic illustrating that the goal impacts the construction of a visual map through the actions selected by the behavioral policy. **b)** The behavioral policy leads to exploitative sampling around the goal heading, which impacts the evolution of the visual map and, in turn, the strength of behavioral preferences around the goal. **c)** Evolution of a visual map for a single model fly with a fixed goal heading. The model fly was initialized with a stable visual map derived from an asymmetric scene. Over time (successive columns), identical views of the symmetric visual scene induce plasticity in the same subsets of ring neurons, which leads to the development of symmetries in the mapping of the visual scene. The difference in net inhibition created by these symmetries shapes the probability that the bump will jump from different orientations (red curves). Over time, the most stable compass heading—at which there is the lowest probability of a bump jump—aligns with the goal heading. We use the circular mean of P(jump) as a measure of the strength of the visual map (Methods). **d)** Left: stronger initial goal headings (grayscale) lead to more pronounced differences in the conditional probability that the bump will jump when it is close to versus far from the goal heading. The conditional probability was estimated as the fraction of all visits to a compass heading that resulted in a jump. Right: Conditional probability of a bump jump as a function of angular distance from the preferred compass heading, aggregated across (real) flies. For each fly, the conditional probability was estimated as the fraction of all visits to a given EB wedge for which the bump jumped by more than 135° (accumulated across trials for which the bump maintained two different offsets relative to the visual scene). Bars: average probability across flies. Solid line: best-fitting cosine curve. **e)** Left: on average, stronger initial goal headings (grayscale) lead to stronger visual maps, which in turn lead to stronger behavioral preferences about the goal heading. Error bars: mean +/- s.e.m. Upper right: Within groups of individual model flies that began with the same initial goal strength, those model flies that developed stronger visual maps also exhibit stronger behavioral preferences. Lower right: Behavioral preferences increase over time as the visual map develops, and show more pronounced increases when model flies are initialized with a random visual map (dashed line) compared to a map initialized in an asymmetric visual scene (solid line). See SI Fig S8 for more analyses of simulations initialized with random maps. All simulations were initialized using a goal heading with a fixed orientation but a variable strength. Prior to learning, this goal heading was used to initialize a visual map in an asymmetric scene (with the exception of the dashed curve in the right panel of **(e)**, which began with a random map). Model results in panels of **(c-e)** are averaged over sets of 200 simulations that began with the same initial strength of the goal heading.

**Figure 6:**
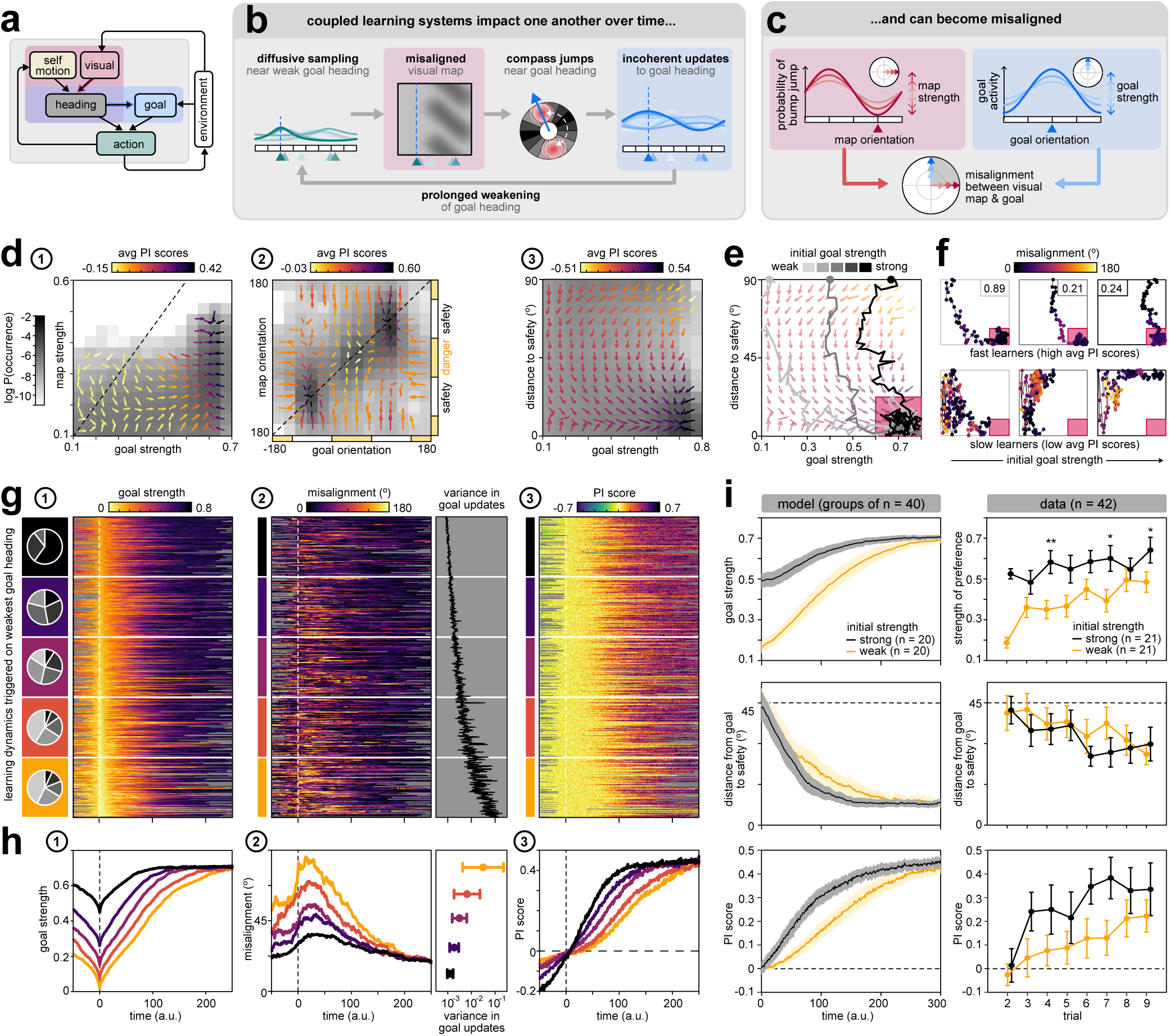
Interaction between visual scene mapping and goal learning introduces variability across individuals. **a)** Schematic illustrating that the two learning systems—one that maps the environment, and the other that learns goals within it—are coupled through the internal compass heading and the actions driven by the behavioral policy. **b)** A weak goal heading leads to diffusive behavior, which impacts the visual map and can cause the bump to become unstable at the goal heading. This, in turn, can lead to incoherent updates of the goal heading, for example if two 180°-opposite headings are successively reinforced or punished. These incoherent updates can then lead to prolonged periods of weak goal headings. **c)** Schematic illustrating features of the visual map and goal heading that we use to dissect the dynamics of learning in this paradigm. **d)** Vector maps illustrating the average effect of learning as a function of different properties of the visual map and goal heading. Individual vectors show how learning changes the strengths and orientations of the visual map and goal heading, depending on their initial strengths and orientations; longer arrows denote stronger average effects of learning (see Methods). Vector flows illustrate the progression of learning, which leads to higher average PI scores (darker arrows) and higher overall frequency of visits to that part of phase space (darker regions of heatmap). (1) Learning first increases the strength of the goal heading (yellow vectors pointing to the right), and then increases the strength of the visual map (purple vectors pointing upward). (2) Learning aligns the visual map with the goal heading (vertical yellow vectors pointing toward the diagonal), and then shifts the orientation of both vectors toward one of the two safe zones (purple vectors pointing inward toward the safe zones). (3) Learning increases the strength of the goal heading and shifts it toward safety (vectors pointing downward and to the right). **e)** Trajectories of individual model flies, superimposed over the vector map from panel **(d)-3**. Consistent with **(d)-1**, model flies eventually strengthen their goal headings and shift them to the center of the safe zone, reaching the lower right corner of the vector map (red shaded box). Darker gray trajectories indicate stronger initial goal headings. **f)** Individual model flies vary substantially in their trajectories through the same space. Individual columns highlight flies that began with increasingly strong goal vectors; top and bottom rows show model flies that had high versus low average PI scores, respectively. Boxed numbers in the top row indicate the fraction of total simulation time required for each fly to stably reach the lower right corner, highlighted in red; flies in the lower row did not reach the lower right corner during the simulation. Trajectories are colored by the degree of alignment between the most stable compass heading and the goal heading (see **(c)**); brighter colors indicate less alignment. Flies that were slow to learn (bottom row) suffered from prolonged periods of misalignment. **g)** Properties of learning trajectories over time, aligned by the time when the goal heading was the weakest (vertical dashed lines) and sorted by the value of the weakest goal heading. Colored rectangles indicate the groupings that were used to construct the averages shown in **(h)**; pie charts indicate the fraction of model flies within each group that began with different initial goal strengths. (1) Model flies that began with strong initial goal headings were more likely to maintain strong goal headings throughout training; flies that began with weaker initial goal headings tended to show further dips in strength subsequently. (2) Left: During periods where the goal heading is weak (bright regions in **(1)**), model flies tend to suffer periods of misalignment, where their visual maps become misaligned with their goal headings (see **(c)**. Right: This, in turn, leads to higher variance in the updates of the goal heading toward its final value (see SI Fig S9 for more details). (3) Periods of misalignment (bright regions in **(2)**) can slow the learning process, leading to prolonged periods of low PI scores. **h)** Same as **(g)**, but averaged over groups of flies that exhibited similar dynamics in their goal headings (colored groupings highlighted in panel **(g)-1**). Note that the top group (dark purple), which consists of flies that began with strong goal headings (primarily dark wedges in pie chart in **(g)-1**), shows only modest drops in their goal strengths over time, whereas the bottom group (orange), which consists of flies that began with weak goal headings (primarily light wedges in pie chart), shows large drops in their goal strengths over time (**(h)-1**). **i)** Left column: when model flies were partitioned based on the initial strength of their goal headings, model flies that began with weaker initial goal headings were (i) slower to reach strong goal headings (as expected; upper left), (ii) slower to shift their goal headings toward safety (middle left), and (iii) slower to increase their overall PI scores (lower left). Shaded regions: mean +/- std, computed over 100 sets of randomly-selected groups of 20 model flies whose initial goal strengths were above or below the median initial goal strength. Right column: when real laser-trained flies were partitioned based on the initial strength of their behavioral preferences (a proxy for the strength of their internal goal headings), flies that began with weaker initial preferences (*n* = 21) were (i) slower reach strong preferences (upper right), (ii) slower to shift their preferences toward safety, and (iii) slower to increase their PI scores compared to flies that began with stronger initial preferences (*n* = 21). Shown for WT flies that underwent laser training; see SI Fig S10 for data combined across three different genotypes of flies (WT, SS00090, and SS00096) and divided into two versus three groups. Significance: two-sided Wilcoxon rank sum test (\**p ≤* 0.05; \*\**p ≤* 0.01) against the null hypothesis that scores measured for flies with strong versus weak initial preferences come from continuous distributions with the same medians. Flies were separated into two groups based on whether their behavioral preferences were stronger or weaker than the median, as measured in trial 2. All simulations used in panels **(d-h)** were initialized using a goal heading with a fixed orientation centered within the danger zone, but with a variable strength (200 model flies were simulated for each strength). Prior to learning, this goal heading was used to initialize a visual map in an asymmetric scene (as in Fig 5). All simulations used in panel **(i)** were initialized analogously, but with variable initial goal orientations.

Although spatial representations make goal-directed behavior more robust to changes in sensory cues [3–5], these representations are themselves impacted by repeating patterns in the animal’s surroundings [16, 18, 19, 31]. In our setting, the symmetry of the visual scene gives us a unique opportunity to study how such an internal representation develops over time and impacts the learning of goal-directed behavior. To this end, we built a simple circuit model of the fly compass (Fig 2a). The fly’s compass heading is maintained as a single bump of activity that typically moves in concert with the fly’s rotations [16]. We assumed that the bump maintains a von Mises activity profile in the EB, and that its responsiveness to the fly’s rotations are accurately maintained by a ring attractor network and by neuron types that we did not explicitly model [29, 31, 47, 57–61]. We focused instead on the dynamics of the HD representation, which, in our model, are determined by a set of plastic synapses from visual ring neurons to compass neurons that evolve over time via an unsupervised learning process [31, 34, 62–64] (Fig 2b). Visual ring neurons have feature-tuned receptive fields that tile space [65–67], and they synapse onto compass neurons via all-to-all inhibitory connections in the EB [29] (Fig 2b, lower panel). During exploration of a visual scene, inhibitory Hebbian-like plasticity is thought to weaken synapses from active ring neurons onto active compass neurons at the location of the compass bump in the EB. We assume that the plasticity rule depends on angular velocity [31], a dependence that acts, at least in part, through angular-velocity-dependent dopaminergic neurons that innervate the EB but that we did not explicitly model (Fig 2b, top panel; [29, 68–70]). Over time, this plasticity creates a self-consistent mapping between the visual scene and the HD representation; in a simple scene with a single landmark, this self-consistency would be characterized by a diagonal band in the matrix of synaptic weights between ring and compass neurons (see heatmaps in the lower panels of Fig 2b,c; [31, 34]). Once stabilized, this mapping would enable the bump to move around the EB in perfect synchrony with the rotation of the visual scene, which would be captured by the activation of ring neurons with the appropriate spatial receptive fields (Fig 2c).

We next explored how this mapping would change in a symmetric scene. In a previous study, we showed that the mapping between a visual scene and the HD representation depends on particular characteristics of that scene; briefly, scenes whose rotational auto-correlation produce a single, dominant peak tend to induce one-to-one mappings from the visual scene onto the HD representation [31]. In contrast, the visual scene used in our behavioral experiments is two-fold symmetric, with two peaks in its rotational auto-correlation. When we simulated a fly moving through a simpler scene with the same two-fold symmetry, the mapping between ring and compass neurons developed multiple bands, such that ring neurons with 180°-opposite receptive fields had approximately equal synaptic weights onto the same compass neurons (heatmaps in Fig 2d). Since the scene is symmetric, these ring neurons will be identically active at two arena headings separated by 180°. However, the corresponding compass neurons—whose HD tuning is separated by 180°—will be inhibited to different degrees. The successive columns in Fig 2d play out the scenario that results as a fly with such a mapping turns through the symmetric scene. The compass bump begins at a location that is only weakly inhibited by active ring neurons. As the fly saccades to the opposite side of the visual scene (that is, to a view identical to its initial starting point), the compass bump moves to a location in which it is strongly inhibited by the same active ring neurons. This triggers a competition between two sets of compass neurons that are activated by the same ring neurons [16, 31, 34, 71], and the bump jumps across the EB to the location that is most weakly inhibited by the same active ring neurons (Fig 2d, right column).

Together, this highlights how the pattern of synaptic weights from ring neurons onto compass neurons influences the dynamics of the bump at different orientations within the EB. More precisely, it is the net inhibition from the population of active ring neurons that determines the tendency of the bump to jump between EB locations: the bump will preferentially occupy regions of the EB that are only weakly, rather than strongly, inhibited by active ring neurons (center row in Fig 2d). Note that such jumps are not restricted to visual scenes with precise symmetries [31]; if similar visual features are present at multiple locations in a scene, this would evoke similar ring neuron activation patterns at multiple arena headings, which in turn would trigger a competition between multiple compass neurons that are tuned to those headings. In the specific case of a two-fold symmetric visual scene, our circuit model assumes that the difference in net inhibition between two locations in the EB separated by 180° determines the probability that the bump will jump between these locations.

We next used our circuit model to track the compass bump over time (Fig 2e,f). After the mapping from ring to compass neurons had stabilized, the compass bump moved in concert with the fly’s saccades and remained tethered to changes in the orientation of the visual scene, but with occasional 180° jumps (Fig 2f). This was captured by the fraction of time that the bump maintained different angular relationships to the visual scene, a parameter that we and others refer to as the ‘visual offset’ [16, 31, 34, 35, 71] (Fig 2e). This offset is arbitrary, varies from fly to fly, and can vary over time as well [16, 31, 33–35]. In our circuit model, the bump jumped between two different visual offsets that were, by construction, separated by 180° and corresponded to identical views of the visual scene (Fig 2f).

We predicted that the HD representation in real flies would exhibit similar dynamics to those observed in our simulations, even though the scenes differ in the number of visual patterns that underlie their two-fold symmetry. To test this prediction, we used two-photon calcium imaging to monitor the HD representation in tethered flying flies in a visual setting similar to that used in the learning assay (Fig 2g; Methods). When we analyzed the consistency of the visual offset, we found that the compass bump tended to jump between two offsets that reflected symmetric views of the visual scene (Fig 2h). Examining offsets across flies, we found that the distribution of offsets was bimodal in a majority of flies (Fig 2i, left panel), with two peaks separated by 180° (Fig 2i, right panels). In correspondence with this, we found that different wedges of the EB were active for symmetric views of the visual scene, and thus their heading tuning curves had two peaks also separated by 180° (Fig 2j). This resulted in a two-to-one mapping from the visual scene onto the HD representation, similar to our simulations (Fig 2d) and to the tuning previously observed in simpler symmetric scenes [16, 31, 34, 71]. Thus, in the two-fold symmetric scenes used in our study and in previous experiments [32, 42, 43], the HD representation likely jumps between orientations that correspond to symmetric views of the scene. These jumps have important implications for behavior that is tethered to the HD representation, which we investigate next.

### A probabilistic policy captures tethered flies’ visually-guided behavior

Having established how the fly’s HD representation behaves in the visual setting of our paradigm, we next asked how this representation is used to guide learned changes in behavior. To answer this question, we sought to construct a generative model of behavior, or behavioral policy [20], that could account for naive and conditioned behavior. Our goal was to first characterize the variability in flies’ behavior in the absence of conditioning, then use this to inform a behavioral policy that could capture this variability, and finally predict how this variability should change based on experience.

Flies’ behavior can be quantified in terms of the execution of different modes of patterned movement [72, 73]. During free and tethered flight, flies exhibit periods of fixation during which they maintain a near-constant heading over time [39, 72, 73]. These fixations are often punctuated by body ‘saccades’, or ballistic turns, that result in abrupt changes in heading [72, 73] (Fig 3a). We observed these behavioral modes in both laser-trained and no-laser control flies (Fig 1d,e). We approximated behavior as being composed of only these two modes (Fig 3a,b), and then determined transitions between them (SI Fig S5a; Methods). Individual fixations tended to be long in duration with near-zero average angular velocity, while individual saccades were distinguished by high angular velocity over short durations (SI Fig S5b). We sought to use the variability in these properties across flies and trials to infer a generative model of behavior in which flies control the distribution of possible actions within each of these two modes. Specifically, our analysis supports a generative model in which flies control the relative probability of initiating clockwise versus counterclockwise saccades through an adaptive rotational bias, and control the average duration of fixations through a drift-diffusion process with an adaptive drift rate (Fig 3b, middle rows, SI Fig S5c-j). Thus, in this model, flies use control parameters to favor turns in one direction over another and to fixate more or less. We hypothesized that each of these control parameters could then be modulated by the fly’s internal compass heading in order to support learning (Fig 3b, bottom row).

How should these control parameters be modified over time based on reinforcement? Fig 3c schematizes two distinct learning strategies that would both lead to lower drift rates, and thus longer fixations, in the safe zone. One strategy would be to learn the value of different parameter settings at each compass heading, similar to a reinforcement learning (RL) algorithm called Q-learning [20]. In the simplest formulation of this algorithm, an agent (i.e., a model fly) builds and stores a policy that associates individual headings with individual action parameters, and these associations are independently updated based on reinforcement. The resulting policy can flexibly take on a variety of different functional forms. However, this flexibility comes at the cost of being very slow to learn, because the agent needs to iteratively sample all headings to learn a complete set of associations. In our setting, this type of policy converges to a profile of drift rates that is lowest—and thus generates the longest fixations—in the center of the safe zone (Fig 3c-1). Before and during learning, this policy would have no guarantee of selecting appropriate actions for headings that the agent had not directly sampled.

Alternatively, more rapid learning might be enabled by *fixing*, rather than *learning*, the functional form of the policy. The ideal form of such a policy might be expected to resemble the form to which a fully flexible policy converges after training (Fig 3c-1, bottom row). Learning could then act to shift and scale this function over time while preserving its overall shape. In contrast to a fully flexible policy, reinforcement experienced at one heading would automatically and immediately impact the selection of actions at all other headings. This, in turn, would significantly speed the learning process, but would limit the complexity of the associations that could be learned. The resulting behavior would have similar profiles of control parameters—and, therefore, similar distributions of selected actions—before, during, and after learning, but with experience-dependent shifts in the location and scale of this profile (Fig 3c-2).

Could a policy of fixed form but flexible location and scale account for flies’ behavior in our paradigm? Flies, like many other insects, display individual heading preferences via menotaxis-like behavior, maintaining a preferred heading for periods of time even when tethered (SI Fig S1) [33, 35, 47]. However, rather than purely fixating on one preferred heading, both walking and flying flies also explore other headings while centering their explorations around the preferred heading [33, 35, 47, 74–76], a behavioral pattern that matches our observations in this paradigm. Consistent with these studies, we reasoned that this behavioral preference was expressed in our paradigm via a fixed-form policy tethered to a flexible internal goal heading stored in the brain. As schematized in Fig 3c, we derived the idealized form of this policy by training model flies to use saccades and fixations to maintain a single goal heading (see *SI: Reinforcement learning framework*). After training, model flies learned to minimize the drift rate of fixations at the goal heading (Fig 3d, top row), and they learned to bias their saccades towards goal heading (Fig 3d, bottom row). The resulting behavior depends only on the relative difference between the compass heading and a single goal heading. However, in our symmetric visual setting, the fly’s compass bump jumps between the two 180°-symmetric views of the visual scene (Fig 2). When coupled to a fixed-form policy, this has the effect of “copying over” the form of policy at two symmetric arena headings (Fig 3e; Methods). Thus, if real flies in this symmetric visual setting rely on a fixed-form policy, we would expect the behavioral patterns of both naive and laser-trained flies to be bimodally structured. Indeed, when we aligned the fixations and saccades from individual flies to their preferred headings in the arena—a proxy for their internal goal headings—we observed bimodal behavior curves, with flies locally directing their saccades towards, and fixating longer at, the two orientations that correspond to symmetric views of the visual scene (Fig 3f). These behavioral patterns were consistent regardless of whether flies experienced laser training. However, as expected from model flies that shift their fixed-form policies upon reinforcement (Fig 3c-2), the preferred headings of laser-trained flies—but not of no-laser control flies—shifted toward the center of the safe zone (Fig 3g; SI Fig S6).

We note that observations of bimodal behavioral patterns have been made in a variety of different tethered flight arena studies of fly visual learning involving symmetric visual scenes [21, 30, 41]. In previous experiments, flies’ avoidance of the punished quadrants in probe trials has been interpreted as evidence for their ability to associate punishment with specific visual patterns [77]. Our observations from compass-neuron silencing experiments, behavioral analysis of saccades and fixations, and optimal RL models suggest instead that flies rely on a fixed-form behavioral policy that is tethered to flexible internal representations of compass and goal headings, and that these internal representations are shaped by the visual environment and by punishment. As a result, bimodal behavioral patterns likely arise from jumps in HD bump dynamics, rather than from actions tethered directly to repeating visual patterns. For this policy to retain its fixed form, circuits in the fly brain must ensure that: (1) the fly’s actions depend only on the relative difference between the compass and goal headings, and not on their absolute orientations; and (2) the behavioral structure about the goal heading remains intact even as the goal heading is changing over time (Fig 3h).

### A connectome-inspired circuit model can structure a goal-driven behavioral policy

How might the fly’s circuitry ensure that its behavior depends only on the difference between its compass and goal headings, and how might learning act to shift and scale this behavioral policy based on experience (Fig 3)? To study this, we constructed a second circuit model that combines two internal representations—a compass heading and a flexible goal heading—to implement a fixed-form behavioral policy (Fig 4a). In this model, the goal heading evolves over time via a reinforced learning process, and is compared to the compass heading to drive behavior. This model (Fig 4b) combines insights from existing models of the insect central complex [78–81] with conceptual insights from the fly CX connectome [29], but it intentionally abstracts away much of the known detail of CX circuit structure and function to focus on key computations that we believe underlie flies’ behavior in this assay. Below, we describe the different components of the model. In cases where CX neuron physiology and/or connectivity strongly suggest a role in such computations, we link these model components to potential CX neurons and circuit motifs (SI Fig S7). The *SI: Linking the Conceptual Model to Known Anatomy* provides a detailed description of specific CX neurons that might implement various aspects of the model.

In order to match our behavioral observations (Fig 3f), a key requirement of our circuit model is that it produce profiles of drift rates and turn biases that vary approximately sinusoidally as a function of the relative difference between the compass and goal headings (Fig 3d). We assume that these profiles are controlled by an underlying set of motor drives. To construct these motor drives, our model assumes that neurons in the protocerebral bridge (PB) shape the HD representation into a sinusoidal activity profile [29, 60] (Fig 4b), which is ideally suited for performing vector computations [82]. We did not impose this constraint on the goal representation, which we assumed is stored in a downstream CX structure called the fan-shaped body (FB) and can be modified by experience to take on arbitrary shapes. The model assumes that the fly’s motor drive is derived from the overlap between this arbitrarily-shaped goal activity profile and the sinusoidally-shaped compass activity profile. This can be computed via a dot product by multiplying the two activity profiles and summing the output (Fig 4b). When repeated for all possible orientations of the compass heading, the resulting profile of motor drives is itself sinusoidal, regardless of the specific profile of goal activity. Mathematically, this corresponds to taking the convolution between the heading and goal profiles (Fig 4c). Under this operation, the circular mean of the goal activity profile specifies the goal heading at which the motor drive profile peaks (see Eq. 31 in *SI: Reinforcement learning framework* for a brief mathematical explanation). Strengthening or weakening the goal activity changes the amplitude of this profile, but does not alter its shape (Fig 4d). Overall, this scheme ensures that the profile of motor drives is sinusoidal and peaks at the goal heading, regardless of its absolute orientation.

How might such a sinusoidal motor drive be appropriately phase-shifted to optimally control the drift rate of fixations and the turn bias in saccades? Our circuit model assumes that these shifts are achieved by ‘action neurons’ that shift the compass heading profile itself, prior to combining the compass and goal profiles to construct the motor drive (Fig 4e). Shifts by 90° in either direction produce the largest output —and are thus suitable to drive CW or CCW saccades— when the fly’s compass heading is 90° to the right or the left of the goal heading, respectively. Similarly, a shift by 180° has the largest output —and is thus suitable to drive short fixations with a high drift rate— when the fly’s compass and goal headings are anti-aligned. Together, this architecture ensures that the fly’s behavior remains appropriately structured with respect to the angular difference between the compass and goal headings, regardless of their absolute orientations.

Putative CX output neuron types that phase-shift the compass heading, and thus seem ideally suited to serve the role of the action neurons, have been described by us and others [29, 80, 83], and these hypothesized roles have found support in recent physiological data [84, 85]. Phase shifts in the CX are determined based on propagating a potential activity bump from the compass neurons to their downstream partners in the PB and the FB [29, 60]. Neurons that receive input in the PB from similarly-tuned compass neurons and project to similar columnar regions of the FB are considered to have a 0° phase shift. These neurons then serve as a reference frame for analyzing projection patterns of neurons that implement non-zero phase shifts. Based on such projection patterns, together with recent physiological recordings [84, 85], the most likely candidates for the action neurons that control saccades are the left and right PFL3 neuron populations. Each population inherits a copy of the heading bump from the compass neurons and from other neurons in the PB. However, we assume that these neurons participate in goal-related computations in the FB, where they project with *−*90° and +90° phase shifts relative to their PB inputs (Fig 4f, middle column). Further, both the left and right PFL3 neuron types project to a premotor region called the LAL, where they are well-positioned to drive downstream controllers for CW or CCW rotations [29, 36, 80, 83]. A third population of PFL2 neurons receives a copy of the current heading bump that is phase-shifted by 180° [29] in the FB and projects bilaterally to both controllers, and thus could serve the role of the action neurons that control fixations (Fig 4f, right column; [29]; see also [83, 85]). The PFL2 and PFL3 populations mutually inhibit each other to prevent saccades during a fixation, and vice versa. In summary, our anatomically-inspired model assumes that heading and goal activity is combined in PFL2 and PFL3 neurons whose net output is used to first determine whether to fixate or saccade, and, next, to control the duration of fixations or the direction of saccades depending on the fly’s current heading relative to its goal heading.

Anatomical and physiological data from the CX provided clear candidates for the action neurons that might integrate heading and goal information to mediate fixations and saccades. By contrast, a far greater number of neuron types with columnar projections in the FB could carry the goal activity profile that is combined with the compass activity profile in the three PFL neuron populations [29]. Here, we hypothesize that the goal heading is stored in synapses from a tangential neuron, which innervates a layer of the FB, onto putative columnar goal neurons. Tangential neuromodulatory neurons could then update the strength of these synapses via bi-directional Hebbian plasticity that is modulated by the fly’s current compass heading (Fig 4g), an idea with anatomical support from the connectome [29]. Negative reinforcement would weaken synapses at the current compass heading and thereby push the goal heading away from the compass heading, while positive reinforcement would strengthen synapses at the current compass heading and pull the goal heading towards the compass heading (Fig 4h). However, the duration of fixations and directionality of saccades would remain structured about the goal heading, even as it evolves over time (Fig 4h, left column). As the goal heading strengthens, the fly’s behavioral expression of its internal goal would also grow stronger, with longer fixation durations near the goal heading and more biased turns towards it (Fig 4h, right column). This would naturally lead to more exploratory, diffusive behavior when the goal heading is weak, and more exploitative, goal-directed behavior when the goal heading is strong. In the *SI: Linking the Conceptual Model to Known Anatomy*, we provide a detailed discussion of candidate neuron types that could serve to store and update the goal heading.

In summary, we propose that tangential and columnar neurons in the FB store and modify the goal heading in their synapses through heading-dependent reinforcement from tangential neuromodulatory neurons. The strengths of these synapses are read out by other columnar goal neurons before being transmitted to PFL neurons, which combine this goal input with the phase-shifted compass heading in a manner that enables flies to maintain structured behaviors around internal goals even as they change. Note that this desired outcome significantly constrains the set of putative circuit implementations: to ensure that the policy remains efficiently structured about the goal heading, the goal neurons must provide a consistent readout of the goal heading independent of the compass heading; however, to update the goal heading based on reinforcement at a particular heading, the goal synapses must be modulated by a signal that depends on the compass heading. Our proposed model is one simple circuit instantiation that simultaneously meets these criteria and is supported by known anatomy (see *SI: Linking the Conceptual Model to Known Anatomy* for more details).

### Goal strength and location influence the mapping of visual scenes onto the fly’s compass

In previous sections, we showed how the symmetric visual scene of our paradigm is mapped onto an internal compass heading in the EB (Fig 2), how circuits in the PB and FB might implement a fixed-form behavioral policy tethered to the difference between the compass and goal headings (Fig 4e-g), and how reinforcement signals in the FB might modify goal headings and thereby shift and scale the policy (Fig 4g,h). Next, we sought to explore how actions selected by this policy in turn impact the mapping of the visual scene onto the compass in the EB (Fig 5a). To this end, we combined our two circuit models (Figs 2 and 4) to study how a fixed goal heading impacts the mapping of visual scenes onto the compass.

As described earlier (Fig 2), the visual mapping from ring neurons onto compass neurons is influenced by repeating patterns in the visual scene: in our two-fold symmetric setting, this mapping develops two bands, such that ring neurons with 180°-opposite receptive fields have approximately equal synaptic weights onto the same compass neurons. Importantly, when this mapping is driven by behavior that is tethered to an internal goal heading, the patterns of synaptic weights develop at specific locations relative to that goal (Fig 5b). Specifically, because most of the fly’s saccades are in the neighborhood of the goal heading, the compass headings around the goal heading are the first to develop weakened synapses from ring neurons activated by the visual scene at those headings; compass neurons with 180°-symmetric tuning receive stronger inhibition from the same active ring neurons (Fig 5c). This net difference in inhibition will cause the HD bump to favor regions of the EB near the goal heading; whenever the bump is away from these regions, it will tend jump back toward the goal, further weakening the synapses from active ring neurons near the goal heading. Over time, the probability that the bump will jump from any given heading —determined by difference in net inhibition between locations in the EB separated by 180°—will develop into a sinusoidal profile whose minimum is aligned with the goal heading. As a result, the bump will be least likely to jump when the fly’s compass heading is aligned with the goal (Fig 5c). The stronger the goal heading, the more pronounced this effect will be (Fig 5d, left panel).

We next examined experimental data to ask how frequently the compass bump tended to jump from different locations in the EB, relative to the number of visits the bump made to that location. We analyzed the location of these jumps relative to an inferred goal heading, which we defined as the location of maximal residency in the EB that corresponded to the fly’s preferred arena heading measured from behavior (see Methods). We found that jumps were least likely to occur at this inferred goal heading, and most likely to occur 180° away from this inferred goal heading (Fig 5d, right panel). Thus, symmetries of the visual scene induce jumps in the compass bump [16, 31, 34, 71]; in a scene with two-fold symmetry, these jumps are not uniform around the EB, but are most likely to occur at the location symmetric to the goal heading in the EB (Fig 5d, right panel), as predicted (Fig 5d, left panel).

Finally, we asked whether the strength of the visual map impacts the behavioral expression of the internal goal heading. When we analyzed the behavioral preferences of model flies, we found that stronger goals led to stronger visual maps, which in turn led to stronger behavioral preferences (Fig 5e, left panel). The correlation between map strength and behavioral preference was maintained even for the same goal strength (Fig 5e, upper right panel). These preferences strengthened over time as the visual map was stabilizing, regardless of whether the model fly began with a random map or one that had previously stabilized in an asymmetric scene (Fig 5e, lower right panel; SI Fig S8). This effect may contribute to the non-specific strengthening of PI scores we observe in both no-laser control flies and laser-trained flies across trials (Fig 1f).

Taken together, these results describe how the exploitative sampling of headings near an internal goal affects the orientation and strength of the visual map, and how this, in turn, impacts the expression of a behavioral preference. In the next section, we allow the goal heading to change over time based on experience, and we explore how the interaction between mapping the visual scene and modifying the goal heading impacts behavioral performance.

### Deciphering fly-to-fly variability in operant visual learning

Up until this point, we have considered the separate learning processes of modifying an HD representation and modifying a goal representation. However, in this operant visual learning paradigm, the fly must form these two representations simultaneously (Fig 6a). These two representations evolve over time via separate, but coupled, learning processes: the mapping of the visual scene onto compass neurons evolves via an unsupervised learning process modulated by the fly’s saccades through the visual scene, while the goal heading evolves via a reinforced learning process guided by positive and negative experiences at different compass headings. These learning processes are coupled through the compass heading, which directly updates the goal heading, and through the behavioral policy, which is tethered to both the compass and goal headings and which determines the visual and thermal feedback via the selected actions. How is a fly’s behavior influenced by the interactions between these two learning processes, and how do the two *evolving*internal representations —of compass heading and goal heading— influence each other and ultimately the fly’s actions (Fig 6b)? To answer these questions, we used our combined circuit models to explore the coupled dynamics involved in learning compass and goal headings. We simulated a learning task that mimicked what real flies experienced, with a two-fold symmetric scene coupled to negative and positive reinforcement in danger and safe zones, respectively. We then tracked the orientation and strength of both the visual map and the goal heading over the course of learning (Fig 6c). Prior to training, each model fly began with a goal heading that was used to form a visual map in an asymmetric scene; for simplicity, we assumed that all model flies began with goal headings centered within the danger zone but of variable strengths. The structure of flies’ exploration of the arena and of different compass headings is at first dictated by their initial visual map and goal heading. Over time, the goal heading strengthens (Fig 6d-1), which in turn structures the flies’ actions and helps to strengthen the visual map (Fig 5). Eventually, both the goal heading and the visual map lock onto one of the two safe zones (Fig 6d-2). Different flies undergo different experiences because of the probabilistic nature of the structured behavioral policy, and because of the probabilistic nature of the bump jumps. This impacts the evolution of both the visual map and the goal heading, as schematized in Fig 6b and characterized below. However, on average, model flies increase the strength of their goal headings and shift their goal heading towards safety, which results in an increase in their PI scores (Fig 6d-3).

We next dissected individual variability in learning. While some model flies exhibited learning dynamics that were representative of the ensemble average and quickly converged to the safe zone (Fig 6e; Fig 6f, top row), other model flies were much slower to learn (Fig 6f, bottom row). We hypothesized that this slower learning might arise from bump jumps that were not coherently aligned with the evolving goal heading, such that jumps would frequently move the bump away from, rather than toward, the goal heading. To test this, we measured the evolving alignment between the most stable compass heading and the goal heading (Fig 6c). We found that a consistently good alignment between the two headings tended to allow flies to more rapidly converge onto strong goal headings in the safe zone (Fig 6f, color map in top row). In contrast, prolonged periods of misalignment tended to slow learning, with model flies often getting ‘stuck’ in bad parameter regimes and requiring more time to ‘escape’ to regimes that allowed them to reliably learn safe goal headings (Fig 6f, color map in bottom row). Although we observed this pattern in flies with weak and strong initial goal headings, prolonged periods of misalignment were more likely to occur in model flies that initially began with weak goal headings. To dissect the relationship between the strength of the goal heading and its alignment with the visual map, we aligned the behavior of all model flies to the timepoint of their weakest goal heading (Fig 6g-1; averages of each category in Fig 6h-1). Following a drop in the strength of the goal heading, we observed that model flies tended to suffer from periods of misalignment; the weaker the goal heading, the longer and more pronounced was the period of misalignment (Fig 6g-2, left panel). This led to jumps in the HD representation away from the goal heading, which in turn led to higher variance in the updates of the goal heading toward its final value (Fig 6g-2, right panel; SI Fig S9). These incoherent updates adversely impacted performance by slowing the evolution of PI scores (Fig 6g-3). Although we observed these trends across all model flies regardless of the initial strength of the goal heading, prolonged periods of poor performance were more likely to unfold in model flies that began with weak goal headings (pie charts in Fig 6g-1).

We next sought to compare these model predictions to the performance of laser-trained flies in our behavioral paradigm. We repeated the simulations shown in Fig 6d-h, this time varying both the strength and location of the initial goal heading and comparing behavior between groups that began with weak versus strong initial goal headings (Fig 6i; SI Fig S10). When we separated real flies into two groups based on the initial strength of their preferred arena heading—a proxy for the initial strength of their goal headings—they showed similar behavior to model flies in the evolution of their goal strengths (Fig 6i, top row). Real flies that began with weaker inferred goal headings tended to maintain significantly weaker goal headings across trials, consistent with our model predictions. We would expect that the two trajectories in real flies would eventually converge like those of model flies if it were possible to perform training for longer durations, an experimental limitation of this tethered flight behavioral paradigm. The evolution of the goal heading toward the center of the safe zone showed similar trends between model and laser-trained flies: flies with weaker initial goal headings were slower to shift their goal headings toward the center of the safe zone relative to flies with stronger initial goal headings. Finally, when we compared PI scores between the two groups of flies, we once again found similar trends: both model and laser-trained flies with weaker initial goal headings are slower to increase their PI scores (Fig 6i, bottom row). In summary, flies’ individual actions determine how the learning process evolves, but starting with a strong goal heading, even if inaccurate, may structure behavior in such a way that it enables a faster remapping of the visual environment onto the compass, a faster determination of good headings within the environment, and—as a consequence of more stable visual mappings aligned with stronger goal headings—a faster and more accurate updating of internal goals.

## DISCUSSION

We sought to explore how flexible behavior is shaped by co-evolving internal representations of compass and goal headings during early experience in a novel environment. We first established that the ability of *Drosophila* to perform an operant visual learning task [21, 30] depends on an internal HD representation carried by their compass neurons, and used calcium imaging to reveal jumps in the population dynamics of these neurons in the symmetric visual setting of the paradigm. These dynamics gave us a window into how the consistency of internal representations impacts operant learning, an issue we explored with an anatomically-inspired circuit model of CX computations underlying this flexible behavior. In contrast to the distributed representations that are typically learned in large, recurrent neural network models (Fig 7a) [86], our proposed architecture is modular and highly structured to optimally implement key computations, with specific cell types carrying, modifying, and combining distinct internal representations to guide behavior (Fig 7b). We did not aim to include all the available details of CX circuitry, instead incorporating only those aspects of CX anatomy and physiology that we believe to be essential to understand circuit function in our task. Based on our behavioral and physiological results, as well as previous physiological, behavioral, connectomic, and modeling studies focused on the CX [16, 29, 31, 33–36, 57, 62, 71, 78–81, 83], we suggest that flies rely on a fixed-form behavioral policy that specifies controllable properties of the fly’s actions relative to the angular difference between two internal representations: one of the fly’s current heading, and the other of a single flexible goal heading. Our abstract model enabled us to examine how individual flies use their HD representation to construct this goal representation based on arena-heading-specific heat reinforcement. However, because the HD representation is learned alongside the goal heading, any inconsistencies in the HD representation impact the formation of the goal representation and, through the policy, the fly’s behavior. The fly’s behavior, in turn, impacts how the animal samples its new environment and, therefore, how it updates both representations. Our framework resolves the conundrum of how a fly with a single internal goal heading increases its residence in two safe zones at opposite headings. More broadly, using an artificial visual environment with repeating patterns tethered to aversive punishment helped us to reveal this interdependence of representation learning and goal-directed behavior. We show how this interdependence plays out in the performance of model flies and how it could explain some of the variability we observe in the performance of real flies as well.

**Figure 7:**
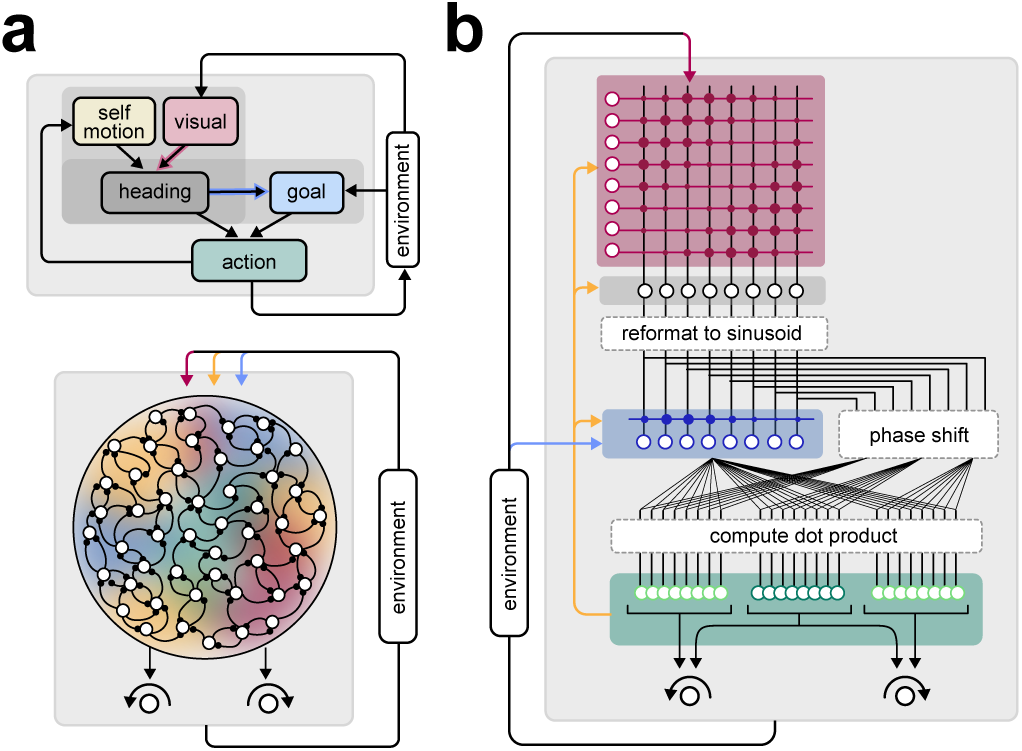
A modular circuit architecture for enforcing the form of a behavioral policy coupled to multiple evolving internal representations. **a)** Recurrent neural network models typically comprise a large ensemble of identical units (circles) connected via plastic synapses. When trained on a task with multiple different inputs (colors), these networks tend to generate highly distributed representations in which individual units exhibit mixed selectivity for several inputs. **b)** In contrast to the diffuse plasticity and distributed representations in **a**, our proposed circuit model exhibits a highly modular structure in which distinct cell types carry and combine internal representations of distinct variables. A set of inductive biases (dotted boxes) enforce the form of a behavioral policy, which in turn restricts plasticity to a set of synapses onto goal neurons (blue shaded box). Another set of synapses onto compass neurons is also plastic (red shaded box).

Several recent models have relied on PFL3 phase shifts, or asymmetries in synapse counts from EPG to PFL neurons in the PB, to move model flies to goal headings [36, 80, 81, 83–85]. In contrast to some of these models (for example, [81]), our model does not rely on the HD representation to maintain a consistent relationship to the visual scene; rather, it allows the HD and goal representations to co-evolve, a key aspect of early learning in a novel environment. Additionally, in most of these models, walking insects generate individual turns of exactly the correct size to reach goal headings, with any deviations from such sizes considered noise. While this may be appropriate in the context of homing behavior [80, 81], our model assumes that the brain can dynamically adjust the degree of exploitative behavior about the goal heading through an intrinsically probabilistic policy, and that CX neurons control parameters of this distribution. We find that this approach better captures the finer-grained aspects of flies’ behavior in our experiments, and specifically how the distributions of saccades and fixations depend on heading. Recent studies carried out in parallel to ours have highlighted the role of PFL3 neurons [84, 85] and PFL2 neurons [85] in walking flies performing menotaxis [84, 85]. Although it is not yet known how these neuron types function during learned behavior in flight, the experimental results in these studies are largely consistent with the model we propose. The potential mechanisms we propose for how putative goal neurons might operate, and for how goal information might be stored, updated, and read out, do not yet have strong support in available physiological and anatomical data. However, there is evidence that an FB columnar neuron type called hΔc, which carries an odor-gated wind-direction signal, may serve as a goal heading during olfactory navigation [83]. More recently, experiments involving FC2A neurons, whose connectivity and morphology [29] make them well-suited to carry a goal signal to the PFLs [87, 88], have shown that they may indeed play such a role [84]. Additional experiments will be needed to ascertain whether these or other FB columnar neurons show changes in synaptic weights and activity profiles required for them to perform the computations that we hypothesize in our model.

Our compass neuron silencing and imaging results suggest that many studies in visual pattern, spatial, and color learning in tethered flying flies [30, 32, 43, 89] should be reinterpreted. In all these studies, flies were believed to have learned to associate their actions with specific visual features or objects; these conclusions were based on the near-symmetric structure of fly behavior in settings with the same patterns in opposing quadrants of circular arenas. This symmetry of the visual environment was believed to rule out the possibility of learning based on heading. We suggest instead that flies in these studies build HD representations tethered to different visual surroundings, and rely on HD-representation-based goal learning to associate rewards or punishments with different compass headings. We further argue that the observed bimodality in flies’ behavioral responses to visual patterns arises from the downstream consequences of symmetry-driven jumps in the HD representation. These dynamics render irrelevant any limitations that a single-goal-heading-based policy might otherwise impose on the fly’s actions in a setting with multiple safe and dangerous headings associated with repeating visual patterns. It is possible that such jumps would be less frequent in free flight, where proprioceptive cues are likely to play a greater role in controlling compass bump dynamics (see, for example, [71]). Notably, neurons encoding the symmetries of an animal’s environment are not limited to tethered insects: neurons in the retrosplenial cortex of freely behaving rodents also display such responses [18, 19], and bidirectional HD tuning in that system has also been modeled as relying on Hebbian plasticity [90]. Although the symmetric environments used here may seem artificial, natural scenes can have repeating visual features, and these can also induce variability in the relationship between the compass bump and the visual scene [31]. The specific impact of such variability on the performance of particular tasks depends on how reliant downstream circuits are on compass-heading-dependent policies. We used model simulations to show how jumps in flies’ internal representations could explain some of the variability we observed in the performance of real flies. However, it is not known whether flies can learn multiple distinct goal headings, nor to what extent they can form more complex policies beyond the single-goal-heading policy invoked here. Performance in a place-learning paradigm suggests that flies can learn more complex associations to guide navigation through 2D environments [91]. It is also possible that, in contrast to the CX-based learning we study here, many spatial navigation behaviors may rely on associations made in the mushroom body [25, 92–98], a brain region that has also been suggested to be involved in some operant visual learning tasks in tethered flying flies [43, 99].

For most of our analyses, we decomposed the tethered fly’s behavior into two different modes: fixations and saccades. Freely flying flies are known to exhibit these different modes [100] that are characterized by distinct kinematic properties and necessitate both continuous and discrete control [73, 101]. The same modes have also been observed previously in tethered flying flies [73]. We explicitly incorporated these modes into the construction of a behavioral policy, and we used the observed variability in kinematic parameters across flies and trials to infer the parametric form and control parameters of this policy. When combined with optimal RL algorithms [20], this enabled us to specify how these control parameters should change based on experience. This approach bears similarities to recent studies proposing that learning operates on generative parameters that control distributions of movements, rather than on the higher-dimensional space of all possible movements [102–104]. We then used this approach to specify how these control parameters should be structured as a function of the fly’s compass heading to maintain a goal heading over time. Rather than learning this structured relationship from scratch, we showed that this relationship is pre-built into how untrained flies sample their surroundings, a strategy that might facilitate dispersal in the absence of explicit goals [74]. Indeed, these same patterns of structured behavior resemble those observed in tethered behaving flies responding to visual features that are innately attractive or aversive [101]; note that an additional parameter, the size of saccades, also varies based on angular distance from a ‘goal’ object in those visual environments, something that was less striking our visual setting (data not shown). We suggest that flies may, in fact, rely on different visuomotor pathways in different settings. Responses to innately attractive or aversive objects [56, 105–108] could rely on direct and hardwired visuomotor pathways that recruit banks of feature detectors in the optic foci [109, 110] and, perhaps, relatively stereotyped motor responses dependent on the spatial receptive fields of feature detector inputs [101]. These pathways may indeed underlie the behavior of compass-neuron-silenced flies that favor high bar stimuli (Fig 1i-k). In contrast, these silencing experiments also suggested that learned responses rely on an indirect pathway that recruits a probabilistic behavioral policy whose fixed form depends only on the relationship between internal representations of compass and goal headings. We did not explore whether these probabilistic biases could be strengthened through longer training protocols. We note that although it might seem optimal to steer towards and then maintain a single goal heading rather than initiate directed saccades that are probabilistically biased toward this heading, using such a default behavioral strategy would likely be too predictable to avoid predation [111] and would minimize exploration. Indeed, many animals, including flies, show stochasticity in their actions when behaving freely in dynamic settings [111, 112]. As discussed above, connectomic [29] and physiological evidence [84, 85] suggest that the architecture of columnar neurons in the FB could enforce the form of a behavioral policy for steering towards a goal heading by using the fly’s compass heading [29, 80, 83] (see also *SI: Linking the Conceptual Model to Known Anatomy*). Importantly, within this model, learning acts to modify the location and strength of the goal heading while preserving the entire structured relationship between different control parameters across different headings. Our results suggest that these associations are mediated via a flexible pathway through the CX; however, direct sensorimotor pathways that instruct reflexive actions might, in fact, work in concert with these flexible pathways, and in the aversive conditioning setting considered here, might enable the fly to quickly escape punished zones. How such pathways are balanced to guide reflexive and flexible actions, and whether the outcome of reflexive actions can be used to inform future flexible actions, is not yet known (but see *SI: Linking the Conceptual Model to Known Anatomy* and *SI: Reinforcement learning framework* for a discussion of how reflexive actions might be incorporated in the framework presented here).

In contrast to many behavioral paradigms in mammals, flies in this paradigm learn within a matter of minutes —without shaping or instruction— to direct their behavior away from punishment and towards safety. Our results suggest that flies’ rapid learning relies on strong inductive biases that enforce the form of behavioral policy and thereby dictate flies’ sampling strategy. This fixed-form policy assumes the existence of a single goal heading for the fly, and efficiently directs the fly towards identifying and orienting towards such a heading, wherever that happens to be. During learning, the fly can then use experience at one heading to effectively update actions at all headings, rather than having to learn associations at each heading individually. This type of inductive bias reduces the possible scenarios that are explored during learning and can thereby speed up the learning process when these scenarios are compatible with the learning task [113, 114]. Recent RL studies have explored how such inductive biases might be constructed by learning common features across different learning tasks [115, 116], a process known as learning to learn [117]. Here, we show how inductive biases that are likely learned over evolutionary timescales can be inferred directly from an animal’s behavior in the absence of an explicit task. The ability to rapidly exploit these inductive biases, in this case by shifting and strengthening a single goal heading, relies on faster-timescale learning. Because these inductive biases guide how flies sample new compass headings, the resulting behavioral structure can help anchor and rapidly update the mapping of the visual world onto an internal representation of compass heading, which can in turn speed the learning of the goal heading derived from the compass heading. The dynamic evolution of these interacting internal representations, expressed through the inductive biases that enforce the form of the behavioral policy, dynamically alters the degree of exploitative behavior with the weakening and strengthening of the goal heading. Whether, and to what extent, this faster-timescale learning could modify the form of the behavioral policy—for example, by further increasing the exploitative component through increased training—remains unknown. This could further be combined with behavioral state information—for example, about walking versus flight—to switch the specific actions that are controlled through the same behavioral policy.

It has been suggested that rapid learning in both artificial and biological systems relies on combining context-dependent memories with efficient exploitation of environmental and task structure [114]. Indeed, inductive biases that exploit such structure may hold the key to animal learning [38]. Here, we provide insights into how specific neural circuits might instantiate a behavioral policy that has evolved to address ecological needs through efficient actions, and how this policy both informs and is shaped by flexible, and perhaps context-dependent, internal representations of the environment. Targeted genetic access to the specific cell types that might mediate this learning provides an avenue for rigorously testing these ideas in the near future.

## ACKNOWLEDGEMENTS

We thank Jason Wittenbach and Daniel Barabasi for pilot modeling efforts, Parvez Ahammad for useful early discussions, and Alice Wang for pilot experiments. Eyal Gruntman, Michael Reiser, Josh Dudman, John Tuthill, TJ Florence, Sung Soo Kim, Yoshi Aso, and members of the Jayaraman and Hermundstad labs provided insightful input at different points of the study. We received useful feedback on the manuscript from Sandro Romani, Marcella Noorman, Hannah Haberkern, and Dan Turner-Evans. We are grateful to Bjorn Brembs for informative email discussions on the design and interpretation of well-established visual learning studies, and to Stephen Thornquist and Gaby Maimon for helpful discussions on the statistical analysis of behavioral data in an earlier version of the manuscript. We thank Gudrun Ihrke for supporting these experiments through her expert management of Project Technical Resources (PTR). We thank Dan Milkie (now at Janelia) and Andy Chiu from Coleman Technologies for help with developing the FPGA Wingbeat Analyzer. We thank Janelia Experimental Technology (jET) for technical assistance, especially: Jinyang Liu for the LED arena controller code, Steve Sawtelle for the D2A converter connected to the FPGA Wingbeat Analyzer, and Tanya Tabachnik, Igor Negrashov, and Bill Biddle for designing and manufacturing the fly mounting assembly used for two-photon imaging. We are grateful to Janelia’s Drosophila Resources team for stock building and maintenance, and to the Media Prep Facility for special fly food that kept our finicky flies flourishing.

This work was funded by the Howard Hughes Medical Institute.

## AUTHOR CONTRIBUTIONS

CD performed all experiments, data processing and initial data analysis, except for Kir-silencing experiments and associated control experiments, which were performed by RK under CD’s supervision. AMH performed all behavioral analyses, with input from CD, VJ, and BKH. VJ and AMH analyzed imaging data with input from CD. All authors interpreted results. AMH developed the theoretical framework and performed all modeling and simulations, with conceptual input from VJ, BKH, and CD. The proposed CX circuit implementation of the model was conceived over multiple feedback loops involving VJ, AMH, BKH, and CD, with BKH contributing, in particular, to the circuit implementation of the behavioral policy. AMH and VJ wrote the paper, with input and editing from CD and BKH.

## METHODS

### Experimental Methods

#### Fly culture

Parental flies were grown sparsely on Wurzburg food in bottles for at least 6 generations [118]. Crosses were first done in vials then transferred to bottles after 1-3 days, followed by transferring to a new bottle every day to limit F1 density. 10 males and 25 virgins were used for each cross. The day after eclosion, F1s were transferred to a new bottle with a piece of kimwipe for self-cleaning and transferred again to a new bottle with kimwipe the day before imaging or behavioral experiments. All experiments were performed with 5-6 days old female flies.

The crosses for the flies used in the experiments are: WT: 11D03AD males x WTB virgins (Figs 1, 3); EPG silencing: SS males x WTB;;UAS-Kir2.1-EGFP virgins (Fig 1); Parental controls: SS males x WTB virgins (Fig 1); EPG two-photon imaging: 60D05 males x 20xUAS-GCaMP6f [su(Hw) attP5; attP2, VK00005] (WTB) (Fig 2). We used the following SS flies: SS00090 [119] and SS00096 [57].

#### Visual arena

A blue LED circular arena [40] was assembled with 44 panels (4 rows and 11 columns, spanning 120° in elevation and 330° in azimuth), with the LED emission peaking at 464 nm (Bright LED Electronics Corp., BM-10B88MD). Two layers of blue filter (Roscolux #59) were laid on top of the LED panels to allow 0.04% transmission. Each fly was tethered at the end of a tungsten wire and positioned in the center of the arena. An 880 nm LED (Digi-Key, PDI-E803-ND) illuminated the fly from above. A custom-built wingbeat analyzer (University of Chicago Electronics Shop) measured the wingbeat frequency and amplitudes for both wings. Yaw turning was computed as the left minus right wingbeat amplitude. A computer (Dell, R5500) controlled the timing of the experiments through a data acquisition device (National Instruments, USB-6229 BNC) and sampled the flight parameters at 1 kHz. This is in contrast to most well-established visual learning studies, which have relied on torque meters to measure the fly’s instantaneous torque and drive the rotation of a paper drum imprinted with visual patterns [30, 42]. In imaging experiments, visual patterns were displayed on a set of cylindrically arranged blue LED panels (464 nm) extending *±*90° in azimuth and *±*45° in elevation, and tilted by 26° from horizon towards the fly. Two layers of blue filter (ROSCO #59 INDIGO), an electromagnetic shield and a diffuser were used to cover the LED panels as previously described. The rest of the LED display was covered with black aluminum foil. Only half of the 360° pattern was displayed at any given time because the LED display only spanned *±*90° in azimuth. A custom camera-based FPGA wingbeat analyzer was used to measure the fly’s intended yaw turning instead of the diode-based wingbeat analyzer.

#### Flight visual learning

All flies first went through a 30 s test trial in a closed-loop environment with a single vertical blue stripe (15°w x 120°h) to assess basic flight performance before operant learning experiments in the horizontal bar environment (SI Fig S4).

For operant learning experiments, the 360° yaw space around the fly was divided into 4 quadrants. A single horizontal blue bar (37.5°w x 11.25°h) was displayed in each quadrant, with alternating elevations at +/- 30°, such that the pattern repeats every 180°. Throughout the assay, the fly had closed-loop control of the visual pattern it was flying towards. The unconditioned stimulus (US) as punishment was a fiber-coupled infrared laser (Edmund Optics, 975 nm, 400 mW) modulated by a 10 kHz square wave with varying duty cycles output from a function generator (Agilent, 33220A, 20 MHz) and gated by the specific positions of the arena pattern such that either the higher bar quadrant or the lower bar quadrant was accompanied by the laser punishment aimed at the back of the fly. The laser was turned off during the pre-training näıve trials and post-training memory/probe trials. The visual pattern was jump-rotated randomly to a new position after every trial. A 100 ms air puff towards the fly was triggered whenever the fly stopped flying. However, only data during flight from flies that flew continuously without any stops or without any airpuffs for more than 60 s in all 3 trial types were included for further analysis. All visual stimulation and behavior parameters were recorded with a data acquisition device (National Instruments, USB-6229 BNC). During no-laser mock experiments, the US laser was not turned on. The EPG silencing and parental control experiments were performed double-blind by RK.

#### Fly preparation for imaging during flight

Flies were transferred to a polypropylene tube using a custom 3D-printed funnel positioned on the top of the opened bottle, then anaesthetized in a custom brass cold plate at 4°C. The largest female was selected to fit onto a custom aluminum mounting bridge cooled to 4°C, and held down with vacuum suction ventrally. The bridge was then rotated to hold the fly upside down and an inverted custom laser-milled PEEK holder pushed up the fly’s head from below, with a center hole lined up under the head. Small drops of UV-activated epoxy were used to glue the fly head, thorax and the back of the head capsule to the holder. Another small drop was used to glue the proboscis. The eyes were kept completely below the holder to allow unhindered visual stimulation and most of the back head plate was exposed through the center hole in the holder. The legs were left intact and the wings kept free to flap during flight because of the inverted pyramid shape of the holder. The back plane of the head was angled at approximately 26° to match the angle of the visual arena under the two-photon microscope. For imaging experiments, we used an LED arena with 18 panels (3 rows and 6 columns, spanning 90° in elevation and 180° in azimuth). For flight experiments, only flies that could fly continuously for 90 s while maintaining closed-loop stripe fixation after mounting were selected. Artificial hemolymph as described previously [120] was used to fill the holder reservoir from the top. A window was carefully opened on the back head capsule with a tungsten dissection probe and fine forceps and the trachea underneath were gently picked away to allow optic access to the brain [121].

#### Two-photon calcium imaging

Calcium imaging was performed on a two-photon microscope (Bruker Nano, formerly Prairie Technologies). A Chameleon Vision II or Discovery laser (Coherent) tuned to 920 nm was used with the power adjusted to the lowest sufficient level, usually between 3 and 20 mW at the sample. A resonant galvanometer mirror was used to scan the laser beam along the x-axis at 8 kHz, resulting in a frame rate of 60 Hz with 256 by 256 resolution. For volume imaging, a piezo motor drove the 40x objective (Olympus, LUMPlanFl/IR, NA 0.8) along the z-axis. The 2-plane z-stack acquisition was repeated over time throughout the trial at a rate of 14.5 volumes/s. The green and red channel signals, when applicable, were collected through a set of dichroic mirror (575 nm) and band-pass filters (525 + 35 nm for green, 607 + 22.5 nm for red). A GaAsP photomultiplier tube (Hamamatsu, 7422PA-40) was used to acquire data from each channel. Each imaging series was triggered from the experiment-controlling computer.

### Behavioral Analysis

All data analysis was performed in MATLAB (MathWorks Inc., Natick, MA).

#### Partitioning behavior into fixations and saccades

All data analyses were performed after segmenting behavioral traces into fixations and saccades. We developed a custom algorithm to perform this segmentation (SI Fig S5a). We first filtered the difference in wingbeat amplitude between left and right wings, *A*^WB^, using a bandpass filter of order 6, with a lower cutoff frequency of 0.1 Hz and an upper cutoff frequency of 10 Hz (we will denote this filtered signal as 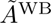). We then used sign changes in the filtered amplitude to segment the trajectory into a set of individual turns; each turn in this set was thus defined as a sequence of time points *{t}* for which 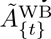 had a consistent sign.

A turn that produces a sustained nonzero difference in wingbeat amplitude 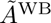 will lead to changes in the arena heading *x* in the opposite direction. We used this to define a quantity 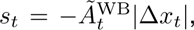 where Δ*x_t_* = *x_t_*_+1_ *− x_t_* is the instantaneous change in arena heading (allowing for wrapping between pixel 96 and pixel 1). This quantity measures the coherence between differences in wingbeat amplitude and changes in arena heading; *s_t_* will be large in magnitude during times when changes in wingbeat amplitude lead to large and coherent changes in arena heading, and will be zero when changes in wingbeat amplitude do not lead to a change in arena heading. We thus used this signal to select turns that fall into the former category, where *s* is large in magnitude.

To this end, we first selected candidate saccades as those turns that led to a total change in arena heading of at least 2 pixels (7.5°). For this subset of turns, we used *s_t_* to refine the beginning and end of individual turns. We defined the beginning of the turn as the timepoint *t*_start_ for which there was the largest instantaneous change Δ*s_t_* = *s_t_*_+1_ *− s_t_*, and the end of the turn as the first timepoint thereafter for which *s_t_* dropped below 1*/*4 of its maximum value (i.e., 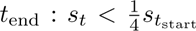). The remaining timepoints (*t < t*_start_, and *t > t*_end_) were segmented as separate turns. We repeated this process until all large turns had been refined in this way.

This resulted in a refined set of turns; these turns included both the candidate saccades that led to a change in arena heading, and the small turns that did not lead to a change in arena heading. We removed all turns during which either (i) the wingbeat frequency *f* ^WB^ dropped below a threshold of 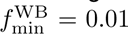 (for the upright arena) and 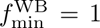 (for the two-photon arena) for any amount of time, or (ii) the turn intersected with a period of time that spanned 500 ms before and 500 ms after an airpuff. We then ranked each remaining turn according to a quantity *r*(turn) = *|(s_t_*_start:*t*end_)(*x*_*t*_end__ *− x*_*t*_start__)*|* that combines the average change in wingbeat amplitude *(s_t_*_start:*t*end_ ) with the total change in arena heading (*x_t_*_end_ *− x_t_*_start_). This quantity will be largest for turns that are large and fast, which comprise a small fraction of the entire set of turns. We thus used an outlier detection procedure to identify those turns for which *r* exceeded a threshold *r*_thresh_ = *Q*3 + exp(3*M*) *∗* 1.5 *∗* IQR. Here, *Q*3 is the third quartile (or 75%) of *r*, IQR is the inter-quartile range, and *M* is a skewness estimated using the med-couple of *r* [122]. *r*_thresh_ was estimated separately for individual flies and trials, based on the distributions of turns produced by the given fly in the given trial.

The set of turns for which *r > r*_thresh_ were classified as saccades. The periods of time between each saccade were events that we then further classified as either fixations or periods of drift. Before this classification, we first removed any periods of time that intersected with an airpuff event. We then identified events for which *f* ^WB^ dropped below 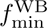 for more than 30ms; we removed the period of time that spanned 500 ms before the first drop in *f* ^WB^ and 500 ms after the last drop in *f* ^WB^. Each remaining portions of the event that exceeded 50 ms was then as a fixation if the variance in *x_t_* was below 36 pixels, and drift otherwise (if *x_t_* is Gaussian distributed with a standard deviation of *σ*, this cutoff ensures that 4*σ* of the distribution falls within 90°). Portions of the event below this 50 ms threshold were classified as “other” and were retained for analyses of PI scores and residencies.

In the main text, we focused our analysis on fixations and saccades. We described these two behavioral modes in terms of their duration and their angular velocity; here, the angular velocity (in deg*/*ms) is given by 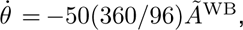 where the factor 50 converts between wingbeat amplitude and pixels*/*ms, and the factor (360*/*96) converts from pixels*/*ms to deg*/*ms. Saccades were characterized by short durations and large average angular velocities, while fixations were characterized by long durations and low average angular velocities (SI Fig S5b).

#### Characterizing fixation properties

Individual fixations varied substantially in their duration, and the distribution of these durations was heavy-tailed. We therefore considered three putative heavy-tailed distributions: log-normal, inverse Gaussian, and generalized Pareto. We fit each of these three distributions to the distribution of fixation durations *P* (Δ*t*) under two different conditions: when fixations were accumulated across flies within a given trial, and separately when fixations were accumulated across trials for a given fly. We performed this fitting for laser-trained and no-laser control flies.

Prior to fitting, we removed fixations whose durations were below a variable threshold Δ*t*_thresh_. We then evaluated fitting performance for 25 evenly spaced values of Δ*t*_thresh_ between 20 ms and 500 ms. We used the MATLAB function *fitdist.m* to perform the fitting, and we used the Bayesian information criterion (BIC) to evaluate fits. We found that the inverse Gaussian distribution, IG(Δ*t*; *µ, λ*), was the best-fitting distribution across a majority of scenarios (trials or flies) for thresholds between 100 ms and 300 ms; within this range, a threshold of 200 ms maximized this number of scenarios for which the inverse Gaussian was the best fit. We therefore performed the remainder of our analysis on fixations whose duration exceeded Δ*t*_thresh_ = 200 ms.

The inverse Gaussian distribution is characterized by two parameters: a mean *µ* and a shape parameter *λ*. This distribution can be generated by a drift diffusion to bound process with a mean drift rate *ν*, spread *η*^2^, and bound *a* (SI Fig S5g). This process yields an inverse Gaussian distribution *P* (Δ*t*) = IG(Δ*t*; *a/ν, a*^2^*/η*^2^) whose parameters *µ* = *a/ν* and *λ* = *a*^2^*/η*^2^ are defined in terms of the parameters of the diffusion process. When we compared the best-fitting values of *µ_F_* and *λ_F_* across different datasets (where the subscript *F* denotes that parameters were fit to the distribution of fixations), we found that the variability in these parameters was consistent with a drift diffusion process with a variable drift rate *ν_F_* but fixed spread 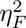 and bound *a_F_*. To illustrate this, note that the mean mean = *µ* and variance var = *µ*^3^*/λ* of the inverse Gaussian distribution satisfy log(var) = 3 log(mean) *− λ*. If the variability in the fit parameters can be explained by changes in *ν* alone (with fixed *η*^2^ and *a*), the plot of log(var) versus 3 log(mean) will be well-described by a line of slope 1 and fixed offset *−λ* = *−a*^2^*/η*^2^. SI Fig S5h shows this comparison when the mean and variance are computed from the fit parameters (filled markers) versus estimated directly from the data (open markers), along with a line of slope of 1 and best-fitting offset *λ_F_* = 0.61 (dashed line). We found that this provided a better fit than a model in which the bound is variable, and the drift rate and spread are fixed (dotted line).

We used this result to posit that fixations are controlled by a drift diffusion process with a fixed spread *η_F_* and bound *a_F_*, but an adaptive drift rate *ν_F_*. Because there are three parameters of the drift diffusion process but only two parameters needed to define the inverse Gaussian distribution, we are free to choose one of the drift diffusion parameters and fit the other two. We chose to set *η_F_* = 1, which requires that *a_F_* = 0.78 (thus satisfying 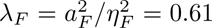). When we restricted our analysis to fixations that were initiated within the danger zone during the first 60 s of the first training trial (and remained within the danger zone for 95% of their duration), we found that they were well fit by the same process, but with a higher drift rate and thus shorter average duration (red star in SI Fig S5h). The reduction in fixation duration in response to heat can thus be captured by an additional “reflexive” drift process with drift rate *ν_F_* = 0.38, spread *η_F_* = 1, and bound *a_F_* = 0.78.

#### Characterizing saccade properties

Individual saccades varied in both their speed and duration. We found that the average duration of saccades depended on their average angular speed; we thus began by characterizing the distribution of angular speeds 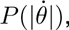 where 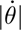 is computed by averaging the instantaneous difference in wingbeat amplitude over the duration of a saccade. The angular change in heading can be computed from this via 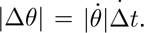 We then characterized the distribution of durations conditioned on speed, 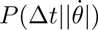.

Prior to fitting, we removed saccades whose speeds were below a threshold 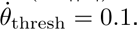 We used the MATLAB function *allfitdist.m* to fit 16 different parametric distributions to the distribution of speeds 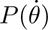 under two different conditions: when saccades were accumulated across flies within a given trial, and separately when saccades were accumulated across trials for a given fly. We performed this fitting for both laser-trained and no-laser control flies, and we used BIC to evaluate fits. We found that the lognormal distribution 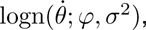 with location *ϕ* and scale *σ*^2^, was the best-fitting distribution across the majority of conditions. We then computed the directional bias in saccades, measured as (*N*_CW_ *− N*_CCW_)*/*(*N*_CW_ + *N*_CCW_), where *N*_CW_ and *N*_CCW_ respectively denote the total number of clockwise and counter clockwise saccades taken within a single trial. We found that this bias also varied across trials (SI Fig S5d upper), suggesting that flies can adaptively control directional bias of their saccades.

The distribution 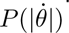 specifies the probability of initiating saccades of different speeds. For saccades of a given speed 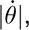 there is significant variability in their duration (SI Fig S5f). To characterize this variability, we considered 36 equally spaced values of 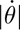 between 0.25 and 2. For each value of 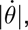 we used the MATLAB function *allfitdist.m* to determine the parametric function that best fit the distribution of saccade durations 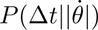 accumulated across flies, trials, and datasets. We found that these distributions were best fit by an inverse Gaussian distribution with fixed spread *η_S_* = 1 and bound *a_S_* = 0.56 but variable drift rate *ν_S_* (SI Fig S5i), analogous to fixations. In this case, the drift rate increased nonlinearly with the speed 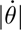 (inset of SI Fig S5i); we used a least-squares fit to determine the parameters of the best-fitting sigmoid 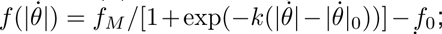 these where given by *f_M_* = 2.32, *k* = 7.55, 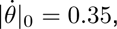 and *f*_0_ = *−*1.26. Thus, for a saccade initiated with speed 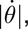 the duration can be generated via a drift diffusion process with a velocity-dependent drift rate 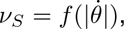 and fixed values of *η_S_* = 1 and *a_S_* = 0.56.

#### Inferring the structure of a behavioral policy

Together, the analysis of fixations and saccades enable us to construct a behavioral policy that accounts for variability in the initiation, speed, and duration of both fixations and saccades (SI Fig S5j). Each behavioral mode (fixation versus saccade) is generated via a sequence of three steps: (i) Select an angular velocity by sampling from a parametrized distribution. For fixations, we approximate the angular velocity to be zero. For saccades, we sample the magnitude of the angular velocity from a lognormal distribution, and we take the directional bias (corresponding to the likelihood of initiating a clockwise versus counter clockwise saccade, which specifies the sign of the angular velocity) to be an adaptive parameter. (ii) Generate the duration via an online drift diffusion process with a variable drift rate. For fixations, we take this drift to be an adaptive parameter. For saccades, this drift is determined by the angular velocity selected in step (i). (iii) Determine the resulting change in heading, which is proportional to the product between the average angular velocity and the duration.

In the circuit model used in the main text, we considered a simplification of this full behavioral policy in which the angular size of saccades (measured in deg) was directly sampled from a lognormal distribution with parameters *ϕ_S_* = 3.89 and *σ_S_* = 0.54 (this approximates the distribution of saccade sizes fit across all flies, trials, and datasets; this generates saccades with an median angular size of 49°), and assuming a fixed saccade duration of *t_S_* = 300 ms (this approximates the median saccade duration measured across all flies, trials, and datasets). Table 1 summarizes these choices.

#### Data selection

For all analyses shown in the main text, we used only those trials for which the fly exhibited at least 30s of continuous flight (these periods of continuous flight were defined using the set of saccades, fixations, drift, and “other” periods that we extracted from our segmentation; see *Behavioral Analysis Methods: Partitioning behavior into fixations and saccades*). For flies and trials that met this selection criterion, we kept all additional periods of flight that exceeded 3.788 ms (defined by the 75th percentile of fixation durations computed across all flies, trials, and datasets). This data was used in its entirety to compute PI scores. When analyzing fixations and saccades, we further excluded periods of drift; we then selected those fixations that exceeded a duration of Δ*t*_thresh_ = 200 ms, and those saccades that exceeded an average angular velocity of *ω*_thresh_ = 0.1 ms*^−^*^1^.

#### Assessing bearings relative to a single stripe

SI Fig S4 shows the strength and orientation of flies’ bearings relative to a single stripe. These were computed by taking the circular mean of the fictive heading directions acquired in each 30 s single-stripe trial (see *Experimental Methods: Flight visual learning*) using a custom MATLAB script modified from the Circular Statistics Toolbox [123]. To display the probability density estimate across flies, we adapted the MATLAB function *violinPlot.m* (Anne Urai, https://github.com/anne-urai/Tools/blob/master/plotting/violinPlot/violinPlot.m).

#### Measuring the strength of behavioral preference

Fig 1 shows the strength of behavioral preferences, measured with respect to the arena preferences of individual flies (Figs 1f and 6i) or with respect to the safe zone (Fig 1g,h,j,k). In both cases, we computed the preference strength using the performance index (PI) score. When measuring the strength of preference for safety, PI scores were computed as PI_safe_ = (*T*_safe_ *− T*_danger_)*/*(*T*_safe_ + *T*_danger_), where *T*_safe_ and *T*_danger_ denote the total time spent in safe and danger zones, respectively [30]. When measuring the strength of individual preferences, PI scores were computed analogously: PI_pref_ = (*T*_pref_ *− T*_anti_*_−_*_pref_)*/*(*T*_pref_ + *T*_anti_*_−_*_pref_), where the “preferred” and “anti-preferred” zones were defined analogously to “safe” and “danger” zones (i.e., arena headings were partitioned into two sets of preferred and anti-preferred zones, each spanning 90°), but were centered around the preferred arena heading measured in a given trial (see *Behavioral Analysis Methods: Aligning to individual preference* for details about determining the preferred arena heading). The left panels of Fig 1f,g,h,j,k show the strength of preference averaged across flies on each trial; the right panels show the strength of preference for data aggregated across trials 1-2 and trials 8-9. The upper right panel of Fig 6i shows the strength of preference for groups of flies that began with preferences that were stronger or weaker than the median initial preference strength.

#### Aligning to individual preference

Figs 1f, 3f, and 6i report features of the behavioral data computed after aligning the data to the arena preference of individual flies. To perform this alignment, we first computed the fraction of time that the fly resided at each of 96 orientations (corresponding to 96 pixel locations) within the arena. We then constructed an idealized residency profile that took a peak value of one at a central set of two pixels, and decayed linearly to zero over a span of 24 pixels in either direction (CW and CCW). We shifted this idealized profile with respect to the true residency profile of the fly, and we identified the preferred orientation as the one that maximized the overlap between the idealized and true residency profiles.

#### Computing heading-dependent averages

To perform the heading-dependent averages shown in Fig 3f, we first selected the set of saccades and fixations taken by each fly within naive trials (trials 1-2), and within training/probe trials (trials 3-9). We binned saccades according to the arena heading at which they were initiated; we binned fixations according to the average heading computed across the duration of the fixation. We used overlapping bins of width 11 pixels, centered on a given pixel. For each fly, we computed the average direction of saccades initiated within each bin, and similarly the average duration of fixations within each bin. For the data shown in Fig 3f, we included only those bins that had at least 2 samples per fly (either 2 fixations or 2 saccades per fly), among those, only those bins for which we had data from at least 5 flies. Fig 3f displays the mean and standard error in each of these quantities, computed across flies.

#### Measuring distance to safety

The middle righthand panel of Fig 6i shows the distance from the preferred arena heading to safety, averaged across flies. To compute this, we first aligned the behavioral data to the preferred arena heading for individual flies on individual trials, as described in *Behavioral Analysis Methods: Aligning to individual preference*. We then computed the minimum angular distance between this preferred heading and the center of the closest safe zone. Fig 6i shows the average and standard error of this distance for laser-trained flies.

### Bump Analysis

#### Computing calcium transients

For volume imaging of EPG GCaMP activity (Fig 2h), we used two z-planes that together captured the dorsal and ventral halves of the EB. The image stack at each time step was converted into a summed intensity projection that was used for further analysis. We manually divided the EB into 32 wedge-shaped ROIs to capture population EPG activity in the structure. An additional ROI without any EPG arborization and outside the EB was selected to estimate background signal, including from leaked LED arena light. Time series of GCaMP activity for all EB ROIs were obtained by taking the average of the fluorescence signal within each ROI at each time step. The calcium transient for each ROI, Δ*F/F*_0_, was computed by subtracting fluorescence in the background ROI from all other ROIs, and using the lowest 10th percentile of background-subtracted fluorescence from each ROI as F0. The resulting time series were Savitzky-Golay filtered with a 3rd order polynomial over 5 frames.

Rather than compute the population vector average (PVA), as in past work [16, 31, 47, 57, 58], we focused here on tracking peaks in EPG population activity (‘bump position’ or ‘compass heading’) at each time step. This allowed us to to track offsets between the EB location of the bump and the position of the visual scene at every time point, and to easily visualize changes in offsets, bump jumps, as seen in Fig 2h. Considering the symmetry of the visual scene, we tracked the position of the visual scene using two 180°-offset time series, with the first being shifted by the first offset and the second by a second offset (if present, see below).

#### Clustering bump offsets

The lefthand panel of Fig 2i shows the offsets between the bump and the visual scene for individual flies. To cluster these offsets, we used the MATLAB function *kde.m* (with 256 mesh points) to perform kernel density estimation of the distribution of offsets for each fly on each trial. We then used the MATLAB function *findpeaks.m* to determine the peaks in this density; we used offset values corresponding to these peaks as our candidate offset values. We then used the same function to determine the minima in this density (using a peak threshold of 10*^−^*^4^), and we used the offset values corresponding to these minima as the bounds between different clusters. We then computed the sum of the density function within these bounds, divided by the sum of the density function over all time, and used this as the fraction of time spent at each offset. The lefthand panel of Fig 2i shows the fraction of time spent at different offsets for individual flies on individual trials.

#### Characterizing the number and angular separation of offsets

The righthand panels of Fig 2i show the total number of and angular separation between offsets. To construct these histograms, we first computed the fraction of time that the HD bump spent at different offsets relative to the visual scene for each fly on each trial, as described above. We then computed the number of instances (aggregated over flies and trials) that we observed a given number of distinct offsets; these results are shown in the upper right panel of Fig 2i. For flies that exhibited two or more offsets on a given trial, we compute the angular distance between the dominant two offsets; this histogram is shown in the lower right panel of Fig 2i.

#### Computing HD tuning curves

Fig 2j shows the tuning of EB wedges to different arena headings. We first determined all times (aggregated across all 9 trials) that the visual scene was oriented at a particular angle relative to the fly, and then computed the average fluorescence transients Δ*F/F* of each wedge for each given scene orientation (see *Bump Analysis Methods: Computing calcium transients*). The HD tuning curves to the right of the main panel of Fig 2j show the average tuning of individual wedges as a function of the fly’s heading in the arena, i.e., the “arena heading”, which differs from the scene orientation by a sign flip.

#### Determining the locations of bump jumps with the EB

The righthand panel of Fig 5d shows the probability of bump jumps as a function of their location within the EB, measured relative to an inferred goal heading. To determine the location of the bump jumps, we first determined the relationship between the arena heading and bump phase that minimized the angular distance between successive time points. We took advantage of our previous results (shown in the lower right panel of Fig 2i) to select those changes in bump phase between 135° and 225°; this range captured the majority of the bump jumps in our data. For each jump, we marked the location within the EB at which the jump was initiated. We then computed the angular distance from this location to the location of the putative preferred compass heading within the EB (the “goal” location). To determine this location, we used the behavioral data for the same fly on the same trial to infer a preferred arena heading, as described above. For a given preferred arena heading, we determined the corresponding location in the EB at which the heading bump spent the most time. We used this as the location of the putative preferred compass heading in the EB. The righthand panel of Fig 5d shows the number of jumps that were initiated at a given angular distance from this goal location, divided by the total number of times that the bump visited locations of the same angular distance, accumulated across flies. To compute this histogram, we included for each fly only those trials for which the bump maintained two different offsets relative to the visual scene. To summarize these jump statistics, we determined the parameters of the best-fitting cosine function that minimized the mean-squared error between the measured and fit values of the histogram.

Note that the behavioral experiments were performed in arenas with a 330° angular span, but the imaging experiments were performed in arenas with a 180° span in the azimuth. Although we cannot rule out the possibility that the reduced horizontal span of the visual scene in imaging experiments affected the probability of the EPG bump jumping, similar bump dynamics have been reported in both flying and walking flies in symmetric visual settings in larger arenas as well [16, 31, 34].

### Modeling

#### Determining the optimal policy for maintaining a goal heading

Fig 3d shows the drift rate and average duration of fixations, and the turn bias of saccades, that result from training a flexible RL agent to exhibit a preference for a goal heading. The learning algorithm is described in detail in *SI: Reinforcement learning framework* (see Algorithm S7); the parameters used in the model are summarized in Table 1.

Briefly, we learned a single set of weights 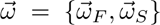 that specify the properties of fixations 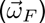 and saccades 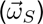 as a function of angular orientation (via a set of 16 von Mises radial basis functions), and we reported the resulting behavior when averaged over 100 different training runs. Reinforcement was delivered as a function of the current arena heading relative to a preferred heading; we assumed this reinforcement decayed linearly away from the preferred heading (see *SI: Reinforcement learning framework* for more details).

Prior to each training run, we initialized the set of flexible policy parameters 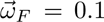 and 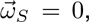 and we randomly initialized the arena heading to one of 96 evenly-spaced values between 0 and 360°. Following each training run, we used Eq. 17 to evaluate the drift rate of fixations, 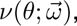 and the probability of rightward saccades, 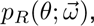 as a function of compass heading *θ* given the learned parameters 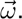 We then computed the average duration of fixations 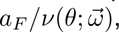 and the average turn bias of saccades 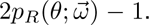 We averaged these across training runs to produce the curves shown in Fig 3d.

Fig 3e illustrates the expected behavioral readout if we couple the optimal policy to an unstable internal representation of heading. To illustrate this, we assumed that the internal heading could jump between orientations that correspond to symmetric views of the visual scene; for the two-fold symmetric scene used here, this corresponds to a jump of 180°. We further assumed that the jumps occurred probabilistically, and were least likely to occur at the preferred heading and most likely to occur at the symmetric (or “anti-preferred”) heading. We used a cosine function to parametrize this probability.

#### Summary of compass circuit model

We constructed a circuit model that could account for the tethering of the compass heading to the visual world, which is dictated by a set of plastic weights 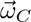 from inhibitory ring neurons onto compass neurons that are updated via anti-Hebbian plasticity. We assumed that these weights are updated during saccades based on the velocity of the saccade, the fly’s current compass heading during the saccade, and the current view of the visual scene [31]. We used these weights to determine the probability that the HD bump would jump between locations that correspond to symmetric views of the visual scene (see Eq. 27 in *SI: Reinforcement learning framework*). Bump jumps were assumed to occur immediately following a saccade. We assumed that these weights could be modified continuously, regardless of the presence or absence of reward/punishment.

#### Summary of policy circuit model

In addition to the compass circuit model, we constructed a policy circuit model that could implement the form of a behavioral policy, and flexibly modify its parameters. This model is described in detail in *SI: Reinforcement learning framework*. Briefly, we modeled a fly that can fixate and saccade. The duration of its fixations and the directionality of its saccades were determined by three populations of action neurons that receive phase-shifted input about the fly’s current heading (from a population of compass neurons) and input about the fly’s goal heading (from a population of goal neurons). Both the duration of fixations and the directionality of saccades were modulated by the strength of the goal heading, and by the fly’s current compass heading relative to this goal heading. The location and strength of the goal heading was determined by a set of plastic goal weights 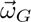 that could change over time based on the fly’s current compass heading and current level of reinforcement. In contrast to the compass weights (see *Modeling Methods: Summary of compass circuit model*), we assumed that the goal weights, 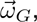 could only be modified in the presence of reward/punishment. Algorithms 3-4 show how we implemented this model.

#### Summary of circuit model simulations

In Figs 2, 5, and 6, we coupled the compass and policy circuit models (described above), such that the heading output of the compass circuit was used to modify goal and select actions within the policy circuit. We assumed that the compass heading was maintained as a von Mises activity profile in the compass circuit, and that it was transformed into a sinusoidal activity profile before passing into the policy circuit; we did not explicitly model this circuit transformation. In Figs 2 and 5, we assumed that the goal heading was fixed and that there was no reward/punishment; we used these simulations to study the evolution of the compass weights. In Fig 6, we mimicked the experimental setup and simulated two safe zones and two danger zones, each spanning 90° of heading space. We defined the centers of the safe zone to be at the arena headings *θ_A_* = *{*90*^°^,* 270*^°^}* = *{θ*_9_*, θ*_25_*}*. We used this model to simulate a period of training in which the compass and goal weights were co-evolving over time. Each training period consisted of a series of iterations, each consisting of a single saccade and a single fixation. Compass weights were iteratively updated at each angular increment (each value of *θ_i_*) during each saccade; compass weights were not updated during fixations. When rewards/punishment was being delivered, we subsampled periods of fixation into 100 ms increments, and iteratively updated the goal weights for each increment; goal weights were not updated during saccades.

The duration of fixations and directionality of saccades were determined by the current goal weights (see Eq. 35 in *SI: Reinforcement learning framework* for more details), and the sizes of saccades were sampled from a lognormal distribution with parameters *ϕ_S_* = 3.89 and *σ_S_* = 0.54 (approximating the values that were fit from data). We assumed that all saccades lasted a fixed duration of 300 ms (approximating the median duration observed in data). During probe periods in which the goal weights remained fixed, there was no simulated reward or punishment. During training periods, the model fly received a reward of +1 per unit time when in the safe zone, and a reward of *−*1 per unit time when in the danger zone.

#### Flexible mapping of visual scenes onto the compass heading

Figs 2 and 5 illustrate the synaptic weights from ring neurons onto compass neurons. We simulated the evolution of this weight matrix for 32 ring neurons and 32 compass neurons. This partitioned the space of both arena headings and compass headings into angular units of 360*^°^/*32 = 11.25°). Compass neurons were assumed to maintain a von Mises bump profile that was normalized to the range [0, 1], with concentration parameters *κ* = *π*, and whose location faithfully tracked changes in heading generated by the behavioral policy described above. Ring neurons were assumed to uniformly tile visual space with a receptive field width of three angular units (i.e., 33.75°), such that two adjacent receptive fields had an overlap of one angular unit. For asymmetric visual scenes (Fig 2b,c), we assumed that each ring neuron fired at a maximum rate of 1 whenever a fixed orientation of the visual scene aligned with the center of its receptive field, and fired at half of its maximum rate whenever the orientation of the visual scene was shifted by one angular unit to either side of its receptive field center. For symmetric visual scenes (Fig 2d,f and 5c-d), we assumed that the ring neuron exhibited these same firing patterns but with respect to two symmetric orientations of the scene separated by 180°. Weights were modified via an anti-Hebbian plasticity rule that weakens weights from active ring neurons onto active compass neurons (see Eq. 39 for the update rule). We updated these weights during saccades and in proportion to the squared velocity of each saccade, assuming that velocity was constant through the duration of each saccade.

All weight matrices in Fig 2b-d were generated for a single simulation that evolved from a randomly-initialized weight matrix; in these simulations, we did not allow the bump to jump, even for symmetric scenes. In Fig 2f, we froze the final weight matrix from 2d, and we used it to generate a short simulation of the heading and arena trajectories. We used the policy circuit model to generate fixations and saccades and assuming a fixed, sinusoidal goal profile normalized between 0 and 1 (see *Modeling Methods: Summary of policy circuit model* for details). In this simulation, we allowed the compass bump to jump by 180° following a saccade; the probability of a jump was determined by comparing the net inhibition from active ring neurons at the orientation of the compass heading versus the symmetric (i.e., 180° shifted) orientation.

In Fig 5c, we randomly initialized a weight matrix and allowed it to stabilize in an asymmetric visual scene (again using the policy circuit model to generate fixations and saccades, and again assuming a fixed, sinusoidal goal profile normalized between 0 and 1). We then changed the visual scene to a symmetric one, and simulated the evolution of the weight matrix in the new scene. The heatmaps in Fig 5c shows one such simulation; the panels below the heatmaps illustrate the probability that the bump would jump from any given orientation in the EB (as described above, this was determined by comparing the net inhibition from active ring neurons at the orientation of the compass heading versus the symmetric orientation).

In Fig 5d-e, we repeated the simulation shown in Fig 5c while varying the strength of the goal heading. In all cases, the goal profile was sinusoidal, normalized between 0 and a max amplitude *A*. We fixed the orientation of the goal profile but varied *A*, using values of [0.2, 0.4, 0.6, 0.8, 1]. For each amplitude, we simulated 200 model flies; the lefthand panels of Fig 5d-e report averages across groups of model flies with the same goal strength. We additionally analyzed the relationship between the final strength of the visual map and the final strength of behavioral preference (measured analogously to how we measure it in real flies; see *Behavioral Analysis Methods: Measuring the strength of behavioral preferences* for details). The upper righthand panel of Fig 5e shows the best linear fit between these quantities for the strongest goal heading, and the *r*^2^ values of this fit for each goal heading. The lower righthand panel of Fig 5e shows the temporal evolution of the behavioral preference for the strongest goal heading, and for two different initial conditions of the visual map.

#### A fixed-form behavioral policy tethered to a flexible goal heading

Fig 4b-e illustrates the behavioral policy whose heading-dependent structure is guaranteed by a multiplicative operation between the compass and goal activity profiles. We used a goal activity profile of 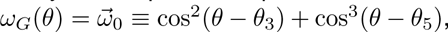 normalized to the range [0, 1]. We assumed a cosine profile of compass activity, also normalized to the range [0, 1]. The motor drive was determined by multiplying the compass and goal profiles and summing the output. We repeated this calculation for each possible circular shift of the compass activity to compute the net output as a function of current relative to goal heading (Fig 4c,d). When computing this for phase-shifted compass headings (Fig 4e), the phase shift was applied before the multiplication with the goal activity profile. We assumed that the compass weights were fixed and uniquely specified the arena heading.

Fig 4h illustrates the temporal evolution of the fixation duration and turn bias as we modify the goal heading via Hebbian plasticity. The Hebbian-like learning update that we used is given in Eq. 39. Fig 4h illustrates a continual updating of the goal weights given positive reinforcement at a fixed compass heading, and it illustrates the corresponding fixation durations and turn biases for each update (again computed via Eqs. 33-35).

#### Co-evolution of two learning systems

Fig 6d-h shows simulations in which both the compass weights and the goal weights are evolving over time. For all simulations, we first initialized the compass weights in an asymmetric environment. For this initialization, we assumed that individual model flies maintained a fixed set of goal weights that was sinusoidal in form, with a strength that was chosen to be one of 5 evenly spaced values ([0.2, 0.4, 0.6, 0.8, 1]), and with an orientation that was centered at *θ* = 0 (i.e., aligned with what would become the center of the danger zone). After initializing the compass weights, we introduce model flies to an environment that mimics the environment used in the learning assay, in which model flies are punished whenever they orient toward one of two repeating patterns in a visual scene. Fig 6d-h summarizes the evolution of the compass weights, goal weights, and PI scores over 1000 model flies (200 model flies for each goal strength). To assess the compass and goal weights, we track the orientation and strength of the most stable compass heading (computed as the circular mean of (1 *− P*_jump_(*θ*)), and described in the text as the orientation and strength of the visual map) and the goal heading (computed as the circular mean of the goal activity profile). PI scores were computed by freezing the compass and goal weights at a given time, running a separate simulation with these frozen weights, and using this residency within this simulation to compute an estimate of the PI score.

The vector maps in Fig 6d summarize the effects of learning across all simulations, as viewed through different quantities related to the orientation and strength of the goal heading and visual map. We binned all quantities into 15 evenly spaces bins; goal strengths were binned between 0.1 and 0.8; visual map strengths were binned between 0.1 and 0.6, the distance to safety was binned between 0 and 90°, and the map and goal orientation were binned between 0 and 360°. For each bin in a given vector map, we identified all instances where a simulation fell in the given bin, regardless of when this occurred during the simulation; the heatmaps in each vector map show the number of instances that fell into any given bin. For a given set of instances within a given bin, we determined the change in those instances over the next timestep of the simulation. In other words, for all instances of an initial goal and map strength, we computed the change in strength for each instance over the next timestep, and averaged these changes across instances. The vector maps show the magnitude and direction of this change, and the are colored by the average PI score of all instances in the bin. Vector maps were generated using the MATLAB function “quiver.m” with scales of 3, 2, and 2 for panels (d-1), (d-2), and (d-3), respectively.

Fig 6e shows the same vector map as in Fig 6d-3, but with individual trajectories superimposed. The selected model flies (which are also shown in the top panels of Fig 6f) were selected by randomly drawing one of the top 10 model flies that had the highest average PI scores over the entire course of the simulation. The model flies in the lower panels of Fig 6f were analogously selected by randomly drawing one of the bottom 10 model flies that had the lowest average PI scores over the entire course of the simulation. For each model fly, we reported the fraction of the simulation time needed to reach a goal strength larger than 0.6 and a distance to safety lower than 22.5°.

Fig 6g-h tracks the strength of the goal heading, the angular difference (or “misalignment”) between the goal and most stable compass headings, and the PI scores over time for all model flies. In all cases, we temporally aligned these trials to the time of the weakest goal heading (vertical dashed lines in Fig 6g), and we sorted trials by the weakest goal heading. We then grouped these ordered trials into equally-sized groups of 200 model flies, and averaged the same quantities over trials within each group; these averages are shown in Fig 6h. The righthand panels of Figs 6g-2 and 6h-2 show the average coherence of goal updates over time. For each model fly, we took the final goal heading at the end of the simulation, and we used this to compute the angular difference between the current and final goal heading as a function of time during the simulation. If the goal is shifting coherently toward its final location, we would expect this angular difference to decrease steadily over time. To measure this, we computed the variance in the distribution of angular differences at successive timepoints (i.e., we computed var(Δ*θ_t_ −* Δ*θ_t−_*_1_), where Δ*θ* is the angular difference between the current and final goal headings). We use this variance as a measure of the degree of incoherence in goal updates.

The simulations in Fig 6g-h were conducted identically to those in 6d-h, with one exception: instead of fixing the initial goal location, we randomly sampled it for each model by centering the goal heading on one of 32 randomly selected bins between 0° and 360°. We then partitioned model flies into two groups depending on whether their initial goal strength was larger or smaller than the median initial goal strength. For each group, we measured the average strength of behavioral preference, average distance between the goal heading and the center of the safe zone, and average PI score as a function of time throughout the simulation.

