## Supplementary Material for "A neural circuit architecture for rapid behavioral flexibility in goal-directed navigation"

### SUPPLEMENTAL FIGURES

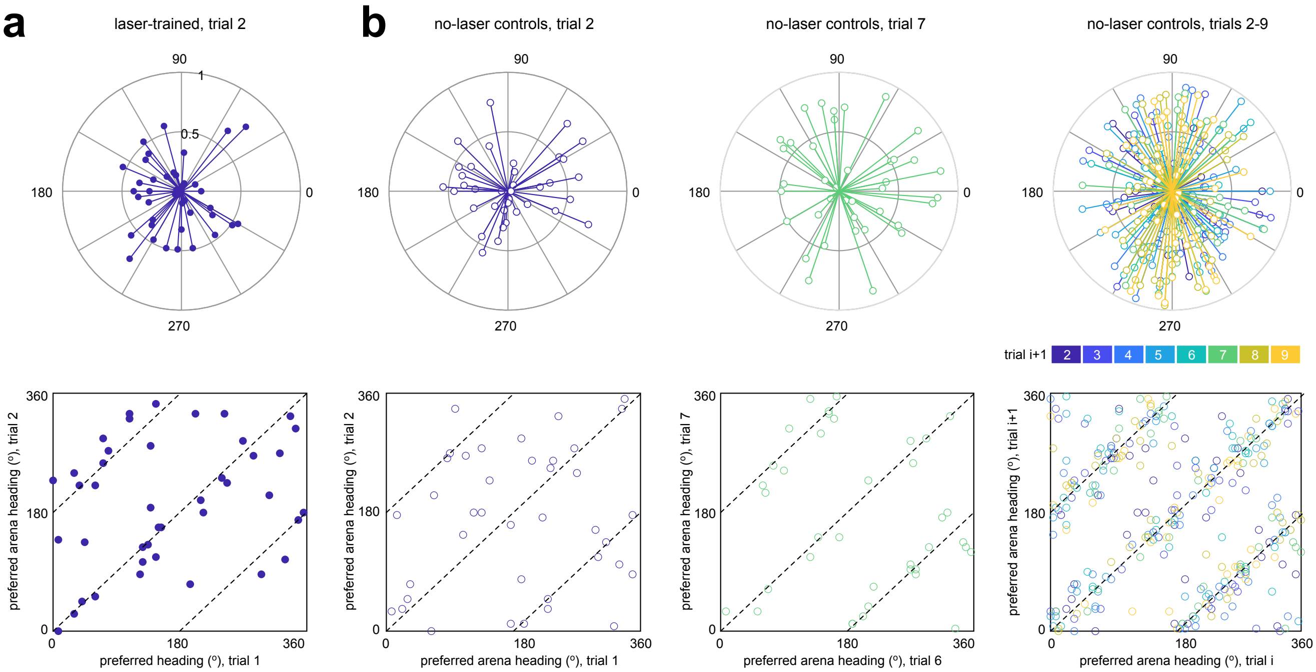


**Figure S1: Flies display variability in heading preferences. a)** Upper: angular location and strength of heading preferences for laser-trained flies, measured in trial 2 (see *Behavioral Analysis Methods: Aligning to individual preference* and *Behavioral Analysis Methods: Measuring the strength of behavioral preferences* for measurements of location and strength, respectively). Lower: comparison of angular location of heading preferences between trials 1 and 2. Due to the symmetry of the visual scene, we would expect preferences between successive trials to be similar up to a shift of $\pm 180°$ (dashed lines). **b)** Upper row: angular location and strength of heading preferences for no-laser control flies. Lower: comparison of angular location of heading preferences between successive trials.

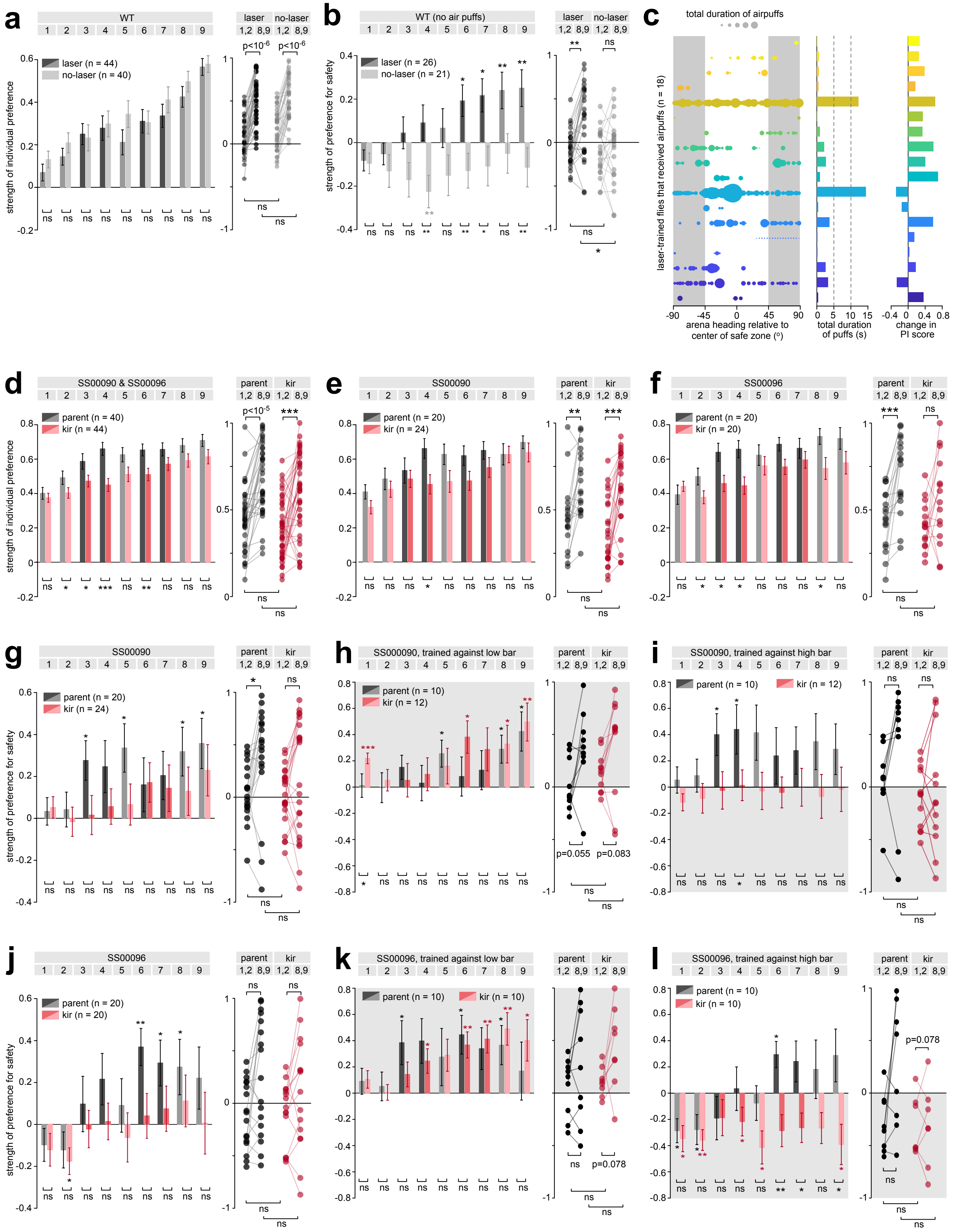
a
WT
laser
no-laser
1,2 8,9 1,2 8,9
laser (n = 44)
no-laser (n = 40)
strength of individual preference
p<10^-6
b
WT (no air puffs)
laser (n = 26)
no-laser (n = 21)
strength of preference for safety
c
total duration of airpuffs
laser-trained flies that received airpuffs (n = 18)
arena heading relative to center of safe zone (°)
total duration of puffs (s)
change in PI score
d
SS00090 & SS00096
parent
kir
parent (n = 40)
kir (n = 44)
p<10^-5
e
SS00090
parent (n = 20)
kir (n = 24)
f
SS00096
parent (n = 20)
kir (n = 20)
g
SS00090
parent (n = 20)
kir (n = 24)
h
SS000090, trained against low bar
parent (n = 10)
kir (n = 12)
p=0.055
p=0.083
i
SS00090, trained against high bar
parent (n = 10)
kir (n = 12)
j
SS00096
parent (n = 20)
kir (n = 20)
k
SS00096, trained against low bar
parent (n = 10)
kir (n = 10)
p=0.078
l
SS00096, trained against high bar
parent (n = 10)
kir (n = 10)
p=0.078

**Figure S2: Flies need their compass neurons to shift their heading preferences based on reinforcement. a)** Same as Fig 1f, but the strength of behavioral preferences are measured relative to the final heading preference in trial 9, rather than relative to the heading preference on individual trials. **b)** Same as Fig 1g, but using flies that never received any airpuffs on any trials. **c)** Left: 18 laser-trained flies (different colors) received airpuffs; for a given fly, each filled circle denotes the total duration of time that it was puffed at a given arena heading, accumulated across all trials. Middle: total duration of airpuffs, accumulated across arena headings and trials for each fly. Right: total change in PI score, measured by computing the average of PI scores in trials 8-9 and 1-2, and taking the difference of this average. **d-e)** Same as Fig 1f, but for genotypes SS00090 and SS00096. **g-l)** Same as Fig 1h,j,k, but split out by genotypes ('SS00090' for panels **(g-i)**; 'SS00096' for panels **(j-l)**).

4-way ANOVA

$\Delta PI \sim 1 + genotype + laser + pattern + kir + pattern*kir$

| factors | sum of squares | degrees of freedom | mean squares | F statistic | p-value |
|---|---|---|---|---|---|
| **genotype** | 0.0686 | 2 | 0.0343 | 0.197 | 0.821 |
| **laser** | 1.19 | 1 | 1.19 | 6.84 | 0.00983 |
| **pattern** | 0.0377 | 1 | 0.0377 | 0.217 | 0.642 |
| **kir** | 0.584 | 1 | 0.584 | 3.36 | 0.0690 |
| **pattern*kir** | 1.21 | 1 | 1.21 | 6.98 | 0.00917 |
| **error** | 25.213 | 145 | 0.174 | | |
| **total** | 28.624 | 151 | | | |

**Figure S3: Variance in PI scores is impacted by laser training and the combination of kir silencing and trained visual pattern.** 4-way ANOVA to compare the effects of genotype, laser training ('laser'), visual pattern on which flies were trained ('pattern'), and kir-silencing ('kir') on the change in PI scores (measured as described in the righthand panel of **(c)**). We considered a model with four main factors ('genotype', 'laser', 'pattern, 'kir') and one interaction term ('kir*pattern'). This revealed a significant effect of laser training and the interaction between kir-silencing and trained visual pattern (rows shaded in gray).

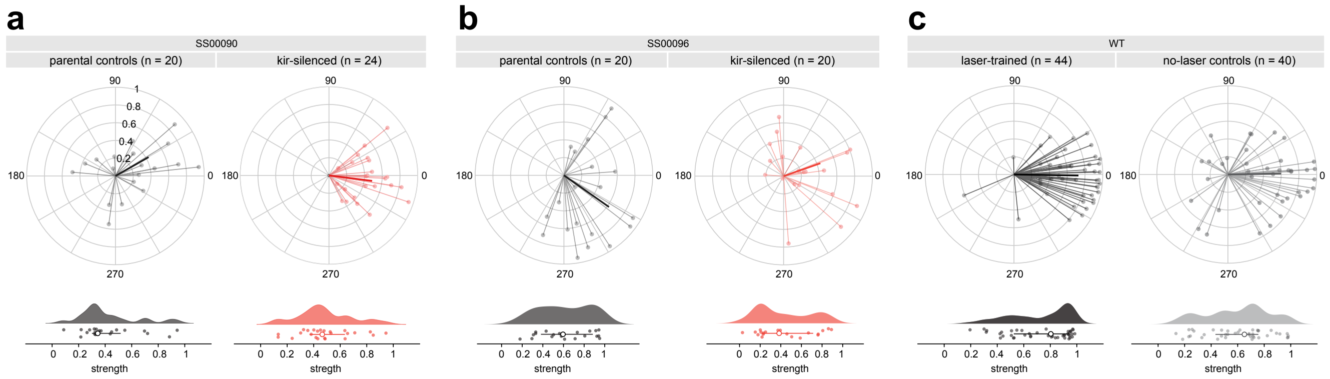


**Figure S4: Flies maintain steady bearings in single-stripe environments.** Upper: angular location and strength of bearings relative to a single stripe, shown for three groups: SS00090 parental controls and kir-silenced flies **(a)**, SS00096 parental controls and kir-silenced flies **(b)**, and WT laser-trained and no-laser control flies **(c)**. $0°$ and $360°$ correspond to maintaining the single stripe straight ahead. Lower: strength of bearing, shown for individual flies (dots) and summarized across each group in terms of the probability density estimate (shaded areas), median vector length (open circles), and quartiles (whiskers). See *Behavioral Analysis Methods: Assessing bearings relative to a single stripe* for details.

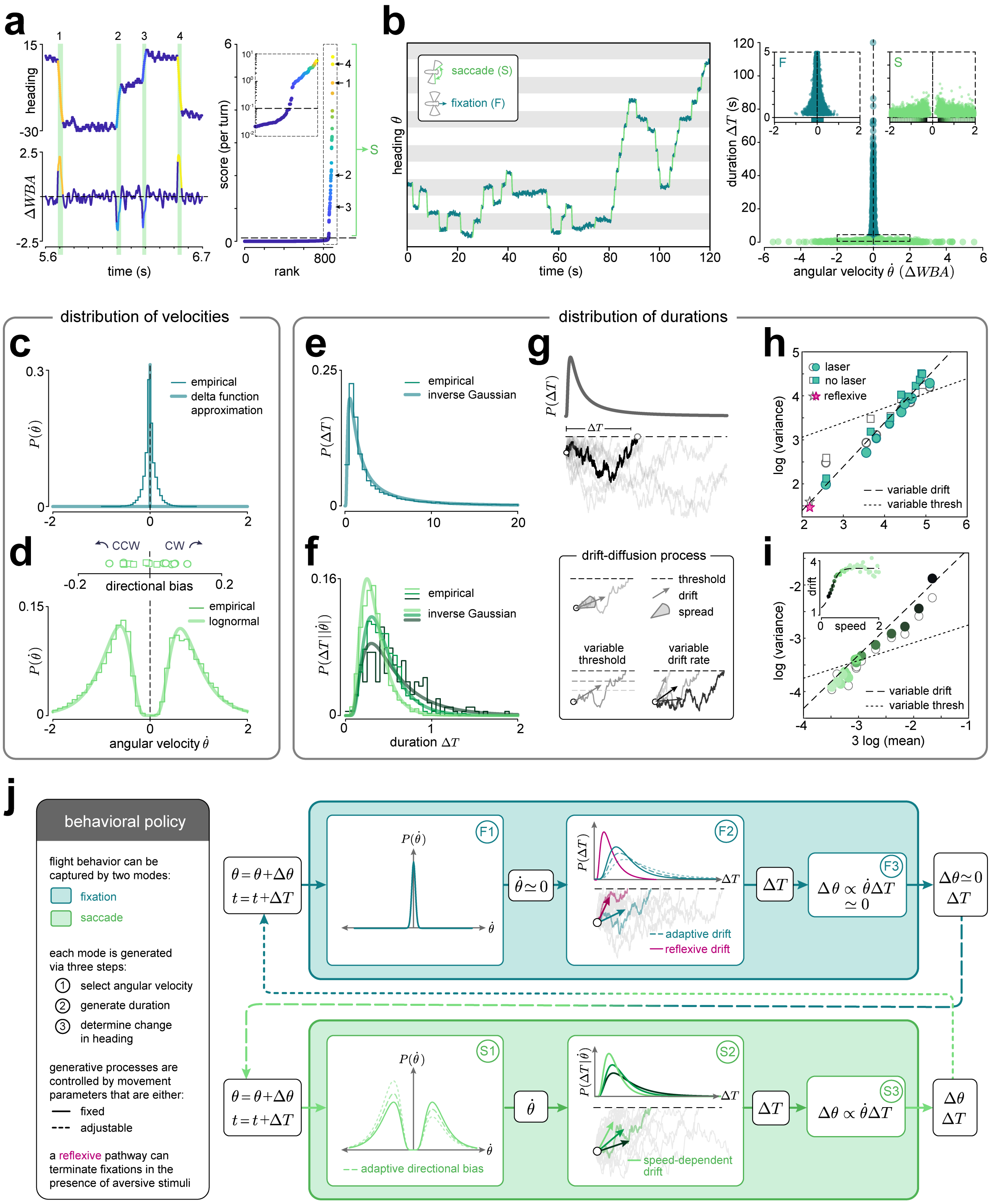

a
heading
time (s)
ΔWBA
score (per turn)
rank
S
b
saccade (S)
fixation (F)
heading θ
time (s)
duration ΔT (s)
angular velocity θ̇ (ΔWBA)
F
S
distribution of velocities
c
empirical
delta function approximation
P(θ̇)
d
CCW
CW
directional bias
empirical
lognormal
angular velocity θ̇
distribution of durations
e
empirical
inverse Gaussian
P(ΔT)
f
empirical
inverse Gaussian
P(ΔT||θ̇|)
duration ΔT
g
P(ΔT)
ΔT
drift-diffusion process
threshold
drift
spread
variable threshold
variable drift rate
h
laser
no laser
reflexive
log (variance)
variable drift
variable thresh
i
drift
speed
log (variance)
variable drift
variable thresh
3 log (mean)
j
behavioral policy
flight behavior can be captured by two modes:
fixation
saccade
each mode is generated via three steps:
1 select angular velocity
2 generate duration
3 determine change in heading
generative processes are controlled by movement parameters that are either:
fixed
adjustable
a reflexive pathway can terminate fixations in the presence of aversive stimuli
θ = θ+Δθ
t = t+ΔT
F1
P(θ̇)
θ̇ ≃ 0
F2
P(ΔT)
ΔT
adaptive drift
reflexive drift
F3
Δθ ∝ θ̇ΔT ≃ 0
Δθ≃0
S1
adaptive directional bias
S2
P(ΔT|θ̇)
speed-dependent drift
S3
Δθ ∝ θ̇ΔT
Δθ

**Figure S5: Inferring a behavioral policy. a)** Example detection of saccades from a portion of the heading trajectory shown in (**b**) (and also shown in Fig 3a). Left column: example saccades, numbered and colored according to score (right column; explained below). Right column: Individual saccades are scored based on changes in heading and wing beat amplitude (see *Behavioral Analysis Methods: Partitioning behavior into fixations and saccades*). Turns with scores above a threshold (dashed line) are defined as saccades ('S'). Periods between saccades are defined as fixations ('F'). Inset: same data, shown using a log scale on the score. **b)** Left: Heading trajectory partitioned into fixations (dark green) and saccades (light green). Right: Distribution of duration $\Delta t$ and angular velocity $\dot{\theta}$ of individual fixations and saccades, accumulated over $9$ trials across laser-trained ($n = 44$) and no-laser control ($n = 40$) flies. For each event (fixation or saccade), $\Delta t$ measures the entire duration of the event; $\dot{\theta}$ is measured as the change in wing beat amplitude, averaged across the event. Insets: distribution of events with short durations and low angular velocity; colored bars along horizontal axis indicate $95\%$ confidence intervals. **c)** Empirical distribution of fixation velocities (using data in **b**), approximated as a delta function. **d)** Empirical distributions of clockwise and counter clockwise saccade velocities (using data in **b**), and best-fitting lognormal approximations. Inset: directional bias of saccades; each marker shows the bias measured across flies within individual trials. **e)** Empirical distribution of fixation durations (using data in **b**), and best-fitting inverse Gaussian approximation. **f)** Empirical distribution of saccade durations conditioned on different saccade velocities (using data in **b**), and best-fitting inverse Gaussian approximations. **g)** Illustration of drift-diffusion (DD) process for generating inverse Gaussian (IG) distribution of durations. For a given choice of the mean drift rate, spread, and threshold of the diffusion process, the time of first threshold crossing ('first passage' time) is IG-distributed. **h)** Properties of the within-trial across-fly distributions of fixation durations, measured from the best-fitting IG distributions (green filled markers) and estimated empirically (gray open markers), compared against a model in which the variability in these same properties arises from changes in either the drift rate (dashed line) or threshold (dotted line) of a DD process (see schematic in **g**). Star: distribution of fixation durations estimated from fixations made within the danger zone during the first $60$ s of the first training trial. **i)** Same as **h**, but shown for properties of the across-trial across-fly distributions of saccade durations conditioned on different angular speeds. Inset: the drift rate of the best-fitting DD process is nonlinearly related to the average angular speed of saccades, and can be fit with a sigmoidal function (dashed line; see *Behavioral Analysis Methods: Characterizing saccade properties*). **j)** Left: ingredients of behavioral policy. Right: schematic of policy consisting of transitions between fixations (dark green box) and saccades (light green box). Fixations are initiated with zero angular velocity (F1), and the duration of a given fixation is generated on-line via a DD process with an adaptive drift rate (F2). When receiving punishment, this process can be short-circuited by a DD process with a reflexive drift rate. The fixation is terminated when either the adaptive or reflexive DD process first crosses a fixed threshold, leading to a change in time but no change in heading (F3). Following the termination of a fixation, a saccade is initiated. The angular velocity of the saccade is sampled from a lognormal distribution with adaptive directional bias (S1), and the duration of the saccade is generated via a DD process whose drift rate depends on this angular velocity (S2). The saccade is terminated when the DD process first crosses a fixed threshold, leading to a change in both time and heading (S3). Following the termination of a saccade, a fixation is initiated, and the process repeats.

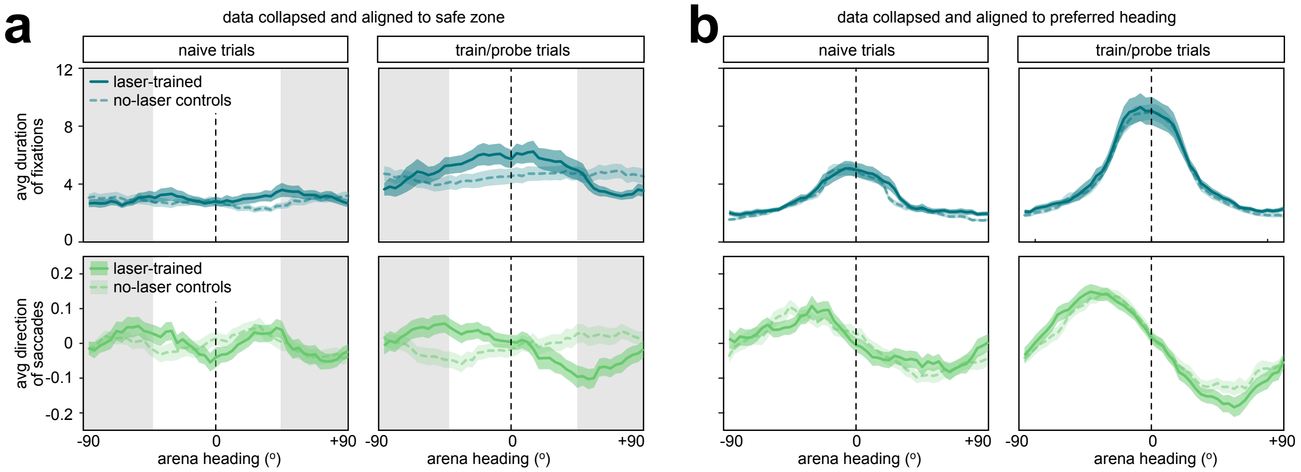


**Figure S6: Laser-trained flies change their behavior relative to safety and danger. a)** Average duration of fixations (upper panel) and direction of saccades (lower panels) generated by in laser-trained ($n = 44$) and no-laser control flies ($n = 40$), measured before training (naive trials; left column) and after training (probe trials; right column). All data is collapsed onto half of the virtual heading space and aligned to the center of the safe zone (vertical dashed line). Dark lines and shaded regions mark the mean +/- s.e.m. **b)** Same as **(a)**, but aligned to the preferred arena headings of individual flies (vertical dashed line).

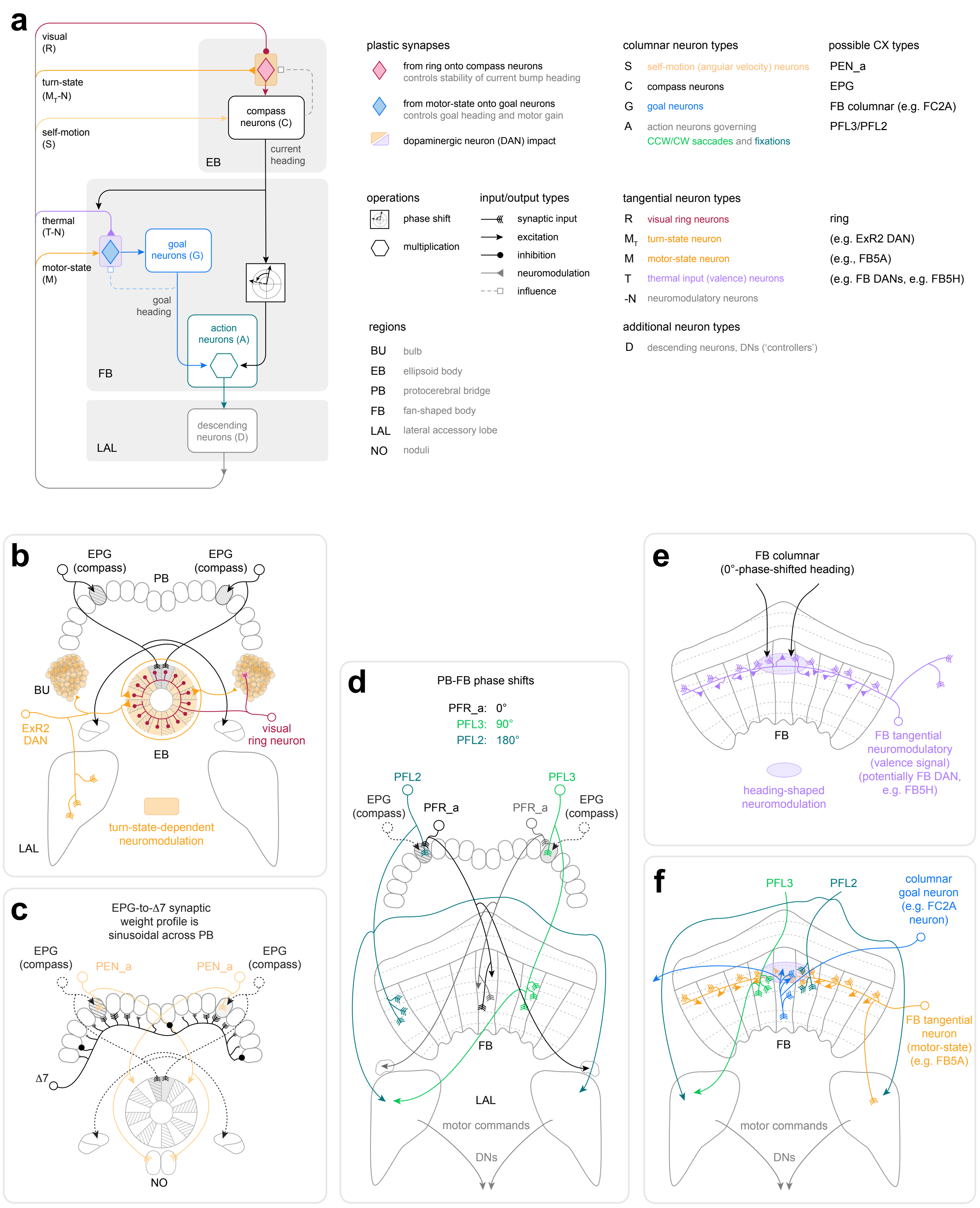

a
visual (R)
turn-state ($M_T$-N)
self-motion (S)
compound neurons (C)
current heading
EB
thermal (T-N)
motor-state (M)
goal neurons (G)
goal heading
action neurons (A)
FB
descending neurons (D)
LAL
plastic synapses
from ring onto compass neurons
controls stability of current bump heading
from motor-state onto goal neurons
controls goal heading and motor gain
dopaminergic neuron (DAN) impact
operations
phase shift
multiplication
input/output types
synaptic input
excitation
inhibition
neuromodulation
influence
regions
BU bulb
EB ellipsoid body
PB protocerebral bridge
FB fan-shaped body
LAL lateral accessory lobe
NO noduli
columnar neuron types
S self-motion (angular velocity) neurons
C compass neurons
G goal neurons
A action neurons governing CCW/CW saccades and fixations
possible CX types
PEN_a
EPG
FB columnar (e.g. FC2A)
PFL3/PFL2
tangential neuron types
R visual ring neurons
$M_T$ turn-state neuron
M motor-state neuron
T thermal input (valence) neurons
-N neuromodulatory neurons
ring
(e.g. ExR2 DAN)
(e.g., FB5A)
(e.g. FB DANs, e.g. FB5H)
additional neuron types
D descending neurons, DNs ('controllers')
b
EPG (compass)
PB
BU
ExR2 DAN
visual ring neuron
EB
turn-state-dependent neuromodulation
LAL
c
EPG-to-Δ7 synaptic weight profile is sinusoidal across PB
EPG (compass)
PEN_a
Δ7
NO
d
PB-FB phase shifts
PFR_a: 0°
PFL3: 90°
PFL2: 180°
PFL2
PFL3
EPG (compass)
PFR_a
FB
LAL
motor commands
DNs
e
FB columnar (0°-phase-shifted heading)
FB
FB tangential neuromodulatory (valence signal) (potentially FB DAN, e.g. FB5H)
heading-shaped neuromodulation
f
PFL3
PFL2
columnar goal neuron (e.g. FC2A neuron)
FB tangential neuron (motor-state) (e.g. FB5A)
FB
motor commands
DNs

**Figure S7: Support for the circuit model from central complex anatomy. a)** In our model, CX circuits drive behavior based on the overlap between the current and goal headings, computed by multiplicatively combining goal neuron activity with phase-shifted compass neuron activity. Plasticity in weights onto compass neurons (red diamond) and goal neurons (blue diamond) enables the co-evolution of internal mappings of sensory surroundings and of goals within those surroundings. As illustrated in the remaining panels, this circuit model relies on abstracted elements and circuit motifs of the fly CX; see *SI: Linking the Conceptual Model to Known Anatomy* for more details. Note that although the compass neurons are thought to carry HD information, the constraints of our tethered flight setup made heading equivalent to HD. *SI: Linking the Conceptual Model to Known Anatomy* also discusses how the model might also be relevant for, and operate on, true heading (traveling direction). **b)** Flexible mapping of a visual scene onto the fly's HD representation. The fly's HD representation tethers to sensory cues in the environment, which are conveyed to the ellipsoid body (EB) through multiple classes of ring neurons. Visual ring neurons (red) receive inputs in the bulb (BU) and project to the the EB, where they make all-to-all inhibitory connections onto, and receive feedback from, compass (EPG) neurons. Plasticity between ring and compass neurons ensure a self-consistent mapping between the sensory world and the internal HD representation. ExR2 dopaminergic neurons (DANs, orange) receive input in the lateral accessory lobe (LAL) and project to the EB and both BUs, making these DANs a key source of motor-state-dependent neuromodulation in the EB. This neuromodulation has recently been shown to depend on angular velocity in walking flies and to drive plasticity in the mapping between ring and compass neurons [70]. **c)** Maintaining, updating, and formatting the HD bump. In addition to being tethered to visual and other sensory input, the location of the HD bump in the EB is also determined by multiple classes of columnar neurons (PB-EB neurons) that link the protocerebral bridge (PB) and the EB. Most notably, PEN_a neurons tuned to HD and the fly's angular velocity update the HD bump's position based on self-motion input. Note that their projection pattern from the PB to the EB ensures that their input is phase-shifted relative to their compass neuron inputs. The sinusoidal weight profile of compass neurons onto downstream $\Delta$7 neurons, which inhibit other compass neurons, ensures that the HD bump maintains a sinusoidal shape. **d)** PB-FB columnar neurons that link the PB and the fan-shaped body (FB) have a range of phase shifts. The PFR_a neurons are shown as an example of a PB-FB columnar neuron type that has a $0^\circ$ phase shift between its PB and FB projections. In other words, a bump inherited from the compass neurons in the PB would be transferred to the matching column of the FB. This is in contrast to the PFL2 neurons, which project to FB columns that are $180^\circ$ shifted in phase relative to their PB arbors, and to the PFL3 neurons, whose projection patterns produce a $90^\circ$ contralateral phase shift. These phase shifts provide the basis for the fixed-form behavioral policy in the model. Note also that PFL2 neurons project to the LAL on both sides of the brain, thereby making them suitable for fixation control, and that the PFL3 neurons project unilaterally, making them ideal saccade controllers. Recent experimental studies have highlighted the roles of PFL2 [85] and PFL3 neurons [84, 85] in controlling the fly's movements. Both neuron types project to the LAL, which is innervated by multiple classes of descending neurons (DNs) that project to motor centers in the ventral nerve cord. **e)** Valence signals shaped by an HD or heading bump. The FB is innervated by numerous tangential neurons that project throughout specific FB layers, where they show both presynaptic and postsynaptic specializations. Specifically, many FB tangential neurons receive columnar inputs from neurons that could carry HD or heading bumps without the phase shifts that characterize PFL FB activity. Some FB tangential neurons are known to be neuromodulatory; for example, FB DANs like FB2A, FB4L, FB4M, and FB5H, and may carry valence signals, such as those associated with heating or cooling. Their neuromodulatory signals in the FB are likely to be shaped by their local columnar input (see *SI: Linking the Conceptual Model to Known Anatomy* for more details), motivating our assumption of heading-shaped neuromodulation as driving plasticity in the goal weights. **f)** A circuit motif for updating goal weights. In our model, FB tangential neurons carrying a heading-shaped valence signal drive plasticity at synapses between motor-state-dependent tangential neurons and largely intrinsic FB columnar neurons that act as goal neurons. Potential candidates for such motor-state neurons include FB5A neurons that receive input in the LAL, including from PFL2 and PFL3 neurons. These and other candidate FB tangential neurons receive input from FB DANs and other neuromodulatory FB tangential neurons in the FB, and themselves make synapses onto multiple classes of FB columnar neurons that could function as the goal neurons we assume in our model. Reinforcement would modify the strengths of these synapses, leading to different profiles of goal-weighted activity in the columnar neurons when the fly is active. Here we show one possible FB columnar neuron type, FC2A, which receives input from tangential neurons in multiple layers of the FB, makes synapses onto the putative action neurons, PFL2 and PFL3, and has recently been shown to exhibit goal-neuron-like properties in the context of menotaxis in walking flies [84]. In our model, this circuit motif would allow individual PFL neurons to multiplicatively combine information from the goal neurons with phase-shifted versions of the HD bump, thereby enabling the population of PFL neurons to enforce the form of the behavioral policy for structuring saccades and fixations relative to the goal.

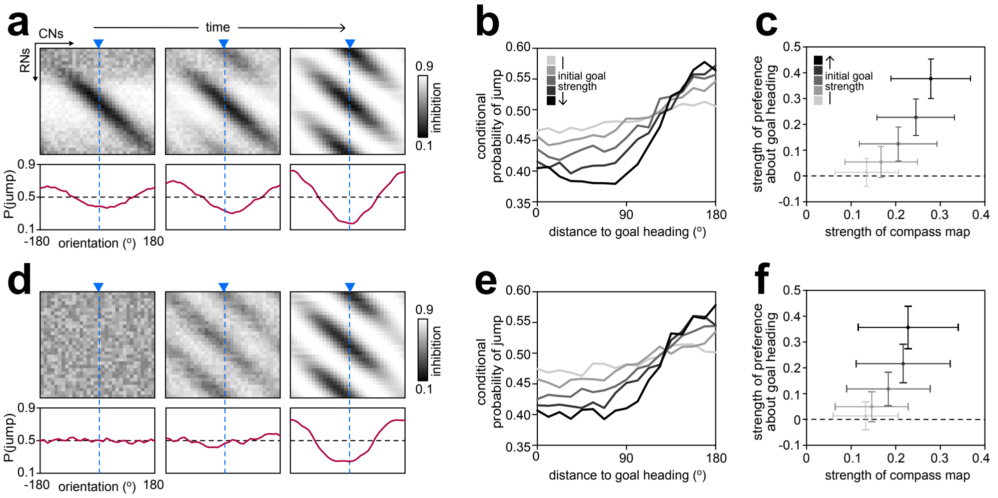


**Figure S8: Different initial visual maps align to an internal goal heading after experience in a symmetric visual scene.** Same as Fig 5c-e, but shown for model flies that began with a visual map developed in an asymmetric scene (panels **(a-c)**) compared with flies that began with a random visual map (panels **(d-f)**).

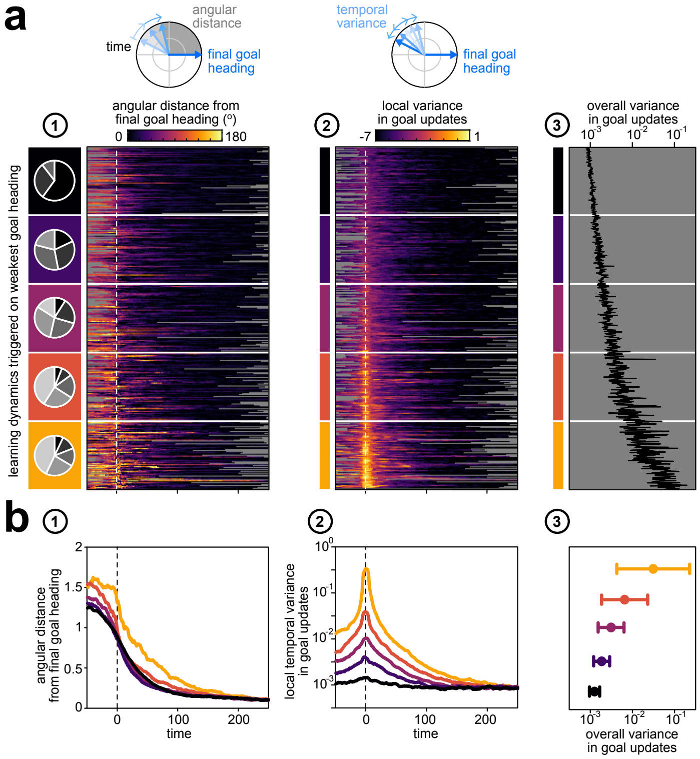


**Figure S9: Misalignment between visual maps and goal headings can lead to inconsistent goal updates. a)** Properties of learning trajectories over time, aligned as in Fig 6g. Colored rectangles indicate the groupings that were used to construct the averages shown in **(b)**. (1) Prolonged misalgnments between the visual map and the goal heading (Fig 6g-2) slow the updating of the goal heading, measured here by computing the angular distance between the current and final goal headings. (2) The convergence of the goal heading can be measured by tracking the rate of change of the angular distance between the current and final goal headings. Fast convergence is marked by a continuously decreasing angular distance; slow convergence is marked by an angular distance that increases and decreases over time. We use the local temporal variance in this rate of change (computed using a sliding window of 10 successive timesteps) as a measure of how consistently the goal heading is shifting toward its final value. (3) Weaker goal headings lead to higher overall variance in goal updates. Here, variance is computed over the entire duration of the simulation. **b)** Same as **(a)**, but averaged over groups of flies that exhibited similar dynamics in their goal headings (colored groupings highlighted in panel **(a)-1**). Model flies that began with weaker goal headings are slower to reach their final goal heading (panel 1) because they show higher temporal variance in goal updates (panels 2-3).

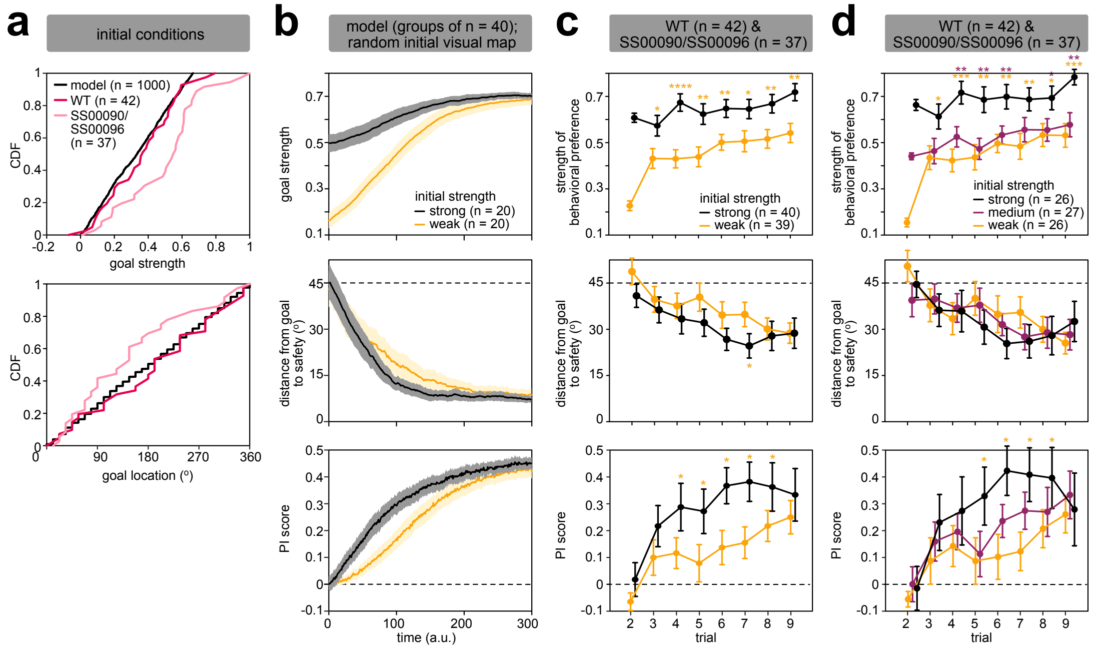


**Figure S10: Different initial visual maps align to an internal goal heading after experience in a symmetric visual scene. a)** Distribution of initial conditions used in model simulations in the left column of Fig 6i, compared to the analogous quantities measured in real flies in trial 2. **b)** Same as the left column of Fig 6i, but model flies began with a random visual map, instead of a visual map developed in an asymmetric visual scene. **c)** Same as the right column of Fig 6i, but including WT flies ('laser-trained') and SS00090 and SS00096 flies ('parental controls'). Flies were divided into two groups based on whether the strength of their behavioral preference was greater or less than the median as measured in trial 2. **d)** Same as **(c)**, but flies were divided into three groups based on whether the strength of their behavioral preference fell into the bottom 33%, middle 33%, or top 33%, as measured in trial 2. Significance in panels **(c-d)**: two-sided Wilcoxon rank sum test (*$p \leq 0.05$; **$p \leq 0.01$; ***$p \leq 0.001$) against the null hypothesis that scores measured for flies with strong versus weak (yellow stars) or intermediate (purple stars) initial preferences come from continuous distributions with the same medians.

### SUPPLEMENTAL INFORMATION

#### Linking the Conceptual Model to Known Anatomy

Our model (SI Fig S7a) builds on physiological, behavioral and connectomic findings from many laboratories, as well as on the many conceptual ideas and models that have been proposed for the CX in recent years. Although our model focuses on the CX, we note that many visual learning behaviors, including those associated with navigation, may also involve the mushroom body [43, 81, 95, 97]. In addition, the task we employ and most other orienting tasks almost certainly recruit more direct sensorimotor pathways as well, for example, when the fly responds to aversive heat in our task. We chose to exclude such parallel pathways in our modeling framework to more thoroughly investigate the dynamics of early learning driven by internal representations. In the CX, several of our modeling assumptions are based on physiological studies of different CX neuron types during visual stimulation and behavior. Importantly, although not all the key features of our circuit model have physiological support, they are all inspired by the known anatomy and connectivity of the CX. However, rather than incorporating all the known details of CX connectivity, we greatly simplified and abstracted the circuit in order to focus on a few key computations that we believe underlie much of the fly's behavior in the visual learning paradigm. We now discuss the many simplifications that we made, and summarize what is known about several neuron types and network motifs that are likely to play a role in relevant CX circuit computations.

**The flexible mapping from visual scene to HD representation: Recurrent connections between ring neurons and EPG neurons.** The HD system is tethered to its sensory surroundings by multiple ring neuron classes —many with tens of ring neurons each— that carry information about sensory cues, such as visual features [65–67, 124], polarized light patterns [125–127], and wind direction [128]. Most of these neurons are thought to be GABA-ergic and inhibitory [129, 130]. Some visual-feature-sensitive ring neurons have spatiotemporal receptive fields [67], and are thus likely to be sensitive to how a visual feature moves across the fly's eyes, a factor that we ignored in our model. Sensory ring neurons are connected all-to-all to other ring neurons of the same class, and, in some cases, across classes as well, and most sensory ring neurons receive feedback from the compass (EPG) neurons [29]. These motifs may ensure that the fly's HD representation tethers to the strongest cues available [29], which we assume to be less relevant in a visual setting with four identical horizontal bars. Plasticity in the synapses between ring and compass neurons has been hypothesized to create a flexible mapping between sensory cues and the HD representation [62, 63], an idea similar to one proposed for the rodent HD system [131] (SI Fig S7b). Recent experimental results strongly support this idea of plasticity between visual inputs and compass neurons [31, 34]. As part of a model proposed in one of these studies [31], we assumed that this plasticity depends on an inhibitory Hebbian-like rule that relies on correlated activity between visual and compass neurons during saccades, and results in changes in the depth of inhibition that compass neurons receive at different angular orientations in their surroundings [34]. Plasticity in the EB may involve nitric oxide signaling [132] and motor-state-dependent neuromodulation. There are multiple sources of neuromodulation in the EB, including the ExR2 dopaminergic neurons (SI Fig S7b), which receive inputs in the LAL and project to the BU and EB (see Figure 14 and associated figure supplements in [29]) and have been linked to circadian changes in locomotion [133]. Recent experimental results show that the ExR2 neurons indeed play a key role in turn-dependent dopaminergic modulation of plasticity during scene mapping, at least in walking flies [70].

Several experimental studies [16, 31, 34] have reported variability in the EPG bump's offset relative to the fly's surroundings in symmetric visual settings. Two visually indistinguishable headings are likely to evoke similar ring neuron population activity, making both EB locations corresponding to those headings viable for the bump to occupy. Which one of those locations the bump resides in would depend on the relative strength of the ring-neuron-to-EPG mapping in those two EB locations. In our behavioral paradigm, we found that some EB locations were more likely to feature bump jumps (see right panel of Fig 5d), and that these bump jumps tended to be $180^\circ$ in magnitude (see lower right panel of Fig 2i), reflecting the symmetry of the scene. Note that the vertical span of some ring neurons' receptive fields may be large enough [65] to evoke responses to horizontal bars at both high and low elevations, making the scene weakly symmetric at $90^\circ$ from the perspective of those inputs to the compass neurons, and triggering a few $90^\circ$ bump jumps as well (small peak in bottom right panel Fig 2i). We explicitly modeled the inhibitory interactions from ring neurons onto EPG neurons, and assumed that the relative, experience-dependent strength of the HD representation in $180^\circ$-opposite EB positions determines the probability of an EPG bump jump between them. Note that these differences in the strength of summed ring neuron inhibition onto EPG neurons at $180^\circ$-opposite EB locations would not necessarily lead to EPG bump amplitude differences at those locations, because of other sources of broad feedback inhibition onto EPG neurons in both the EB and the PB [29, 47, 60,

134].

**Maintaining and updating the HD representation: A ring attractor circuit involving EB and PB neurons.** There are ∼48 EPG neurons, which each occupy one of 16-18 compartments in the EB and PB [29, 135]. In our model, we assumed that the HD representation is carried by 32 EPG-like neurons, each with distinct HD tuning equally spaced across $360°$. The dynamics of the HD representation match those produced by ring attractor networks [47, 57, 58]. These dynamics depend not just on sensory inputs, but also on self-motion input from the PEN_a (SI Fig S7c) and other columnar neurons that link the EB and PB [47, 58, 59]. These self-motion inputs are also important for the mapping of visual scenes onto the EPG population [31]. In our model, the HD representation was entirely driven by visual input through the ring neurons, and we ignored recurrent connections involving the PEN and PEG neurons, as well as the intra-EB connections between different EPG neurons. These connections could, in effect, tether the EPG bump more strongly to locations near its current location, reducing the probability of bump jumps even if the relative strength of connections from ring neurons to different EPG neurons make them more likely. Further, recent extracellular recordings from candidate EPG-like neurons in monarch butterflies suggest that bump jumps may be less likely when animals are allowed to physically turn [71] rather than being in closed-loop VR as in our imaging experiments.

**Neurons that reformat the HD bump and link the PB to the FB** Our model used a von Mises function to represent the shape of the EPG population's HD bump in the EB, but assumed that the bump shape becomes sinusoidal as it goes through the PB. Note that we did not explicitly include connections from EPG neurons onto the broadly arborizing PB neurons that are thought to be key to ensure the sinusoidal shape, the $\Delta 7$ neurons [29, 60] (SI Fig S7c). Both the EPG neurons and $\Delta 7$ neurons contact a wide range of columnar neurons in the PB. Some of these neurons project back to the EB, but most are FB columnar neurons [29], such as the PFNs and PFRs [129, 136, 137] (SI Fig S7d). Many of these FB columnar neurons receive additional inputs in other CX structures [29, 60, 83, 138, 139]. In combination with neuron-type-specific anatomical phase shifts in their projection patterns from individual glomeruli in the PB to columns of the FB [29, 60, 139] (see SI Fig S7d for illustrative examples), these inputs likely allow the PB-FB columnar neuron types to participate in vector computations that transform the HD representation in different ways [29, 60, 139]. This may be highly relevant to the computations that flies use when navigating over long distances in natural conditions [49, 53]. In such natural settings, flies would likely need to select actions to maintain a particular traveling direction ('goal heading') rather than to maintain a specific head direction ('goal HD'). Recent conceptual insights from the connectome [29] and, in parallel, confirmatory evidence from physiological experiments [60, 139] suggest that some FB columnar neurons use translational self-motion cues to transform HD into an explicit representation of traveling direction (heading). At the end of this Supplemental section, we discuss how a potential circuit mechanism to learn and express a heading (traveling direction) preference that is robust to perturbations from wind might be implemented in FB circuitry. However, in a head-fixed preparation in which the fly only controls and receives visual feedback for its angular movements, the distinction between HD and heading is less relevant. Thus, although we use the term 'heading' in the main text and below, our model did not incorporate mechanisms that would be necessary for this flexible behavior to operate in the space of heading rather than HD.

**Learning and storing a goal heading: Candidate neurons and circuitry in the FB.** Recent studies have proposed simple conceptual models for how goal headings might be stored in the strength of synaptic connections between neurons of the FB [29, 36]. An entirely different model for visually-guided homing is that heading-dependent views or visual snapshots are stored in the strength of synapses between visually-responsive Kenyon cells and mushroom body output neurons (MBONs) in the MB [93, 94, 96, 140]. There is, as yet, only indirect experimental evidence for the MB model [95, 141]. However, there is evidence that flexibility in goal headings depends on visual input to ring neurons [47] and on output from EPG neurons [33, 35]. Further, this flexibility does not involve changes in the mapping between the visual scene and the EPG HD representation [35], suggesting that goal headings are stored downstream of the EPG neurons. The behavioral genetics evidence implicating FB neurons in a visual learning task that inspired ours [32] suggest that goal headings may be stored in plastic synapses between neurons in the FB.

We assumed that the goal heading is stored in the strength of synapses from hypothesized tangential motor state neurons to putative columnar 'goal neurons' (SI Fig S7e-f). Such a mechanism would be metabolically efficient, allowing goal neurons to be activated into a goal-heading pattern specifically when the fly is moving, but not otherwise. We further assumed that these synaptic strengths are modified through the action of neuromodulatory tangential FB neurons, which, we assume, deliver reinforcement signals shaped like the current heading bump (SI Fig S7e). There is already some physiological evidence for reinforcement signals in the FB [142], although the neuronal players involved are as yet unknown. Prime candidates for such a role are the ventral FB dopaminergic neuron (DAN) types. MB DANs are known to carry reinforcement (and movement) signals and be involved in associative learning in that brain region [143–147], and a subset of FB DANs —potentially the more ventral FB DANs, such as FB2A, FB4L, FB4M, and FB5H, rather than the dorsal FB DANs [148, 149]— may well perform similar functions. Interestingly, most tangential neurons that innervate the FB receive local input from columnar FB neurons —including subtypes of PFN, PFR, v$\Delta$ and h$\Delta$ neurons— near their presynaptic sites in the FB [29]. Any excitatory input that an FB DAN receives from a columnar neuron is likely to depolarize much of the tangential neuron's layered arbors, given the short electrotonic distances involved, but we speculate that input from columnar neurons may control local Ca, a mechanism hypothesized to enable presynaptic modulation of DA vesicle release in the mammalian striatum [150, 151]. Such a mechanism would ensure that any reinforcement signals that the FB DANs carry would be locally influenced by heading input, as required by our model. In our model, this heading-shaped neuromodulation would act on synapses between tangential motor-state neurons and goal neurons (SI Fig S7e-f). FB DANs as well as other tangential FB neurons send their outputs to other FB tangential neurons and to many columnar neurons, including the h$\Delta$, v$\Delta$ and FC neurons [29]. Several of these FB tangential neurons receive inputs in the LAL. For example, FB5A neurons receive synaptic inputs in the LAL, including from PFL2 and PFL3 neurons, making them ideal candidates to carry motor state signals (SI Fig S7f). Although different FB columnar neurons could act as goal neurons in different contexts, recent experiments suggest that FC2A neurons, which we hypothesized as candidates to carry a goal signal [87, 88], may be capable of influencing walking flies' actions in a manner consistent with such a function [84], while in other experiments, a class of h$\Delta$ neurons have been suggested to store goal headings [83].

**Implementing a policy of a fixed form: Phase shifts of CX output neurons.** A remarkable feature of several CX columnar neurons is the precision of their projection patterns within different CX structures. In particular, most CX columnar neurons that project from the PB to either the EB or the FB show precise phase shifts between their localized arbors in the different structures. These phase shifts are computed relative to the projected position of the EPG bump in different structures. For example, PEN neurons whose arbors overlap with EPG neurons in the PB project to a location in the EB that is shifted by $45°$ relative to their input EPGs [137]. In the case of the PEN_a neurons, this phase shift has been proposed to allow self-motion-derived angular velocity input to shift the position of EPG population activity in the EB [58, 59]. Our model relies on the projection patterns of the PFL2 and PFL3 neurons, which show phase shifts of $180°$ and $90°$ respectively between their PB and FB arbors (SI Fig S7d,f; note also that PFL2 neurons project to both left and right LALs). Our proposal for how these phase shifts might enable action selection is based on ideas proposed in [29] and, in the case of the PFL3 neurons, also shares similarities with a model that was originally proposed for path integration in the sweat bee [80]. The same idea was used for models in previous versions of this manuscript [87, 88], and related ideas have been used in work from other laboratories [81, 83–85]. As shown in Fig 4e, the phase shifts automatically enable a multiplication of a phase-shifted version of the fly's current heading with its goal heading. For the PFL2 neurons, the $180°$ shift means that the product of this multiplication peaks when the fly is heading in exactly the opposite direction to the goal heading. If activity in the PFL2 neurons modulates drift rate within neurons that control fixation, as we propose, then the result of peak activity would be high drift rate that results in shorter fixation and transitions to saccades. Thus, PFL phase shifts provide a potential mechanism to directly induce actions that would steer the bump towards a goal heading, allowing learning to work in the lower-dimensional space of merely updating the goal weight vector, rather than the space of actions necessary to direct the fly to its goal. In addition to their direct projections onto descending neurons (DNs) [29, 36] that send motor commands to the thorax [152–154] (SI Fig S7f), the PFL2 and PFL3 neurons also converge onto LAL neurons that themselves project onto DNs [29]. These projection patterns justify our modeling assumptions regarding how heading-tethered PFL (action neuron) activity is converted into directional motor commands. Recent experimental data from walking flies [84, 85] further support our assumptions.

**A potential circuit mechanism to learn and travel in the direction of a true goal heading.** As we discussed above, our model ignores the distinction between HD and heading for the purposes of modeling the fly's behavior in our paradigm. However, we believe that this paradigm exploits a mechanism that the fly uses during dispersal and long-range navigation in more natural settings [49, 53]. The FB's circuits could provide the requisite mechanism for the fly to travel in a specific direction, that is, to learn and then progress towards a goal heading. In essence, to produce appropriate movements, PFL neurons would need to receive FB input that has already accounted for the effects of different head-body angles and perturbations such as wind input through appropriate vector computations

[29, 60].

For example, imagine a fly attempting to travel northeast on a windy day. At every moment in time, the fly's total translational velocity ('TV') vector will be the sum of two components: the influence of the wind and the fly's self-generated movement, which we assume is in the same direction as its head direction (i.e., the fly's head direction matches its body direction). That is, $\mathrm{TV}(t) = \mathrm{Wind}(t) + \mathrm{Fly}(t)$. In this case, if the wind were blowing the fly east, the fly would have to fly north with the same velocity as $\mathrm{Wind}(t)$ to maintain a northeast heading. Here we assume that $\mathrm{Fly}(t)$ is the HD bump scaled by the fly's forward flight velocity, and that $\mathrm{TV}(t)$ is the fly's total translational velocity vector, carried by h∆B neurons [60, 139]. With these two signals, the fly could compute the allocentic wind direction (i.e., $\mathrm{Wind}(t) = \mathrm{TV}(t) - \mathrm{Fly}(t)$) (but see [83, 138]). Next, to compute the desired, or 'goal' (G), head direction, the fly would have to subtract the instantaneous wind direction from the goal translation vector (i.e., $\mathrm{HD}_G(t) = \mathrm{TV}_G - \mathrm{Wind}(t)$). Here we assume that $\mathrm{TV}_G$ is a vector that is stored in the FB that encodes the fly's desired travel direction. Once the desired head direction ($\mathrm{HD}_G(t)$) has been computed, the PFL2/3 neurons could generate goal-directed motor commands as described above. The complexity of this computation arises because the PFL2/3 neurons are thought to inherit the fly's HD in the PB, which would prevent them from directly comparing the fly's instantaneous TV ($\mathrm{TV}(t)$) to its goal TV ($\mathrm{TV}_G$). Instead, to accurately account for the influence of the wind, the PFL2/3 neurons would have to receive the desired HD vector ($\mathrm{HD}_G(t)$) as input in the FB. In this way, the PFL2/3 neurons could compare the fly's current HD to its goal HD. This conceptual model demonstrates that the fly CX could, in principle, account for external perturbations when selecting appropriate actions. Alternatively, instead of storing a desired travel direction, the fly could store a desired HD. Doing so could allow the fly to set an approximately accurate course under some circumstances. For example, if the wind consistently blew the fly east, maintaining a north head direction would yield an overall northeast travel, on average, but this mechanisms would not allow the fly to account for external perturbations like the wind. Future physiological recordings during behavior are required to assess which of these two classes of goal vectors the fly may be using and in which contexts.

#### Reinforcement learning framework

In this study, we used a reinforcement learning framework to explore how the fly's behavioral modes, namely fixations and saccades, should be structured as a function of the fly's heading and updated based on the fly's experience. We first segmented behavior into these two modes, and used the variability in these modes to inform the structure and adaptable control parameters of a behavioral policy, as described in *Behavioral Analysis Methods: Inferring the structure of a behavioral policy*. Here, we build an agent that uses this policy to structure its behavior as a function of its orientation within its visual surroundings. We then used reinforcement learning to train the control parameters of this policy.

In the main text, we considered the objective of maintaining a goal heading. We used this objective to explore how behavior might have been structured over evolutionary timescales, and then compared the results of this learning to naive fly behavior. As shown in Fig 3d, this objective gives rise to control parameters that are sinusoidally structured as a function of the fly's compass heading. We showed that these control parameters, when mediated by an unstable internal representation of heading and controlled with respect to an internal goal heading, produce behavior that qualitatively resembles fly behavior. With such a policy that is structured with respect to this internal goal heading, the fly need only shift the location of the goal heading to adapt to new surroundings. We thus refer to this policy as a "fixed-form" policy that specifies a set of actions that are tethered to the fly's internal representation of heading relative to an internal goal heading. Below, we show how reinforcement learning can be used on shorter timescales to shift the location of the goal heading based on experience. Finally, we illustrate how a simplified circuit model can be used to implement this fixed-form policy, and how plasticity in two sets of weights within the circuit can be used to update the goal heading and most stable compass heading over time.

In what follows, we first outline the general form of the policy and the training algorithm. We then show how the control parameters of this policy can be learned using a policy gradient algorithm with function approximation; we use this approach to study how the control parameters should be structured as a function of heading to maintain a preference for a specific visual pattern or a specific heading. We then assume that the structure in these control parameters is built-in to the policy and maintained with respect to a single goal heading, and we show how a policy gradient algorithm can be used to update the location of this goal heading while otherwise maintaining the structure of the policy. Finally, we outline the circuit implementation of this fixed-form policy, and we show how Hebbian-like plasticity can be used to implement the learning process.

#### General architecture

**Policy.** We consider a general scenario in which the behavior of an agent (model fly) is governed by a stochastic policy $\pi(\dot{\theta}, \Delta t|\theta)$. This policy determines the probability of maintaining an average angular velocity $\dot{\theta}$ over a duration of time $\Delta t$ given an initial heading $\theta$, and thereby determines the probability of generating a change in heading $\Delta\theta = \dot{\theta}\Delta t$. We will use $\Delta t$ to denote the duration of a single action that is sampled from the policy; depending on the scenario, this can correspond to the entire duration of a saccade, the entire duration of a fixation, or the duration of a sampling event within a fixation. The policy is parameterized by a set of fixed parameters $\vec{\beta}$ that do not change over time, and a set of flexible parameters $\vec{\omega}$ that can be modified through experience. For notational simplicity, we will introduce these parameters as they become necessary. When writing a conditional distribution $P(A|B)$, we will explicitly denote the dependence on $\vec{\omega}$ when it exists (i.e., $P(A|B;\vec{\omega})$), and will assume implicit dependence on $\vec{\beta}$ when it exists.

We decompose behavior into two different behavioral modes: fixations ('F'), and saccades ('S'), such that the policy can be written as:

$$\begin{aligned}\pi(\dot{\theta}, \Delta t|\theta) &= P(\dot{\theta}, \Delta t|\theta, F)P(F|\theta) + P(\dot{\theta}, \Delta t|\theta, S)P(S|\theta) \\ &= P(\Delta t|\dot{\theta}, \theta, F)P(\dot{\theta}|\theta, F)P(F|\theta) + P(\Delta t|\dot{\theta}, \theta, S)P(\dot{\theta}|\theta, S)(1 - P(F|\theta))\end{aligned} \tag{1}$$

Here, we have used the constraint that $P(F|\theta) + P(S|\theta) = 1$, and we have incorporated the observation that the duration of each mode can be conditioned on the angular velocity (see *Behavioral Analysis Methods: Characterizing fixation properties* and *Behavioral Analysis Methods: Characterizing saccade properties*).

**Fixation policy.** We assume that fixations are generated in a timepoint-by-timepoint manner via a drift diffusion process with integrated signal $\xi$; when this signal crosses a fixed threshold, the fixation is terminated and a saccade is initiated. The probability of maintaining a fixation thus depends on $\xi$:

$$\pi(\dot{\theta}, \Delta t|\theta, \xi) = P(\Delta t|\dot{\theta}, \theta, F)P(\dot{\theta}|\theta, F)P(F|\theta, \xi) + P(\Delta t|\dot{\theta}, \theta, S)P(\dot{\theta}|\theta, S)(1 - P(F|\theta, \xi)) \tag{2}$$

We assume that the integrated signal $\xi$ is updated in time increments of $\delta t$, and we assume that there is no change in heading during this time increment. This allows us to define:

$$P(\Delta t|\dot{\theta}, \theta, F) = \delta(\Delta t - \delta t) \tag{3}$$

$$P(\dot{\theta}|\theta, F) = \delta(\dot{\theta}) \tag{4}$$

During fixations, the integrated signal $\xi$ is updated by an amount $\Delta\xi$ that is determined by the drift diffusion process. We model this process with a fixed spread $\eta_F^2$ and a heading-dependent drift rate $\nu_F(\theta;\vec{\omega})$ that is parameterized by the flexible parameters $\vec{\omega}$ :

$$P(\Delta\xi|\theta, \xi; \vec{\omega}) = \mathcal{N}(\nu_F(\theta;\vec{\omega})\delta t, \eta_F^2\delta t) \tag{5}$$

This update will terminate the fixation and result in a saccade if the net signal $\xi + \Delta\xi$ crosses a fixed threshold $a_F$. Note that this produces an inverse Gaussian distribution of fixation durations $\Delta T$, with average durations $a_F/\nu_F(\theta;\vec{\omega})$ (where here, we use $\Delta T$ to denote the duration of the entire fixational event, computed in increments of $\delta t$):

$$P\left(\Delta T|\dot{\theta}, \theta, F; \vec{\omega}\right) = \mathrm{IG}\left(\Delta T; a_F/\nu_F(\theta;\vec{\omega}), a_F^2/\eta_F^2\right) \tag{6}$$

This allows us to write the probability of fixating as:

$$\begin{aligned}P(F|\theta, \xi; \vec{\omega}) &= \int P(F|\Delta\xi, \theta, \xi)P(\Delta\xi|\theta, \xi)d\Delta\xi \\ &= \int^{a_F} P(\Delta\xi|\theta, \xi)d\Delta\xi \\ &= \frac{1}{2}\left(1 + \mathrm{erf}\left(\frac{a_F - (\xi + \nu_F(\theta;\vec{\omega})\delta t)}{\sqrt{2\eta_F^2\delta t}}\right)\right)\end{aligned} \tag{7}$$

In the presence of heat, one can include second, reflexive drift process that can short-circuit the termination of a fixation (see *Behavioral Analysis Methods: Characterizing fixation properties* for motivation). To illustrate how this could be carried out, we consider a process that is governed by an integrated signal $\xi_R$ that obeys the same dynamics as above, with the same spread $\eta_F^2$ but with a fixed drift rate $\nu_R$. A fixation is terminated whenever either $\xi$ or $\xi_R$ cross the fixed threshold $a_F$. The probability of fixating is thus determined by:

$$P(F|\theta, \xi, \xi_R; \vec{\gamma}) = \min\Big\{P(F|\theta, \xi; \vec{\gamma}), P(F|\theta, \xi_R)\Big\} \tag{8}$$

where

$$P(F|\theta, \xi_R) = \frac{1}{2}\left(1 + \mathrm{erf}\left(\frac{a_F - (\xi_R + \nu_R \delta t)\Theta(\mathrm{heat})}{\sqrt{2\eta_F^2 \delta t}}\right)\right) \tag{9}$$

Here, $\Theta(\mathrm{heat})$ is a heaviside function that takes a value of $1$ if there is a perceived heat, and $0$ otherwise. Note that in the absence of perceived heat, $P(F|\theta, \xi; \vec{\gamma})$ will always be less than $P(F|\theta, \xi_R)$, and will thus determine the probability of fixating through Eq (8).

**Saccade policy.** We assume that saccades are initiated in a ballistic manner following the termination of a fixation, such that taking a saccade results in an abrupt change in heading $\Delta\theta = \dot{\theta}\Delta t$ over a duration of time $\Delta t$.
We assume that the directionality of saccades is controlled via a heading-dependent directional bias $d_S(\theta; \vec{\omega})$ that is parameterized by the flexible parameters $\vec{\omega}$; $d_S(\theta; \vec{\omega})$ specifies the probability of initiating a rightward, or CW, saccade at heading $\theta$. We then assume that the angular speed of a saccade $|\dot{\theta}|$ is drawn from a lognormal distribution with parameters $\varphi_S, \sigma_S^2$. Together, this results in the following distribution over angular velocities $\dot{\theta}$:

$$P(\dot{\theta}|\theta, S; \vec{\omega}) = \left[d_S(\theta; \vec{\omega})\,\mathrm{sgn}(\dot{\theta}) + \left(\frac{1 - \mathrm{sgn}(\dot{\theta})}{2}\right)\right] \mathrm{logn}(|\dot{\theta}|; \varphi_S, \sigma_S) \tag{10}$$

We further assume that the duration of saccades, analogously to the duration of fixations, can be generated via a drift diffusion process. Here, we assume that the drift rate is not flexible, but depends on the angular speed of the saccade:

$$P(\Delta t|\dot{\theta}, \theta, S) = \mathrm{IG}(\Delta t; a_S/\nu_S(|\dot{\theta}|), a_S^2/\eta_S^2) \tag{11}$$

where $\nu_S(|\dot{\theta}|)$ is well-captured by a sigmoidal function of $|\dot{\theta}|$ (see *Behavioral Analysis Methods: Characterizing saccade properties*). We can now specify the full policy and its parameter dependence:

$$\pi(\dot{\theta}, \Delta t|\theta, \xi; \vec{\omega}) = P(\Delta t|\dot{\theta}, \theta, F)P(\dot{\theta}|\theta, F)P(F|\theta, \xi; \vec{\omega}) + P(\Delta t|\dot{\theta}, \theta, S)P(\dot{\theta}|\theta, S; \vec{\omega})(1 - P(F|\theta, \xi; \vec{\omega}))$$

$$\begin{aligned}
P(\Delta t|\dot{\theta}, \theta, F) &= \delta(\Delta t - \delta t) \\
P(\dot{\theta}|\theta, F) &= \delta(\dot{\theta}) \\
P(F|\theta, \xi; \vec{\omega}) &= \frac{1}{2}\left(1 + \mathrm{erf}\left(\frac{a_F - (\xi + \nu_F(\theta; \vec{\omega})\delta t)}{\sqrt{2\eta_F^2 \delta t}}\right)\right) \\
P(\Delta t|\dot{\theta}, \theta, S) &= \mathrm{IG}(\Delta t; a_S/\nu_S(|\dot{\theta}|), a_S^2/\eta_S^2) \\
P(\dot{\theta}|\theta, S; \vec{\omega}) &= \left[d_S(\theta; \vec{\omega})\,\mathrm{sgn}(\dot{\theta}) + \left(\frac{1 - \mathrm{sgn}(\dot{\theta})}{2}\right)\right] \mathrm{logn}(|\dot{\theta}|; \varphi_S, \sigma_S)
\end{aligned} \tag{12}$$

where $\vec{\omega}$ controls the heading dependence in both the duration of fixations (through the drift rate $\nu_F$) and the directional bias of saccades (through $d_S$). As noted above, the policy depends implicitly on a set of fixed parameters $\vec{\beta} = [\delta t, a_F, \eta_F, \varphi_S, \sigma_S, a_S, \eta_S]$ that controls inflexible aspects of behavior.

**Training.** We use an online policy-gradient method to iteratively update the flexible policy parameters $\vec{\omega}$ based on the agent's actions and on the outcome of these actions. In this way, the agent (i) samples an action $[\dot{\theta}, \Delta t]$ from its policy based on its current heading $\theta$, (ii) observes the outcome of this action (a change in heading $\Delta\theta = \dot{\theta}\Delta t$, and a heading-dependent sensory response $R(\theta + \dot{\theta}\Delta t)$), and (iii) updates the policy weights $\vec{\omega}$ to modify the probability of taking the same action from the same heading in the future (depending on whether that action led to a good or bad outcome). We use a policy gradient algorithm [20] to update these weights:

$$\begin{aligned}\Delta\vec{\omega} &= R(\theta+\dot{\theta}\Delta t)\nabla_{\vec{\omega}}\log\pi(\dot{\theta},\Delta t|\theta;\vec{\omega})\Big|_{\theta^*,\dot{\theta}^*,\Delta t^*} \\ &= R(\theta+\dot{\theta}\Delta t)\frac{\nabla_{\vec{\omega}}\pi(\dot{\theta},\Delta t|\theta;\vec{\omega})}{\pi(\dot{\theta},\Delta t|\theta;\vec{\omega})}\Big|_{\theta^*,\dot{\theta}^*,\Delta t^*}\end{aligned} \quad (13)$$

where $\theta^*$, $\dot{\theta}^*$, and $\Delta t^*$ denote specific values of the heading, angular velocity, and duration of an action, respectively. We assume that $R(\theta+\dot{\theta}\Delta t)$ is computed directly from changes in sensory experience, rather than from a comparison between sensory experience and expected value. Note that the sensory response effectively acts as the step size, or learning rate, in the update equation for $\vec{\omega}$. As a result, a strong sensory response can result in quick but coarse updates, whereas as weak sensory response will result in slower but finer updates. The policy gradients can then be computed as follows:

$$\begin{aligned}\nabla_{\vec{\omega}}\pi =& P(\Delta t|\dot{\theta},\theta,F)P(\dot{\theta}|\theta,F)\nabla_{\vec{\omega}}P(F|\theta,\xi;\vec{\omega}) + \\ & P(\Delta t|\dot{\theta},\theta,S)\Big[(1-P(F|\theta,\xi;\vec{\omega}))\nabla_{\vec{\omega}}P(\dot{\theta}|\theta,S;\vec{\omega}) - P(\dot{\theta}|\theta,S;\vec{\omega})\nabla_{\vec{\omega}}P(F|\theta,\xi;\vec{\omega})\Big] \\ =& \Big[P(\Delta t|\dot{\theta},\theta,F)P(\dot{\theta}|\theta,F) - P(\Delta t|\dot{\theta},\theta,S)P(\dot{\theta}|\theta,S;\vec{\omega})\Big]\nabla_{\vec{\omega}}P(F|\theta,\xi;\vec{\omega}) + \\ & \Big[P(\Delta t|\dot{\theta},\theta,S)(1-P(F|\theta,\xi;\vec{\omega}))\Big]\nabla_{\vec{\omega}}P(\dot{\theta}|\theta,S;\vec{\omega})\end{aligned} \quad (14)$$

We can use Eq. (12) to further simplify the gradients:

$$\nabla_{\vec{\omega}}P(F|\theta,\xi;\vec{\omega}) = -\delta t\mathcal{N}(a_F;\xi+\nu_F(\theta;\vec{\omega})\delta t,\eta_F^2\delta t)\nabla_{\vec{\omega}}\nu_F(\theta;\vec{\omega}) \quad (15)$$

$$\nabla_{\vec{\omega}}P(\dot{\theta}|\theta,S;\vec{\omega}) = \mathrm{sgn}(\dot{\theta})\,\mathrm{logn}(|\dot{\theta}|;\varphi_S,\sigma_S)\nabla_{\vec{\omega}}d_S(\theta;\vec{\omega}) \quad (16)$$

Further evaluating the policy gradients depends on the form of $\nabla_{\vec{\omega}}\nu_F(\theta;\vec{\omega})$ and $\nabla_{\vec{\omega}}d_S(\theta;\vec{\omega})$, which are determined by the specific implementations that we consider below.

**Implementation.** Here and in the main text, we consider different variants of this basic framework. The first variant, shown in Fig 3d, considers a “fully flexible” policy in which $\nu_F(\theta;\vec{\omega})$ and $d_S(\theta;\vec{\omega})$ can be modified in a heading-dependent manner via function approximation with a set of weights $\vec{\omega} = [\vec{\omega}_F, \vec{\omega}_S]$. Learning then acts to change the functional form of $\nu_F(\theta;\vec{\omega}_F)$ and $d_S(\theta;\vec{\omega}_S)$ by modifying $\vec{\omega}$. An extreme version of this policy, in which learning acts independently at each of a finely discretized set of headings, is schematized in Fig 3c-1. The second variant, schematized in Fig 3c-2 and detailed in this SI, considers a “fixed-form” policy in which the heading-dependence in both $\nu_F(\theta;\vec{\omega})$ and $d_S(\theta;\vec{\omega})$ is structured with respect to a “goal heading” $\theta_G$. In this case, the flexible parameters $\vec{\omega} = \theta_G$ specify the goal heading, and learning acts by shifting the goal heading while preserving the structured heading dependence in $\nu_F(\theta;\vec{\omega})$ and $d_S(\theta;\vec{\omega})$. The final variant, shown in Figs 4-6, considers a circuit-based implementation of this fixed-form policy that is informed by physiology and connectomic data (see *SI: Linking the Conceptual Model to Known Anatomy*).

In what follows, we outline the assumptions and training algorithms for these model variants. In each variant, we remove the temporal variability in the duration of saccades by assuming that $P(\Delta t|\dot{\theta},\theta,S) = \delta(\Delta t - t_S)$, where $t_S$ is a constant (we will assume $t_S = 300$ms, based on the analysis described in *Behavioral Analysis Methods: Characterizing saccade properties*; see Table 1).

#### Flexible policy for maintaining a behavioral preference.

**Policy.** We approximate the limited angular resolution of the heading representation via a set of radial basis functions $g_i(\theta)$ ($i = 1...n$) that mimic the anatomical tiling of compass neurons in the Ellipsoid Body. Unless otherwise specified, we used $n = 16$ von Mises basis functions that uniformly tiled the range $[0, 360]$, with concentration factor $\kappa = 8$. With this representation, heading dependence in the drift rate of fixations $\nu_F$ and the directional bias of saccades $d_S$ can be achieved by constructing different weighted combinations of these basis functions, with weights $\vec{\omega}_F$ controlling the heading dependence in $\nu_F$, and weights $\vec{\omega}_S$ controlling heading dependence in $d_S$:

$$
\begin{aligned}
\nu_F(\theta; \vec{\omega}_F) &= f(\vec{\omega}_F^T \vec{g}(\theta); k_F, f_{0F}, f_{MF}) \\
d_S(\theta; \vec{\omega}_S) &= f(\vec{\omega}_S^T \vec{g}(\theta); k_S, f_{0S}, f_{MS})
\end{aligned}
\tag{17}
$$

where $\vec{\omega}_a^T \vec{g}(\theta)$ ($a \in \{F, S\}$) is a weighted sum over basis functions evaluated at $\theta$. The sigmoidal function $f(x; k, f_0, f_M) = f_M/(1 + \exp(-kx)) - f_0$ enforces bounds on the drift rate of fixations and the probability of a rightward saccade, given a set of parameters $[k, f_0, f_M]$ that control the slope, minimum, and maximum values of the sigmoid. We chose the values of these parameters to bound the drift rate between $0.01$ and $1.01$, and to bound the probability of rightward saccades between $0$ and $1$.

As noted above, this policy depends implicitly on a set of fixed parameters $\vec{\beta}$, which now includes the additional parameters that specify this flexible model: $\vec{\beta} = [\delta t, a_F, \eta_F, \varphi_S, \sigma_S, a_S, \eta_S, n, \kappa, k_F, f_{0F}, f_{MF}, k_S, f_{0S}, f_{MS}]$. The values of these parameters are listed in Table 1.

**Training.** The policy gradients are given by:

$$
\begin{aligned}
\nabla_{\vec{\omega}} \nu_F(\theta; \vec{\omega}) &= f'(\vec{\omega}_F^T \vec{g}(\theta)) \vec{g}(\theta) \\
\nabla_{\vec{\omega}} d_S(\theta; \vec{\omega}) &= f'(\vec{\omega}_S^T \vec{g}(\theta)) \vec{g}(\theta)
\end{aligned}
\tag{18}
$$

where $f'(x) = df/dx$ is the derivative of the sigmoidal function $f$. Together with Eqs. (13), (14)-(16), these gradients can be used to compute the update to $\vec{\omega}_F$ and $\vec{\omega}_S$. Training thus acts to change the heading dependence in fixations and saccades by reweighting the basis functions through changes in $\vec{\omega}_F$ and $\vec{\omega}_S$, as detailed in Algorithms 1-2.

In the main text, we used this flexible policy to determine how the average duration of fixations and direction of saccades should be structured as a function of heading in order to (i) maintain a preference for a specific visual pattern $P_G$ (with orientation $\theta(P_G)$), or (ii) maintain a goal heading $\theta_G$. In the first case, we used $n = 16$ von Mises basis functions that uniformly tiled the range $[0, 180]$. The weights corresponding to these basis function specify the actions that agent takes with respect to a particular orientation of a visual pattern; thus, these weights specify the same actions for the two possible orientations of the visual scene that produce the same orientation of a given visual pattern. In the second case, we used $n = 16$ von Mises basis functions that uniformly tiled the range $[0, 360]$. We used the following sensory response function that decays linearly with angular distance from the preferred heading:

$$
R(\theta) = \begin{cases} \frac{\pi}{10}\left(\frac{1}{2} - \min(|\theta - \theta(P_G)|)\right) & \text{preference for goal pattern } P_G \\ \frac{\pi}{10}\left(\frac{1}{2} - |\theta - \theta_G|\right) & \text{preference for goal heading } \theta_G \end{cases}
\tag{19}
$$

where $\min(|\theta - \theta(P_G)|$ returns the distance to the nearest orientation of a preferred pattern.

**Algorithm 1:** Learn flexible policy parameters $\vec{\omega}$ via policy-gradient method

**input:** parametrized policy $\pi(\dot{\theta}, \Delta t|\theta, \xi; \vec{\omega})$
**define:** total simulation time $T_{tot}$; fixed policy parameters $\vec{\beta}$
**initialize:** policy parameters $\vec{\omega} \in \mathbb{R}^d$; integrator $\xi = 0$; time $t = 0$; heading $\theta \in [0, 360]$

**while** $t < T_{tot}$ **do**
  **sample action from policy**
  $\dot{\theta}, \Delta t, \Delta\xi \sim \pi(\cdot|\theta, \xi; \vec{\omega})$
  **observe sensory response**
  $r \leftarrow R(\theta + \dot{\theta}\Delta t)$
  **update policy parameters**
  $\vec{\omega} \leftarrow \vec{\omega} + r\nabla_{\vec{\omega}} \log \pi(\dot{\theta}, \Delta t|\theta, \xi; \vec{\omega})$
  **update heading, time, integrator**
  $\theta \leftarrow \theta + \dot{\theta}\Delta t$
  $t \leftarrow t + \Delta t$
  $\xi \leftarrow \xi + \Delta\xi$
**end while**
**return** $\vec{\omega}$

**Algorithm 2:** Sample action $\dot{\theta}, \Delta t$ from flexible policy $\pi(\dot{\theta}, \Delta t|\theta, \xi; \vec{\omega})$

**inputs:** heading $\theta$; integrator $\xi$; flexible policy parameters $\vec{\omega}$; fixed policy parameters $\vec{\beta}$; basis functions $\vec{g}(\theta)$

**get current drift rate, directional bias**
$\nu_F(\theta; \vec{\omega}_F) \leftarrow f(\vec{\omega}_F^T \vec{g}(\theta); k_F, f_{0,F}, f_{M,F})$
$d_S(\theta; \vec{\omega}_S) \leftarrow f(\vec{\omega}_S^T \vec{g}(\theta); k_S, f_{0,S}, f_{M,S})$
**integrate drift signal**
$\Delta\xi \sim \mathcal{N}(\nu_F(\theta; \vec{\omega}_F)\delta t, \eta_F^2 \delta t)$
**if** $\xi + \Delta\xi > a_F$ **then**
  **saccade**
  **if** $\text{rand}(\cdot) < d_S(\theta; \vec{\omega}_S)$ **then**
    $\dot{\theta} \sim +\text{logn}(\varphi_S, \sigma_S^2)$ (CW saccade)
  **else**
    $\dot{\theta} \sim -\text{logn}(\varphi_S, \sigma_S^2)$ (CCW saccade)
  **end if**
  $\Delta t \leftarrow t_S$
**else**
  **fixate**
  $\dot{\theta} \leftarrow 0$
  $\Delta t \leftarrow \delta t$
**end if**
**return** $\dot{\theta}$,$\Delta t, \Delta\xi$

#### Fixed-form policy tethered to a single goal heading.

In Fig 6, we showed how a circuit-based implementation of a fixed-form, goal-heading-dependent policy (derived via the learning algorithm described in the previous section), when tethered to an unstable internal representation of heading, could qualitatively capture the observed structure of fly behavior. Here, we illustrate a simplified form of this fixed-form policy, and we show how learning could act to shift the goal heading via a policy-gradient learning algorithm. In the following section, we show how this fixed-form policy could be implemented in a circuit model, and how learning could be implemented through Hebbian-like plasticity, rather than through the explicit computation of policy gradients.

**Policy.** In what follows, we will use $\theta_A$ to specify the angular orientation of the fly in arena coordinates, and we will distinguish this from the orientation $\theta_C$ of the compass heading bump. Before accounting for jumps in the compass bump, we assume that changes in the arena heading $\Delta\theta_A$ are accompanied by the same change in compass heading $\Delta\theta_C$, such that $\Delta\theta_A = \Delta\theta_C$ and $\dot{\theta}_A = \dot{\theta}_C = \dot{\theta}$ (in the following section, we will explicitly account for the fact that in the fly heading circuit, the compass heading and the arena heading move in opposite directions during saccades). Bump jumps, which arise from symmetries in the visual environment, will further alter the compass heading $\theta_C$ relative to the arena heading $\theta_A$.

The fixed-form policy specifies the heading dependence in the drift rate of fixations $\nu_F$, and the directional bias of saccades $d_S$, relative to a goal heading $\theta_G \in [0, 360)$. Based on the results shown in Fig 3d, and given the prevalence of sinusoidal signals in the central complex, we define these dependencies to have the following functional forms:

$$\begin{aligned} d_S(\theta_C; \theta_G) &= -\frac{G_S}{2}\sin(\theta_C - \theta_G) + B_S + \frac{1}{2} \\ \nu_F(\theta_C; \theta_G) &= \frac{G_F}{2}(1 - \cos(\theta_C - \theta_G)) + B_F \end{aligned} \tag{20}$$

where $B_S$ and $G_S$ control the baseline direction and heading-dependence of saccades, and $B_F$ and $G_F$ similarly control the baseline duration and heading dependence of fixations. As before, the average duration of fixations at any given heading $\theta_C$ is given by $a_F/\nu_F(\theta_C; \theta_G)$, where $a_F$ is the threshold of the drift diffusion process.

We showed in the main text that the heading representation is unstable in symmetric scenes (Fig 2), and that this manifests in bump jumps whose size reflects the degree of symmetry. We consider the two-fold symmetric scene used in the main text, for which a bump jump results in an angular change of $\Delta\theta_J = \pm 180°$. We further assume that the probability of a jump varies non-uniformly with $\theta_C$. We expect that the bump will have the lowest probability of jumping from locations through which the fly has frequently turned (as these are locations where the visual map from ring neurons onto compass neurons will have most quickly stabilized). Because the goal heading defines the angular location toward which the fly will turn, we define the probability of a jump $p_J(\theta_C; \theta_G)$ to be sinusoidal, with a value that depends on the angular distance from the goal heading:

$$p_J(\theta_C; \theta_G) = \frac{G_J}{2}(1 + \cos(\theta_C - \theta_G)) + B_J \tag{21}$$

where $B_J$ and $G_J$ control the baseline probability and heading-dependence of bump jumps. When $G_J$ is 0, the bump has the same probability of jumping at every heading, determined by the value of $B_J$. For $G_J$ larger than 0, the probability of a jump will vary nonuniformly with $\theta$, and the minimum probability will be determined by $B_J$. The jump probability determines whether or not the heading will jump by $\Delta\theta_J$:

$$\theta_C = \begin{cases} \theta_C & \text{with probability } 1 - p_J(\theta_C; \theta_G) \\ \theta_C + \Delta\theta_J & \text{with probability } p_J(\theta_C; \theta_G) \end{cases} \tag{22}$$

From the perspective of a given arena heading $\theta_A$, the effective drift rate and saccade probabilities are given by:

$$\begin{aligned} d_S^{\text{eff}}(\theta_A; \theta_G) &= (1 - p_J(\theta_A; \theta_G))d_S(\theta_A; \theta_G) + p_J(\theta_A; \theta_G)d_S(\theta_A + \Delta\theta_J; \theta_G) \\ \nu_F^{\text{eff}}(\theta_A; \theta_G) &= (1 - p_J(\theta_A; \theta_G))\nu_F(\theta_A; \theta_G) + p_J(\theta_A; \theta_G)\nu_F(\theta_A + \Delta\theta_J; \theta_G) \end{aligned} \tag{23}$$

As before, this policy depends implicitly on a set of fixed parameters $\vec{\beta}$, which now includes the additional parameters that specify the structure setpoint policy: $\vec{\beta} = [\delta t, a_F, \eta_F, \varphi_S, \sigma_S, a_S, \eta_S, G_S, G_F, G_J, B_S, B_F, B_J]$. The values of these parameters are listed in Table 1.

**Training.** The fixed-form policy can be trained using the same policy gradient algorithm discussed above (see Algorithms S7-S7). We briefly discuss a simple algorithm for implementing this. We assume that the gain parameters $\{G_S, G_F, G_J\}$ and baseline parameters $\{B_S, B_F, B_J\}$ remain fixed during training, but that the goal heading $\theta_G$ is updated during training and only after a saccade. We further assume that bump jumps occur just after saccades, and that the goal heading is updated only after the bump has had an opportunity to jump. Thus, a single update consists of the following sequence of steps: (i) sample the duration of a fixation from an inverse Gaussian distribution, (ii) sample the direction and size of saccade, (iii) execute the change in heading in both arena and bump coordinates, (iv) flip a biased coin to determine whether the bump will jump; if the bump jumps, shift the bump location by $180°$, and (v) update the location of the goal heading.

Because the goal heading is shifted only after a saccade (and not during the process of fixation), we can further simplify the process by directly sampling the fixation duration from an inverse Gaussian distribution; i.e., a fixation of total duration $\Delta t$ is sampled directly from $P(\Delta t|\dot{\theta}, \theta_C, F) = \mathrm{IG}(a_F/\nu_F(\theta_C;\theta_G), a_F^2/\eta_F^2)$, rather than being generated in a timepoint-by-timepoint manner. We then sample a saccade of angular velocity $\dot{\theta} \sim \pi_S(\dot{\theta}|\theta_C;\theta_G)$ and fixed duration $t_S$ (where $\pi_S(\dot{\theta}|\theta_C;\theta_G) = P(\dot{\theta}|\theta_C, S;\theta_G)$ is given in Eq. (12)). Finally, we update the location of the goal heading based on the gradient of the policy with respect to goal heading:

$$\Delta\theta_G(\dot{\theta}^*, \theta_A^*, \theta_C^*;\theta_G) = R(\theta_A + \dot{\theta}\Delta t_S)\frac{1}{\pi_S(\dot{\theta}|\theta_C;\theta_G)}\frac{\partial \pi_S(\dot{\theta}|\theta_C;\theta_G)}{\partial\theta_G}\bigg|_{\dot{\theta}^*, \theta_A^*, \theta_C^*} \tag{24}$$

where $\Delta\theta_G(\dot{\theta}^*, \theta_A^*, \theta_C^*;\theta_G)$ specifies the change in the location of the goal heading that is produced by initiating a saccade with velocity $\dot{\theta}^*$ (and fixed duration $t_S$) from an arena heading $\theta_A^*$ and compass heading $\theta_C^*$, given an initial goal heading of $\theta_G$. The gradient is given by:

$$\begin{aligned}\frac{\partial \pi_S}{\partial\theta_G} &= \frac{\partial}{\partial\theta_G}P(\dot{\theta}|\theta_C, S;\theta_G)\\ &= \mathrm{sgn}(\dot{\theta})\,\mathrm{logn}(|\dot{\theta}|;\varphi_S,\sigma_S)\frac{\partial}{\partial\theta_G}d_S(\theta_C;\theta_G)\\ &= \frac{G_S}{2}\mathrm{sgn}(\dot{\theta})\,\mathrm{logn}(|\dot{\theta}|;\varphi_S,\sigma_S)\cos(\theta_C - \theta_G)\end{aligned} \tag{25}$$

#### Circuit implementation of a fixed-form policy

[RL Framework: Circuit Implementation of a Fixed-Form Policy] In Figs 4-6, we developed a circuit-based implementation of the fixed-form policy discussed in the previous section, and we used Hebbian learning to update the parameters of that model (note that this differs from the policy-gradient algorithms discussed above). We constructed this circuit model from populations of so-called columnar neurons that tile $360°$ of angular space. In what follows, we parametrize this space by $\theta$, and we express all quantities as functions of $\theta$. See *SI: Linking the Conceptual Model to Known Anatomy* for a discussion of the relationships between this model and anatomical and functional observations.

**Policy.** In the EB, the current heading $\theta_C$ is represented by a bump of activity maintained by a population of columnar compass neurons. We approximate this bump of activity using a von Mises function whose amplitude is normalized to 1:

$$r_C^{EB}(\theta, \theta_C) = \exp(\kappa\cos(\theta - \theta_C))/2\pi I_0(\kappa) \tag{26}$$

When the fly saccades by an angle $\Delta\theta_A$ in the arena, we assume that this is perfectly captured by the heading circuit, such that the current heading shifts by an equal but opposite angle of $\Delta\theta_C = -\Delta\theta_A$.

Flies are known to exhibit variability in the offset between the orientation of the heading bump and the orientation of the visual scene; we assume here that this offset is zero. In visual scenes without repeating patterns, plasticity between ring neurons and compass neurons ensures that this offset remains stable over time by reinforcing a relationship between visual features in the scenes (as conveyed through the ring neuron receptive fields) and the location of the heading bump. However, in scenes with repeating patterns, ring neurons with a given receptive field will respond similarly when the fly is oriented toward different symmetric views of the same scene. This will result in the strengthening of different sets of ring-to-compass-neuron weights that correspond to the same visual patterns, and will lead to jumps in the heading representation (Fig 2d). We incorporate these dynamics through a set of heading-dependent compass weights $\vec{\omega}_C = \omega_C(\theta_A, \theta_B)$ that capture the net inhibition from each ring neuron whose receptive field is positioned at $\theta_A$ onto each compass neuron with preferred heading $\theta_B$, and thereby determine the relative stability of different headings that correspond to symmetric views of the visual scene. For the visual scenes considered here, there is a two-fold symmetry, such that the fly sees identical views of the same scene at the two orientations $\theta$ and $\theta + \Delta\theta_J$, where $\Delta\theta_J = 180°$. These identical views will both activate the same two subsets of rings neurons; we assume here the scene fully activates rings neurons whose receptive fields are positioned at $\theta$ and $\theta + \Delta\theta_J$, and partially activates ring neurons whose receptive fields are positioned at $\theta \pm \delta\theta$ and $\theta + \Delta\theta_J \pm \delta\theta$, where $\delta\theta$ is the spacing between preferred headings. We take the net weight profile summed across these active ring neurons, $\omega_{\text{net}}(\theta) = \sum_{\text{act}} \omega_C(\theta_{A,\text{act}}, \theta)$, and we compare it between the two symmetric compass headings, $\theta_C$ and $\theta_C + \Delta\theta_J$. The higher the net weight at the current compass heading, the stronger the inhibition from the ring neurons, and the more likely the bump will jump by $\Delta\theta_J$ to the symmetric compass heading. We define this jump probability to be:

$$p_J(\theta_C; \vec{\omega}_C) = \frac{\omega_{\text{net}}(\theta_C)}{\omega_{\text{net}}(\theta_C) + \omega_{\text{net}}(\theta_C + \Delta\theta_J)} \tag{27}$$

From the EB, the heading representation travels through the protocerebral bridge to the fan-shaped body (FB), where the profile of compass activity takes on a sinusoidal shape:

$$r_C^{FB}(\theta, \theta_C) = \frac{1}{2}\left(\cos(\theta - \theta_C) + 1\right) \tag{28}$$

We assume that information about the fly's current compass heading is combined with information about the goal heading in the FB and then used to drive premotor activity in the lateral accessory lobe (LAL). Specifically, we assume that the information about the goal heading is stored in a set of synaptic weights $\vec{\omega}_G = \omega_G(\theta)$ from tangential motor-state neurons onto columnar goal neurons. Here, we assume that the motor state neurons are active (with a constant activity of one, i.e. $r_M(\theta) = 1\ \forall_\theta$) whenever the fly is flying. Thus, the activity profile $r_G(\theta)$ of the goal neurons gives a direct readout of the goal weights:

$$\begin{aligned} r_G(\theta; \vec{\omega}_G) &= r_M(\theta)\omega_G(\theta) \\ &= \omega_G(\theta) \end{aligned} \tag{29}$$

As we will detail below, the set of weights $\vec{\omega}_G$ fully determines the properties of the goal heading.

Finally, we consider populations of output neurons that receive goal activity $r_G(\theta)$ and phase-shifted heading activity $r_C(\theta, \theta_C + \vartheta)$ as inputs, and whose summed output depends on the overlap between current and goal heading through a multiplicative operation:

$$r_O(\theta_C, \vartheta; \vec{\omega}_G) = \frac{\sum_\theta r_C^{FB}(\theta, \theta_C + \vartheta) r_G(\theta; \vec{\omega}_G)}{\sum_\theta r_C^{FB}(\theta, \theta_C + \vartheta) r_C^{FB}(\theta, \theta_C + \vartheta)} + B_O \tag{30}$$

where $\vartheta$ is a phase shift, and $B_O$ is a baseline shift (described below). The form of the output activity in Eq. 30 ensures that this output activity will be structured sinusoidally as a function of the fly's current compass heading $\theta_C$ relative to the circular mean of the goal weights. To see this, note that the numerator of Eq. 30 can be written as:

$$\begin{aligned} \text{num}[r_O(\theta_C, \vartheta; \vec{\omega}_G)] &= \sum_\theta \frac{1}{2}\left(1 + \cos(\theta - (\theta_C + \vartheta))\right)\omega_G(\theta) \\ &= \frac{1}{2}\sum_\theta \omega_G(\theta) + \frac{1}{2}\left(\frac{e^{-i(\theta_C+\vartheta)}}{2}\sum_\theta e^{i\theta}\omega_G(\theta) + \frac{e^{i(\theta_C+\vartheta)}}{2}\sum_\theta e^{-i\theta}\omega_G(\theta)\right) \\ &= \frac{|\omega_G|}{2} + \frac{r_G}{2}\left(\frac{e^{-i(\theta_C+\vartheta-\theta_G)} + e^{i(\theta_C+\vartheta-\theta_G)}}{2}\right) \\ &= \frac{|\omega_G|}{2} + \frac{r_G}{2}\cos(\theta_C + \vartheta - \theta_G) \end{aligned} \tag{31}$$

where $\sum_\theta e^{i\theta}\omega_G(\theta)d\theta \equiv r_G e^{i\theta_G}$ specifies the modulus $r_G$ and angle $\theta_G$ of the circular mean of the goal weights, and $|\omega_G| = \sum_\theta \omega_G(\theta)$ specifies the total strength of the goal weights. As we will next show, saccades will drive the compass heading toward $\theta_G$, and fixations will be maintained longer at $\theta_G$, and thus we define $\theta_G$ to be the goal heading. The denominator of Eq. 30 is fixed; we will denote this as $D = \sum_\theta r_C^{FB}(\theta, \theta_C + \vartheta) r_C^{FB}(\theta, \theta_C + \vartheta)$.

The dynamic range of the output activity determines how strongly the goal drives behavior; the larger the range, the bigger the differential between the behavior at versus away from the goal location. In such cases with a large differential, we describe the behavior as “highly structured”. As can be seen in Eq. 31, this range will be maximized when the modulus of the circular mean, $r_G$, is maximized. Because we constrain the heading and goal weights to lie in the range $[0, 1]$ (more on this below), the weight profile that maximizes $r_G$ is a square wave in which $N/2$ consecutive weights take a value of $0$, and the remaining $N/2$ consecutive weights take a value of $1$. Below, we describe a learning rule that will drive the weights toward a sinusoidal profile with a single peak, which approximates this square wave profile. Thus, within the constraints of this learning rule, the more strongly sinusoidal the weight profile, the more structured the behavior.

We construct three different output populations, denoted left (‘L’), fixation (‘F’), and right (‘R’), that differ in the phase shift of their heading-tuned inputs [29]:

$$\begin{aligned}
r_L(\theta_C; \vec{\omega}_G) &= r_O(\theta_C, \vartheta = +90; \vec{\omega}_G) = \frac{|\omega_G|}{2D} + \frac{r_G}{2D}\cos(\theta_C + 90 - \theta_G) \\
r_F(\theta_C; \vec{\omega}_G) &= r_O(\theta_C, \vartheta = 180; \vec{\omega}_G) = \frac{|\omega_G|}{2D} + \frac{r_G}{2D}\cos(\theta_C + 180 - \theta_G) + B_C \\
r_R(\theta_C; \vec{\omega}_G) &= r_O(\theta_C, \vartheta = -90; \vec{\omega}_G) = \frac{|\omega_G|}{2D} + \frac{r_G}{2D}\cos(\theta_C - 90 - \theta_G)
\end{aligned} \tag{32}$$

where we chose $B_L = B_R = 0$, $B_C = \nu_{\max} - |\omega_G|/D$, and $\nu_{\max} = 1.1$; as described below, this ensures that gain of fixations will increase with increasing $|\omega_G|$. We further assume, based on known projection patterns [29], that the populations of left and right output neurons project unilaterally to descending neurons that control leftward and rightward saccades, respectively, and that the population of center output neurons projects bilaterally to both sets of descending neurons. We assume that the activity of the right and left output neurons thus determines the average directionality of saccades for a given heading $\theta$:

$$\begin{aligned}
d_S(\theta_C; \vec{\omega}_G) &= \frac{1}{2}(1 + r_R(\theta_C; \vec{\omega}_G) - r_L(\theta_C; \vec{\omega}_G)) \\
&= \frac{1}{2} + \frac{r_G}{D}\cos(\theta_C - 90 - \theta_G)
\end{aligned} \tag{33}$$

Note that this function will be largest, and will thus drive the highest probability of clockwise saccades (and counterclockwise bump rotations), when the heading is $90°$ to the right of the goal heading.

We similarly assume that the activity of the center output neurons determines the drift rate (and thereby the average duration) of fixations:

$$\begin{aligned}
\nu_F(\theta_C; \vec{\omega}_G) &= r_F(\theta_C; \vec{\omega}_G) \\
&= \nu_{\max} - \frac{|\omega_G|}{2D} + \frac{r_G}{2D}\cos(\theta_C + 180 - \theta_G)
\end{aligned} \tag{34}$$

$$\langle \Delta t_F(\theta_C; \vec{\omega}_G) \rangle = \frac{1}{\nu_F(\theta_C; \vec{\omega}_G)} \tag{35}$$

Thus, the baseline values of saccade and fixation properties are given by:

$$\begin{aligned}
\min[d_S] &= \frac{1}{2} - \frac{r_G}{D} \\
\min[\nu_F] &= \frac{2D\nu_{\max} - |\omega_G| - r_G}{2D} \\
\min[\langle \Delta t_F \rangle] &= \frac{2D}{2D\nu_{\max} - |\omega_G| + r_G}
\end{aligned} \tag{36}$$

and the gain of saccade and fixation properties are given by:

$$\begin{aligned}
\text{gain}[d_S] &= \frac{2r_G}{D} \\
\text{gain}[\nu_F] &= \frac{r_G}{D} \\
\text{gain}[\langle \Delta t_F \rangle] &= \frac{r_G/D}{\nu_{\max}^2 + |\omega_G|^2/(4D^2) - r_G^2/(4D^2) - \nu_{\max}|\omega_G|/D}
\end{aligned} \tag{37}$$

As can be seen from Eqs. 33-35, the modulus $r_G$ of the circular mean of the goal weights determines the gain of both the fixational drift rate and the saccade directionality; the larger $r_G$, the larger the directional bias when the heading bump is to the right or left of the goal heading, the longer the fixations at the goal heading, and the shorter the fixations away from the goal heading. In this way, the multiplicative operation performed by the output neuron populations (and specified by Eq. 31) guarantees that the fly's internal policy will remain structured as a function of the current heading relative to the goal heading, regardless of their specific values.

**Training.** We assume that there is plasticity in both $\vec{\omega}_C$ and $\vec{\omega}_G$ that is mediated by the activity of the heading bump in the EB and FB ($r_C^{EB}(\theta, \theta_C)$ and $r_C^{FB}(\theta, \theta_C)$, respectively):

$$\begin{aligned}
\Delta\omega_C(\theta, \theta_C; \vec{\omega}_C) &= \alpha_C \Delta_C v^2 \\
\Delta\omega_G(\theta, \theta_C; \vec{\omega}_G) &= \alpha_G \Delta_G
\end{aligned} \tag{38}$$

where

$$\begin{aligned}
\Delta_C &= \left[r_C^{EB}(\theta, \theta_C) - \omega_C(\theta)\right]_+ \Theta(1 - \omega_C) - \left[\omega_C(\theta) - r_C^{EB}(\theta, \theta_C)\right]_+ \Theta(\omega_C) \\
\Delta_G &= \begin{cases} +\left[r_C^{FB}(\theta, \theta_C) - \omega_G(\theta)\right]_+ \Theta(1 - \omega_G) - \left[\omega_G(\theta) - r_C^{FB}(\theta, \theta_C)\right]_+ \Theta(\omega_G) & R(\theta_A) > 0 \\ -\left[r_C^{FB}(\theta, \theta_C) - \omega_G(\theta)\right]_+ \Theta(\omega_G) + \left[\omega_G(\theta) - r_C^{FB}(\theta, \theta_C)\right]_+ \Theta(1 - \omega_G) & R(\theta_A) < 0 \\ 0 & R(\theta_A) = 0 \end{cases}
\end{aligned} \tag{39}$$

Here, $[\cdot]_+$ denotes rectification, and $\Theta(\cdot)$ is the heaviside function.

The first of these plasticity rules is similar to that used in [31] in that the change in weights is proportional to the coactivity between ring neurons (whose activity here is implicilty conveyed through the compass weights) and compass neurons (whose activity here is assumed to have a fixed profile $r_C^{EB}(\theta, \theta_C)$), and is modulated by fly's velocity $v$. In simulations, we use the size of the saccade, $\Delta\theta_S$, as a proxy for this velocity. We assume that the compass weights are only updated only during saccades, and that the goal weights are updated only during fixations. In practice, we partition saccades into angular increments $\delta\theta$ (i.e., based on the spacing between preferred headings), and we partition fixations into time increments of $100$ms. We then iteratively update weights at each angle/time increment. The second plasticity rule differs from the first in that it additionally incorporates the valence, $R(\theta_A)$, of the current arena heading $\theta_A$. We assume that this valence is carried by tangential neuromodulatory neurons that innervate the FB and themselves receive input from heading-tuned neurons [29]. We assume this valence takes the following form:

$$R(\theta_A) = \begin{cases} +1 & \theta_A \in \text{safe} \\ -1 & \theta_A \in \text{danger} \end{cases} \tag{40}$$

Algorithms 3-4 detail how this circuit model is implemented and updated through training.

**Algorithm 3:** Learn heading and goal weights, $\omega_C(\theta)$ and $\omega_G(\theta)$

**input:** parameterized policy $\pi(\Delta\theta, \Delta t|\theta; \vec{\omega}_C, \vec{\omega}_G)$
**define:** total simulation time $T_{tot}$, fixed policy parameters $\vec{\beta}$, learning rates $\alpha_C$, $\alpha_G$, heading resolution $\delta\theta$
**initialize:** weights $\vec{\omega}_C$ and $\vec{\omega}_G$; compass heading $\theta_C \in [0, 360)$; arena heading $\theta_A = \theta_C$; time $t = 0$, $\Delta t = 0$;

**while** $t < T_{tot}$ **do**
  **sample action from policy**
  $[\Delta\theta_S, \Delta\theta_F], [\Delta t_S, \Delta t_F] \sim \pi(\cdot|\theta_C; \vec{\omega}_C, \vec{\omega}_G)$
  **update arena heading, compass heading, and compass weights during a saccade**
  $t \leftarrow t + \Delta t_S$
  $\Delta\theta = 0$
  **while** $\Delta\theta < \Delta\theta_S$ **do**
    $\omega_C(\theta) \leftarrow \omega_C(\theta) + \alpha_C \Delta_C \Delta\theta_S^2$
    $\theta_C \leftarrow \theta_C + \delta\theta$
    $\theta_A \leftarrow \theta_A - \delta\theta$
    $\Delta\theta = \Delta\theta + \delta\theta$
  **end while**
  **determine whether bump will jump**
  **if** $\mathrm{rand}(\cdot) < p_{\mathrm{jump}}(\theta_C; \vec{\omega}_C)$ **then**
    $\theta_C \leftarrow \theta_C + \Delta\theta_C$
  **end if**
  **observe sensory response**
  $r \leftarrow R(\theta_A)$
  **update goal weights during fixation**
  **while** $\Delta t < \Delta t_F$ **do**
    $\omega_G(\theta) \leftarrow \omega_G(\theta) + \alpha_G \Delta_G$
    $\Delta t \leftarrow \Delta t + 0.1$
  **end while**
  $\Delta t = 0$
**end while**
**return** $\vec{\omega}_C, \vec{\omega}_G$

**Algorithm 4:** Sample action sequence $[\Delta\theta_S, \Delta\theta_F]$, $[\Delta t_S, \Delta t_F]$ from setpoint policy $\pi(\Delta\theta, \Delta t|\theta; \vec{\omega}_C, \vec{\omega}_G)$

**inputs:** heading $\theta_C$; weights $\vec{\omega}_C$, $\vec{\omega}_G$

**get current drift rate and directional bias**
$\nu_F(\theta_C; \vec{\omega}_G) \leftarrow r_F(\theta_C; \vec{\omega}_G) + B_F$
$d_S(\theta_C; \vec{\omega}_G) \leftarrow (1 + r_R(\theta_C; \vec{\omega}_G) - r_L(\theta_C, \vec{\omega}_G))/2$
**saccade**
**if** $\mathrm{rand}(\cdot) < d_S(\theta_C; \vec{\omega}_G)$ **then**
  $\Delta\theta_S \sim +\mathrm{logn}(\varphi_S, \sigma_S^2)$ (CW)
**else**
  $\Delta\theta_S \sim -\mathrm{logn}(\varphi_S, \sigma_S^2)$ (CCW)
**end if**
$\Delta t_S \leftarrow t_S$
**fixate**
$\Delta\theta_F \leftarrow 0$
$\Delta t_F \leftarrow 1/\nu_F(\theta_C; \vec{\omega}_G)$
**return** $[\Delta\theta_S, \Delta\theta_F]$, $[\Delta t_S, \Delta t_F]$

| | | | |
|---|---|---|---|
| **General policy parameters** | | | |
| $T_{tot}$ | 240 | total simulation time in sec | duration of two trials |
| $t_S$ | 320 | duration of saccade in msec | median duration in data |
| $\delta t$ | 0.001 | timescale of drift diffusion process in sec | sampling rate of behavioral data |
| $\eta_F$ | 1 | spread of drift diffusion process | estimated from distribution |
| $a_F$ | 0.79 | threshold for drift diffusion process | of fixation durations |
| $\varphi_S$ | 3.89 | parameters of lognormal distribution | estimated from distribution |
| $\sigma_S$ | 0.54 | over saccade sizes (in deg) | of saccade sizes |
| **Parameters for flexible policy** | | | |
| $k_F$ | 1 | sensitivity of drift rate | chosen for illustration |
| $f_{0,F}$ | -0.01 | sets minimum and maximum scale | constrains avg fixation duration |
| $f_{M,F}$ | 10 | of drift rate | between 100ms and 120s |
| $k_S$ | 1 | sensitivity of saccade probability | chosen for illustration |
| $f_{0,S}$ | -0.01 | sets minimum and maximum scale | constrains saccade probability |
| $f_{M,S}$ | 0.98 | of saccade probability | between 0.01 and 0.99 |
| $n$ | 16 | number of von Mises functions | matched to EB tiling |
| $\kappa$ | 8 | concentration of von Mises functions | |
| **Parameters for setpoint policy** | | | |
| $G_S$ | 0.9 | gain of saccades (controls heading-dependence) | chosen for illustration |
| $B_S$ | 0 | baseline direction of saccades | |
| $G_F$ | 0.8 | gain of fixations (controls heading-dependence) | chosen for illustration |
| $B_F$ | 0.05 | baseline duration of fixations | |
| $G_J$ | 0.7 | gain of bump jump (controls heading-dependence) | chosen for illustration |
| $B_J$ | 0.05 | baseline probability of jump | |
| $\Delta\theta_J$ | 180 | size of bump jump in deg | matched to data |
| **Parameters for circuit implementation of setpoint policy** | | | |
| $N$ | 32 | discretization of heading space | chosen for illustration |
| $\kappa$ | $\pi$ | concentration of von Mises functions | chosen for illustration |
| $\alpha_C$ | 0.01 | learning rate of compass weights | chosen for illustration |
| $\alpha_G$ | 0.001 | learning rate of goal weights | |

**Table 1:** Parameter values used in RL and circuit models.
